# Large-scale DNA-based phenotypic recording and deep learning enable highly accurate sequence-function mapping

**DOI:** 10.1101/2020.01.23.915405

**Authors:** Simon Höllerer, Laetitia Papaxanthos, Anja Cathrin Gumpinger, Katrin Fischer, Christian Beisel, Karsten Borgwardt, Yaakov Benenson, Markus Jeschek

**Affiliations:** Department of Biosystems Science and Engineering, ETH Zurich, Basel, Switzerland; Swiss Institute of Bioinformatics, Switzerland

**Author notes:** Equal contribution (Simon Höllerer, Laetitia Papaxanthos).

## Abstract

Predicting quantitative effects of gene regulatory elements (GREs) on gene expression is a longstanding challenge in biology. Machine learning models for gene expression prediction may be able to address this challenge, but they require experimental datasets that link large numbers of GREs to their quantitative effect. However, current methods to generate such datasets experimentally are either restricted to specific applications or limited by their technical complexity and error-proneness. Here we introduce DNA-based phenotypic recording as a widely applicable and practical approach to generate very large datasets linking GREs to quantitative functional readouts of high precision, temporal resolution, and dynamic range, solely relying on sequencing. This is enabled by a novel DNA architecture comprising a site-specific recombinase, a GRE that controls recombinase expression, and a DNA substrate modifiable by the recombinase. Both GRE sequence and substrate state can be determined in a single sequencing read, and the frequency of modified substrates amongst constructs harbouring the same GRE is a quantitative, internally normalized readout of this GRE’s effect on recombinase expression. Using next-generation sequencing, the quantitative expression effect of extremely large GRE sets can be assessed in parallel. As a proof of principle, we apply this approach to record translation kinetics of more than 300,000 bacterial ribosome binding sites (RBSs), collecting over 2.7 million sequence-function pairs in a single experiment. Further, we generalize from these large-scale datasets by a novel deep learning approach that combines ensembling and uncertainty modelling to predict the function of untested RBSs with high accuracy, substantially outperforming state-of-the-art methods. The combination of DNA-based phenotypic recording and deep learning represents a major advance in our ability to predict quantitative function from genetic sequence.

## Introduction

Recent progress in DNA sequencing and synthesis has facilitated reading and (re-)writing of the genetic makeup of biological systems on a massive scale^1, 2^. Despite this progress, the relationship between a genetic sequence and its functional properties is poorly understood, and thus the question “what to write” remains largely unanswered^3, 4^. Since the number of possible sequences scales exponentially with their length, the theoretical sequence space cannot be exhaustively explored by experiments, even for small GREs^5–7^. Therefore, innovative high-throughput (HTP) approaches are required that allow to collect a quantitative functional readout for large numbers of genetic sequences^7, 8^. At the same time, novel methods are required that identify statistical patterns and dependencies in the resulting datasets to generate models that accurately predict the properties of untested sequences. Deep learning maximizes the benefit of data collection at large scale owing to its ability to capture complex, nonlinear dependencies and to its computational scalability^9^, which led to several successful applications in computational biology, from genomics to proteomics^10–15^. These methods promise to be able to model sequence-function dependencies with minimal prior assumptions, provided that large experimental training datasets that link sequence to quantitative measure of function^16, 17^ are available.

While next-generation sequencing (NGS) allows obtaining sequence information at extremely large scale, our ability to assign a quantitative functional readout to each sequence has not kept pace. In previous efforts to alleviate this experimental bottleneck^3, 18–21^, the functional readout is performed in a separate technical step, and retroactively mapped back to the corresponding sequence by statistical inference. This introduces errors and limits data quality^21, 22^ impairing prediction accuracy. In other cases, quantitative readouts, such as ribosome loading^23^, DNA methylation status^7, 24^, or enrichment by growth selection^25^, are technically challenging and case-specific. RNA sequencing techniques avoid some of these limitations but are restricted to transcriptional effects and can be greatly biased due to variability in reverse transcription, barcode-induced bias, and DNA amplification efficiencies^26, 27^. Therefore, the need for widely applicable, technically simple and yet accurate approaches to ascribe functional (or phenotypic) readouts to genetic sequences at high throughput persists.

Here, we introduce for the first time a method that relies on DNA-based phenotypic recording and addresses the limitations enumerated above. Its core innovation is a three-component genetic architecture that combines on the same DNA molecule a gene for a site-specific DNA recombinase, a GRE controlling the expression of this recombinase, and the recombinase substrate. Thus, a functional link between GRE and the state of the recombinase substrate is established, and the latter serves as a stable and heritable record of the GRE’s effect on gene expression. Each DNA molecule embodying this architecture contains information about both the GRE sequence and a measure of its function (i.e. the modified vs. unmodified state of the substrate), both of which can be read in a single sequencing read and thus unambiguously linked. Relying on this principle, large libraries of GREs of interest can be assessed solely relying on NGS rendering separate functional experimentation obsolete. This greatly simplifies experimental procedures, enables measurements at high kinetic resolution, eliminates bias, and avoids the need for inference of the functional readout. Further, while any single DNA molecule will generate a binary functional record, the resolution of the readout can be arbitrarily increased by sequencing multiple DNA copies to obtain a frequency of modified substrates for each individual candidate GRE.

We use this approach termed uASPIre <u>(u</u>ltradeep <u>A</u>cquisition of <u>S</u>equence-<u>P</u>henotype Inter<u>re</u>lations) to record more than 2.7 million sequence-function data points in a single experiment to kinetically measure translation from 303,503 RBS variants in *Escherichia coli*. Further, we exploit the resulting high-resolution kinetic data to train a residual convolutional neural network ensemble (SAPIENs: <u>S</u>equence-<u>A</u>ctivity <u>P</u>rediction In <u>E</u>nsemble of <u>N</u>etwork<u>s</u>) that quantitatively predicts translational behaviour and quantifies reliably the uncertainty of the predicted RBS activities. Crucially, the combination of uASPIre and SAPIENs leads to hitherto unmatched prediction accuracy for RBSs as reflected by a coefficient of determination R^2^ of 0.927 and mean absolute error MAE of 0.039, notably without requirement for prior mechanistic knowledge about the underlying process of translation.

## Results

### The uASPIre Principle

In its broadest sense, uASPIre relies on a three-component DNA architecture comprising a genetic sequence to be investigated (“diversifier”), the gene of a DNA-modifying enzyme (“modifier”), and the cognate DNA substrate of this enzyme (“discriminator”), all located on the same DNA molecule (Fig. 1a). The modifier can alter the sequence of the discriminator, which can thus appear in at least two discrete “states” corresponding to modified and unmodified DNA substrate, respectively. The diversifier is placed in a genetic context that allows it to either directly or indirectly affect the activity of the modifier through gene regulation (e.g. if the diversifier is a GRE) or otherwise. The more a diversifier activates (or inactivates) the modifier, the higher (or lower) is the likelihood of discriminator modification. Hence, the discriminator serves as a DNA record of functional information about the diversifier’s activity, a concept we term DNA-based phenotypic recording. Consequently, both sequence and function of the diversifier can be determined concomitantly in a single sequencing read. While binary on the level of a single DNA molecule, the fraction of modified discriminators amongst all DNA copies that share the same diversifier constitutes a direct, quantitative, and internally-normalized readout of diversifier function that can be precisely tracked over time. If only a single diversifier variant is present per cell or compartment, an unambiguous link between a diversifier’s sequence and its function is stably and heritably established and maintained on the level of single DNA molecules. Therefore, NGS can be used to assess extremely large diversifier libraries. Crucially, dynamic range and resolution of the functional readout (i.e. fraction of modified discriminators) as well as the overall throughput of the method can be arbitrarily increased by adapting sequencing depth (i.e. the number of reads obtained per diversifier variant) and the number of total reads. Herein, we establish of a proof of concept for the described approach of DNA-based phenotypic recording by demonstrating the assessment of large numbers of RBSs as exemplary diversifiers.

**Fig. 1.**
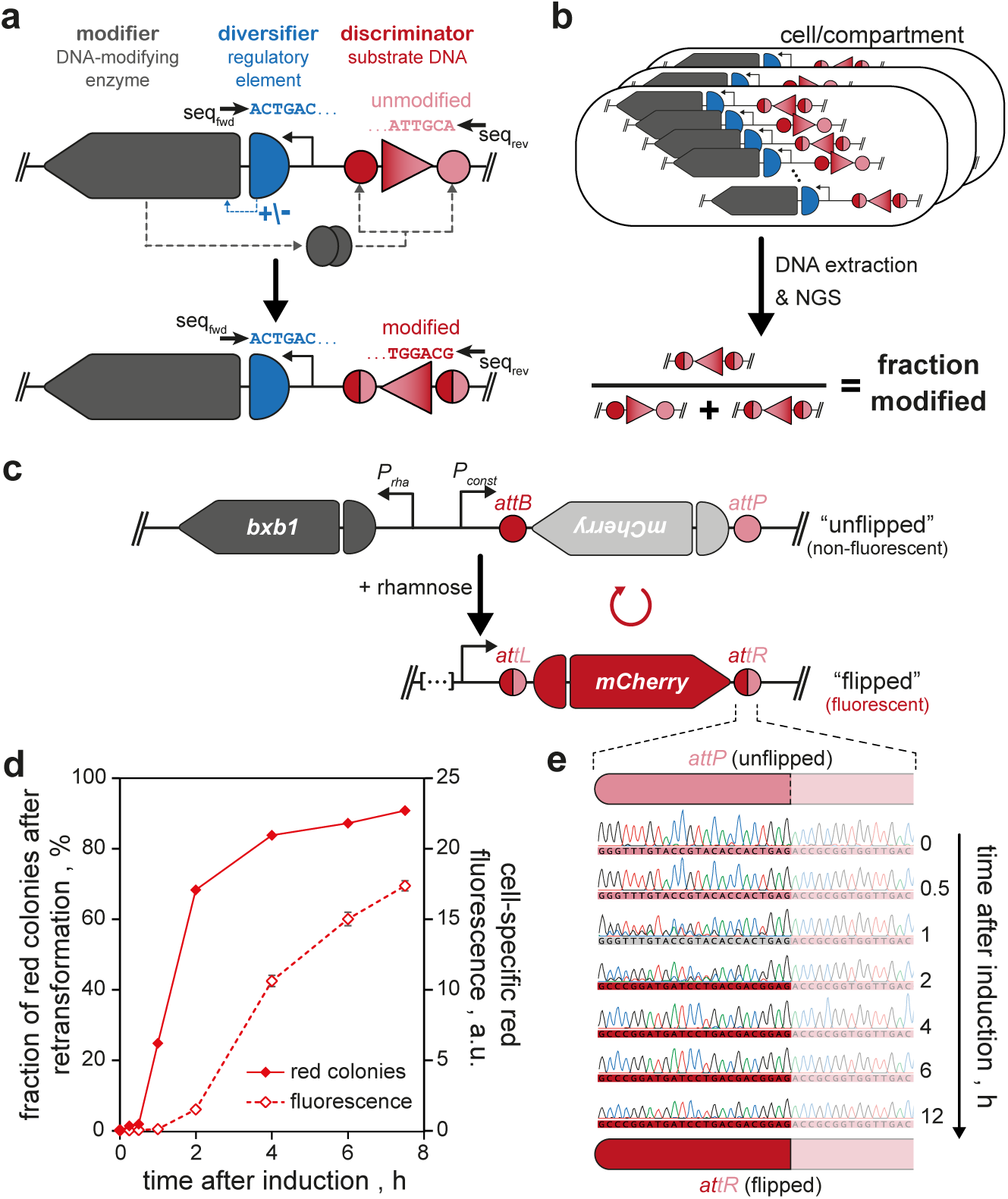
Basic principle of uASPIre and prototype DNA architecture. **a**, Generalized genetic architecture underlying the uASPIre approach. A genetic sequence of interest (diversifier; e.g. a GRE) controls, either positively (+) or negatively (-), the activity of a DNA-modifying enzyme (modifier), which can modify its cognate substrate DNA (discriminator). If placed on the same DNA molecule, diversifier sequence and discriminator state can be both determined by sequencing, for instance by NGS using forward (seq_fwd_) and reverse (seq_rev_) primers. **b**, Readout of the uASPIre method. Under monoclonal conditions (i.e., only one diversifier variant per compartment/cell), sequencing of multiple DNA copies that share the same diversifier allows to determine the fraction of modified discriminators, which can be used as a continuous, normalized readout for diversifier function. **c**, Prototype plasmid employing recombinase Bxb1 as a modifier controlled by the rhamnose-inducible promoter *P_rha_*. Bxb1 inverts an mCherry CDS into the correct orientation relative to a constitutive promoter *P_const_*, thus activating *mCherry* expression. *attB*/*P* and *attL*/*R*: Bxb1 attachment sites before and after recombination. **d**/**e**, Kinetics of Bxb1-mediated discriminator modification in shake flask cultivations of *E. coli*. Recombination is detected by direct fluorescent measurement and counting of red colonies after plasmid extraction and retransformation (**d**) and bulk Sanger sequencing of the discriminator sequence (**e**). Data points in (**d**) are means of three technical replicates with standard deviation (error bars).

### uASPIre for GRE Assessment

While several DNA-modifying enzymes are available, we chose site-specific recombinases as a modifier for practical realization of uASPIre in this study. Recombinases have been used to record cellular events, for example by inducing a reporter gene in certain cell types^28, 29^, or to discover cell- and tissue-specific promoters^30^. Diversification of a recombinase coding sequence (CDS) was used to discover variants with altered specificity^31^. We selected the well-characterized integrase from bacteriophage Bxb1 (*bxb1*/Bxb1) because it is self-sufficient in catalysing irreversible recombination, active in Pro- and Eukarya, and highly specific due to its long attachment sites *attB* and *attP* (50 bp and 53 bp) making off-target effects unlikely^32–34^. The two-state discriminator used in conjunction with Bxb1 is a short DNA sequence flanked by *attB* and *attP* sites in an orientation that leads to irreversible DNA sequence inversion (referred to as “flipping” hereafter) by the recombinase.

We anticipated two critical prerequisites for the use of recombinases such as Bxb1 for uASPIre. First, recombinase expression must be tightly regulated to ensure precise control over discriminator modification. Second, modification of discriminators should occur within a practical time window of a few hours that allows to make use of the full dynamic measurement range. To evaluate technical feasibility of these requirements, we constructed an *E. coli* prototype plasmid (pASPIre1) to track the activity of Bxb1 using fluorescence measurements as a proxy. In this construct, Bxb1-mediated recombination inverts an mCherry CDS into sense orientation relative to its promoter to convert a non-fluorescent discriminator state into a fluorescent one (Fig. 1c). The requirement for a tightly regulated expression of the recombinase prompted us to use the L-rhamnose-inducible promoter *P_rha_*^35^ to control *bxb1* transcription. We used pASPIre1 to assess Bxb1 recombination relying on bulk fluorescence measurements, counting of red colonies after retransformation of plasmid isolates from the culture (Fig. 1d), and by Sanger sequencing (Fig. 1e). These experiments collectively showed that Bxb1 can be tightly controlled by addition of rhamnose to the culture without substantial recombination occurring beforehand. Moreover, the reaction proceeds for several hours allowing to flexibly sample the dynamic measurement range of discriminator inversion.

Next, we adapted the prototype architecture (Fig. 1a) to enable large-scale assessment of GREs, specifically RBSs, directly by NGS (Fig. 2a). First, superfolder green fluorescent protein (sfGFP) was fused to the Bxb1 C-terminus for later recording of calibration curves (see below). This Bxb1-sfGFP fusion retained activity exhibiting similar reaction dynamics as the sfGFP-less variant (plasmid pASPIre2, Supplementary Fig. 1). Moreover, we replaced *mCherry* in the discriminator with 150 bp of non-coding DNA (plasmid pASPIre3, Fig. 2a, Supplementary Fig. 2) and constructed a rhamnose utilization-deficient strain to avoid consumption of inducer and ensure stable induction throughout the cultivation (*E. coli* TOP10Δ*rhaA*, Supplementary Fig. 3). Next, we used this system to characterize libraries of RBSs at high throughput relying on the uASPIre principle. As a part of the 5’-untranslated region (5’-UTR) of bacterial mRNAs, RBSs dictate the rate-limiting initiation of translation^36^. Because few mutations in this region can lead to orders-of-magnitude differences in protein expression, RBSs have become proven targets for optimization of cellular protein levels, in particular in multi-protein systems such as metabolic pathways^37, 38^. This trend has been largely fuelled by models that predict the relative strength of RBSs^39–42^ and tools for smart RBS library design^43, 44^. However, current models are insufficiently accurate to reliably allow accurate prediction and rational forward engineering^44–46^, mainly due to the fact that they are based on small datasets of experimental endpoint measurements (<10^3^ RBS variants), which do not cover a representative fraction of the vast number of possible RBSs and disregard the highly dynamic nature of translation initiation. We thus hypothesized that time-resolved activity data for much larger RBS populations could be used for the development of predictive models with greatly improved accuracy.

**Fig. 2.**
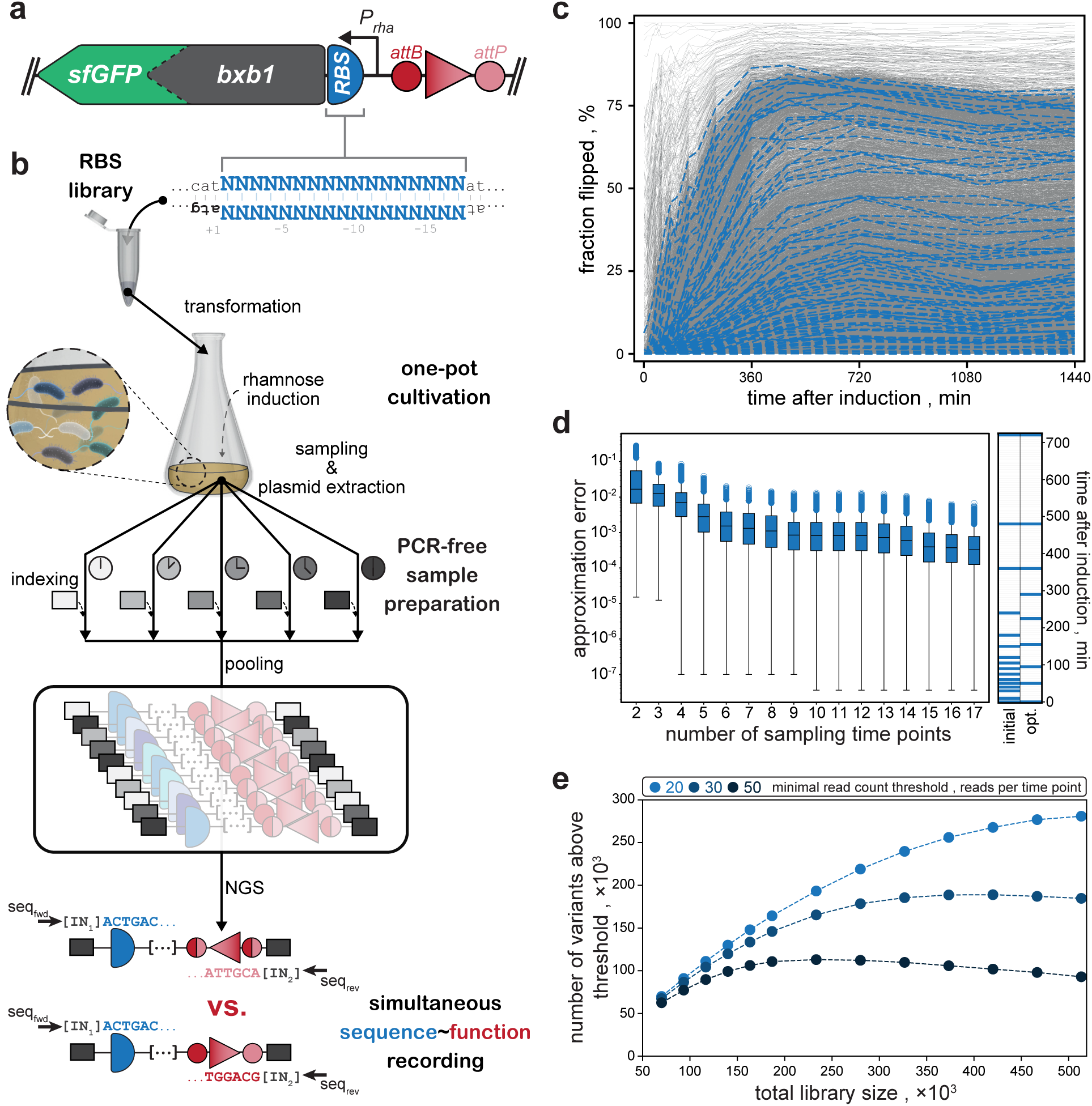
Establishment and throughput optimization of uASPIre for RBSs in *E. coli*. **a**, DNA architecture to facilitate simultaneous sequence-function assessment using NGS. As discriminator, a stretch of non-coding DNA flanked by *attB/P* sites is used. A Bxb1-sfGFP fusion is used as a modifier and controlled by different RBSs as target for diversification (plasmid pASPIre3). **b**, Experimental workflow for the uASPIre of RBSs. An RBS library with 17 randomized bases upstream of the the start codon (bold atg, shown upside down) of Bxb1 is used to transform *E. coli*. Monoclonal transformants each carry a different RBS variant (indicated by cells in different shades of blue). [IN_1/2_]: sample-specific indices. **c**, Kinetic behaviour of 10,427 RBSs (grey lines) recorded by uASPIre. The fraction of flipped discriminators is plotted over time after induction. Flipping profiles for 100 randomly selected RBSs are highlighted in blue for clarity. Data retrieved from a single Illumina NextSeq run (∼400 million paired-end reads, average of ∼18,700 reads per RBS). **d**, Optimization of sampling schedule. Left panel: Effect of reducing the number of sampling time points on the approximation error between reduced (2- to 17-time-point) and initial (18-time-point) flipping profiles for the RBSs from the library (Methods). Boxes contain interquartile range with median as centre line. Whiskers contain 1.5-fold interquartile range with outliers as circles. Right panel: Initial (18 time points) and optimized (nine time points) sampling schedule. **e**, Throughput optimization by adjustment of NGS loading. The effect of the total library size on throughput (i.e. number of variants above read-count threshold) of uASPIre for the optimized sampling schedule is shown for different read-count thresholds.

We established the following experimental workflow for the uASPIre of RBSs in *E. coli* (Fig. 2b, Methods). An RBS library with diversified 5’-UTR is used to transform *E. coli* and monoclonal transformants are co-cultivated. After induction with rhamnose, the culture is sampled at different time points and plasmid DNA is extracted, followed by agarose-gel purification of target DNA fragments spanning both RBS and discriminator. Next, duplex DNA-adapters containing time-sample-specific indices are ligated to the target fragments, samples are pooled, and the library pool is subjected to NGS. RBS sequence and discriminator state (here “unflipped” or “flipped”) are determined using paired-end reading. Finally, NGS raw data is processed (Methods) to obtain the dynamics of translation for each RBS, as reflected by the fraction of flipped discriminators among all sequencing reads obtained for this RBS (“fraction flipped” hereafter) over time (“flipping profiles” hereafter).

We used this workflow to analyse a library of ∼10,500 variants with 17 randomized bases (N_17_) in the 5’-UTR of the *bxb1-sfGFP* mRNA over 18 time points (Fig. 2c, Methods). Importantly, a single NGS run (Illumina NextSeq, ∼400 million reads) was used at an excessively high coverage (on average ∼18,700 reads per variant). While the library size for this proof of concept was chosen conservatively small, it pointed towards important features of our approach: (i) the achievable throughput is very high (here 187,686 sequence-function pairs distributed over 18 time points), and it could be significantly increased by optimizing read coverage and number of time samples; (ii) The RBS activity as represented by the flipping profiles is directly and quantitatively assessed at high resolution for the functional readout (fraction flipped) and with high technical reproducibility (Supplementary Fig. 4); (iii) Measurements can be performed in short intervals down to a few minutes or less facilitating acquisition of precise kinetic data. Notably, this is required to properly resolve the diversity of translation rates in the variant library, since end-point measurement would either lead to underestimation of strong variants (“late sampling”) or low resolution for weak RBSs (“early sampling”); (iv) No PCR-amplification is required during the entire workflow, which we have identified as a major source for the introduction of non-systematic bias (Supplementary Fig. 5) that is absent for the PCR-free workflow (Supplementary Fig. 6).

Using this proof-of-concept dataset, we optimized critical experimental parameters to increase the throughput of uASPIre. First, we evaluated the effect of reducing the number of sampling time points on the ability to reconstruct the full 18-time-point flipping profiles from the proof-of-concept experiment (Methods). This analysis indicated that the number of sampling time points can be significantly reduced without major deviation from the full flipping profiles (Fig. 2d) to save NGS read capacity and thus increase the overall throughput. We selected an optimized schedule with nine sampling time points for the following experiments as a compromise between throughput increase and accuracy of flipping profiles (98% of sequences below 5% approximation error). Afterwards, we simulated how the total library size affects the throughput by estimating the number of variants above different read-count thresholds (Fig. 2e, Methods). Here, the read-count threshold is defined as the minimum number of NGS-reads per variant and time point above which the functional readout (fraction flipped) is considered statistically robust. The throughput is the number of variants above this threshold. At a given limit of obtainable reads per NGS run, the total library size (i.e. the number of variants subjected to NGS) represents the main experimental parameter that can be tuned to adjust the overall throughput of our method. Increasing total library size is expected to increase the throughput but also to lead to a higher relative fraction of RBSs below threshold. Our analysis indicated that a library of approximately 250,000-500,000 RBS variants would be optimal to robustly retrieve high-quality (i.e. above-threshold) data for a maximized number of variants (Fig. 2e).

### Ultrahigh-throughput Characterization of RBSs

We created a second, larger RBS library diversifying all 17 bases directly upstream of the *bxb1* start codon (N_17_). Such libraries are known to be prone to strong skew towards weak RBSs^44^, which we also observed for the first RBS library (Supplementary Fig. 7a). Initial efforts on training a machine learning (ML) model on these data indicated a systematic underestimation of translation activity particularly for strong variants (Methods, Supplementary Fig. 7b). This observation, which we attributed to the skew in the initial library, prompted us to construct three additional libraries (High1-3) likely enriched for intermediate and strong RBSs. Libraries High1-3 were designed based on the first dataset and added to an approximate total of one fifth to the N_17_ library (Fig. 3a, Supplementary Fig. 8, Methods). The composite library (∼350,000 pooled transformants) spiked with a set of 31 internal-standard RBSs spanning a wide range of activities (Methods) was subjected to the uASPIre workflow. This experiment yielded the fraction flipped for 303,503 RBSs over nine time points constituting over 2.7 million sequence-function pairs (Fig. 3b). The applied threshold of at least 20 reads per RBS and time point corresponds to a robust minimum coverage of 180-fold for each variant with the average coverage amounting to 587-fold. This threshold resulted from comparing the predictive performance of ML models (see below) trained on datasets with different thresholds and tested on a validation set (Supplementary Fig. 9, Methods). Notably, while the same NGS platform was used, the throughput was increased about 29-fold compared to the proof-of-concept experiment due to optimized sampling and library size. This experiment was done in three independent biological replicates with low variability between replicates substantiating high reproducibility of uASPIre (Supplementary Fig. 10).

**Fig. 3.**
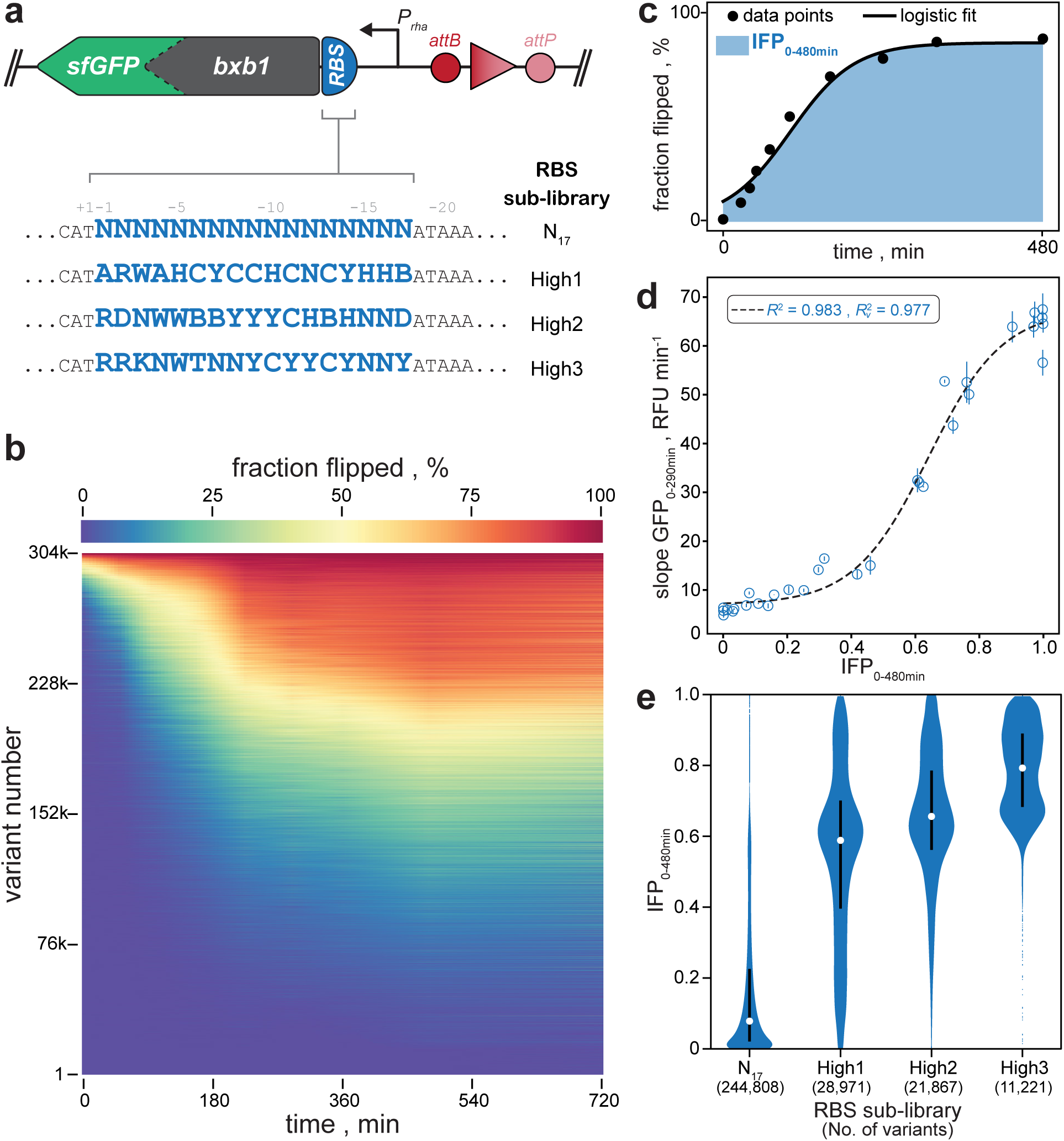
Ultrahigh-throughput characterization of RBSs by uASPIre. **a**, Diversification strategy for RBSs. Four RBS sub-libraries were designed randomizing 17 bases in the 5’-UTR with full (N_17_) and partial (High1-3) degeneracy. High1-3 are designed to achieve a predicted reduced skew towards weak RBSs (Methods). Sequences are displayed in reverse complement. **b**, Kinetic behaviour of 303,503 RBS variants from the composite RBS library. Horizontal lines are the 9-time-point flipping profiles with fraction flipped shown in colour code, and variants are ranked according to their IFP_0-480min_. Data corresponds to more than 2.7 million sequence-function pairs recorded in a single Illumina NextSeq run. **c**, Schematic representation of IFP_0-480min_ (blue shaded area). **d**, Correlation of IFP_0-480min_ and cellular Bxb1-sfGFP fluorescence as shown for 31 internal-standard RBSs spanning a wide range of activities (Methods). R^2^ and R ^2^ are coefficient of determination and coefficient of determination after leave-one-out cross validation for the logistic fit (dashed line), respectively. **e**, Distribution of IFP_0-480min_ in the four sub-libraries. Violins contain percentiles 0.5 to 99.5 of variants with median and outliers as white and blue dots, respectively. Black bars contain the 25^th^ to 75^th^ percentiles.

To correlate the functional readout (fraction flipped) obtained by NGS with the cellular concentration of Bxb1-sfGFP, we recorded cell-specific fluorescence of the Bxb1-sfGFP fusion for the aforementioned set of 31 internal-standard RBSs in individual shake flask cultivations (Supplementary Fig. 11, Methods). We compared the resulting curves with the corresponding flipping profiles obtained in NGS by analysing pairs of integral- and slope-based summary statistics (i.e. quantitative curve representations) for the two measurement types (Methods). Discriminator inversion strongly correlated with the prevailing cellular Bxb1-sfGFP concentration as indicated by high R^2^ values (0.85-0.98) for all tested combinations (Supplementary Fig. 12a,b). We selected the integral of the flipping profile between 0 and 480 minutes after induction (IFP_0-480min_, Fig. 3c) for further steps due to its high correlation with different summary statistics for Bxb1-sfGFP fluorescence and the high degree of diversity for IFP_0-480min_ in the library (Fig. 3d, Supplementary Fig. 12c). Notably, in comparison to the GFP measurements, the functional readout obtained by NGS exhibited a larger dynamic range and higher sensitivity at the lower (and to a lesser extent higher) range of the RBS activity spectrum (Fig. 3d). Relying on IFP_0-480min_, we assessed different sub-libraries (Fig. 3e). As expected for fully degenerate RBS diversification, we found the N_17_ sub-library to be strongly skewed towards low activity. By contrast, all three designed sub-libraries (High1-3) were enriched for intermediate and strong RBSs, indicating that our design goal was met.

### Deep Learning for RBS Prediction through SAPIENs

We developed a machine learning approach to exploit the datasets obtainable by uASPIre for quantitative prediction of RBS strength from sequence. Our deep learning model SAPIENs uses the RBS sequence as an input in a binary matrix representation (Fig. 4a, Methods). The SAPIENs architecture is an ensemble of ten residual neural networks (ResNets)^47, 48^, each consisting of three residual blocks of two convolutional layers^49^ each. The last convolutional layer’s output is fed into two sets of fully connected layers, which integrate information across all positions of the RBS. These two layer sets provide the final output of each ResNet of the ensemble, which is two shape parameters of a probability distribution for RBS activity (beta distribution). The ten ResNet models were independently trained with different randomly initialized parameters, random hyperparameters and batches of sequences. In this way, SAPIENs models the predicted distribution of IFP_0-480min_ as a uniformly weighted mixture of ten beta distributions, parametrized by ten independent sets of shape parameters. The mean of the mixture distribution is the predicted IFP_0-480min_ and the standard deviation is a measure of prediction uncertainty (Methods). Crucially, this does not only provide quantitative predictions of the RBS activity but also a well-calibrated confidence score. A detailed description of all components of SAPIENs is available in the ML Annex.

**Fig. 4.**
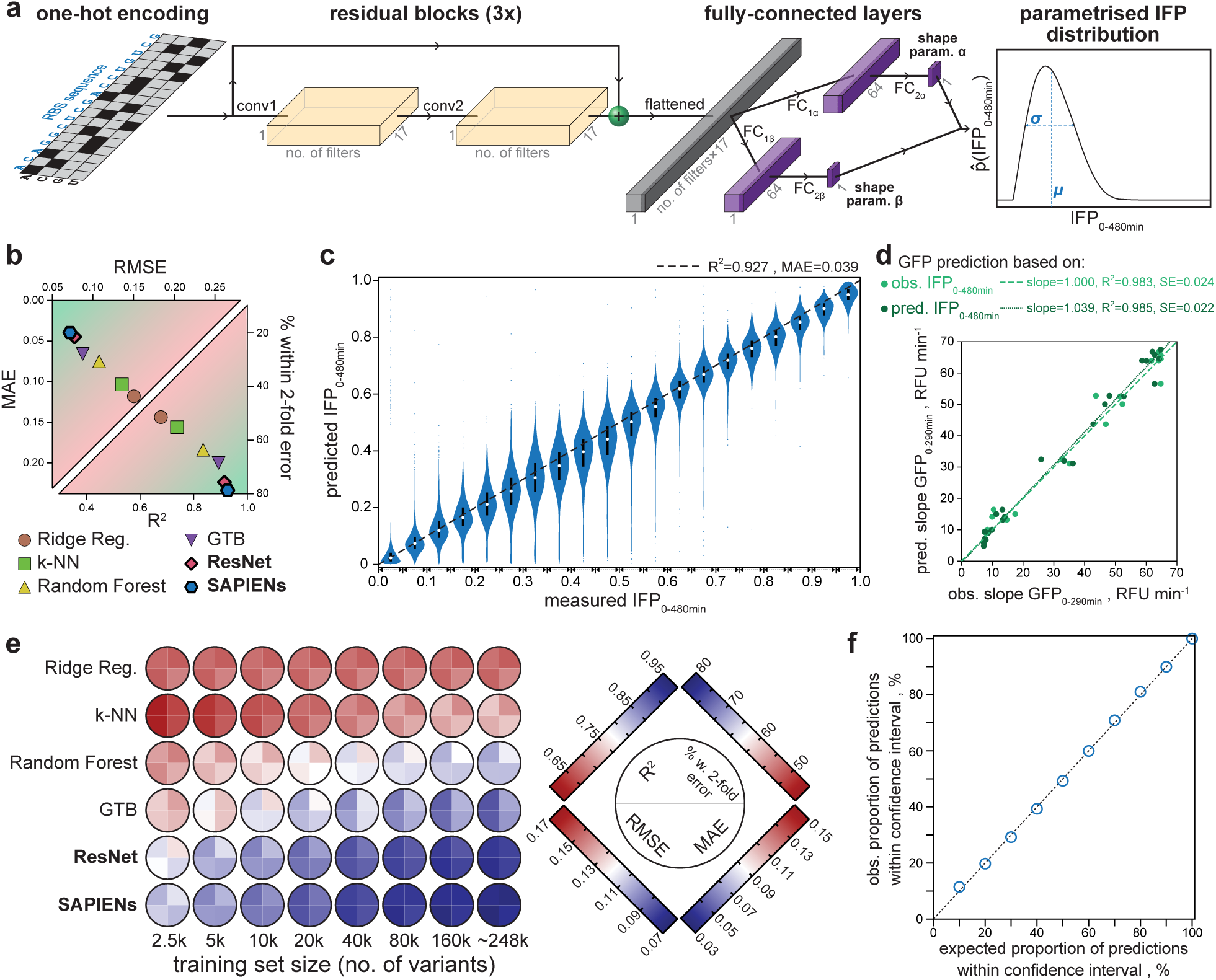
Quantitative prediction of RBS activity with SAPIENs. **a**, Schematic architecture of a single ResNet. One-hot encoded 17-bp RBS sequences are fed into three residual blocks, composed of two convolutional layers (conv1, conv2), and two sets of two fully-connected layers (FC_1*α/β*_, FC2*_α/β_*). Yellow and purple boxes represent the output of the convolutional and fully-connected layers, respectively. The grey box represents the output of the flattening operation. The model yields a probability distribution of IFP_0-480min_ for each sequence from which the corresponding predicted IFP_0-480min_ value (mean *µ*) and an uncertainty estimate (standard deviation *σ*) can be calculated. SAPIENs overall architecture is a combination of ten individually parametrized ResNets. **b**, Comparison of the predictive performance of single ResNet and SAPIENs with classical linear and non-linear ML models trained on the same set of 248,451 RBSs (Methods). MAE: mean absolute error, RMSE: root-mean-square error. **c**, Comparison of IFP_0-480min_ values as predicted by SAPIENs with the corresponding experimental values measured by uASPIre. Sequences in the test set were binned (bin size: 0.05) according to measured IFP_0-480min_. Violins comprise percentiles 0.5 to 99.5 of predicted values per bin with median and outliers represented as white circles and blue dots, respectively. Black bars contain the 25^th^ to 75^th^ percentiles. **d**, Cellular Bxb1-sfGFP concentrations can be reliably predicted from experimentally determined as well as predicted IFP_0-480min_ values as shown for the 31 internal-standard RBSs. IFP_0-480min_ values were converted into the slope of the cell-specific Bxb1-sfGFP signal between 0 and 290 min after induction relying on the logistic fit parameters determined for these two summary statistics earlier (Fig. 2e). **e**, Dependence of the predictive performance of different ML models on the size of the training dataset (Methods). The predictive performance is evaluated with four metrics: percentage of predicted IFP_0-480min_ values within 2-fold error, MAE, RMSE and R^2^. The colour scheme indicates performance, from dark blue (best) to dark red (worst). **f**, The confidence intervals of the predicted probability distributions (horizontal axis) fully assess the uncertainty of the prediction values (vertical axis), i.e. x% of the predicted values lie within x% confidence interval. The obtained values (blue dots) are well aligned with a theoretical perfect uncertainty assessment (dotted line).

We trained SAPIENs and several classical linear and non-linear machine learning models on the same 248,451 RBS sequences chosen at random from the larger uASPIre dataset. Hyperparameters were optimized exclusively on a validation set (∼30,000 sequences) and afterwards all models were evaluated on a held-out test set (∼30,000 sequences, Methods). The linear model Ridge Regression^50^ (R^2^=0.678) was clearly outperformed by non-linear models k-nearest neighbors^51^ (k-NN, R^2^=0.738), Random Forest^52^ (R^2^=0.835) and gradient tree boosting^53^ (GTB, R^2^=0.893), which highlights the importance of interactions between nucleotides in the RBS (Fig. 4b,c). Notably, SAPIENs outperformed all other approaches reaching an R^2^ of 0.927 and MAE of 0.039. It exhibited consistently high predictive performance across the entire range of RBS activities including the 31 internal-standard RBSs that were excluded from training (Fig. 4c, Supplementary Fig. 13). Moreover, the systematic inaccuracy in predicting strong RBSs was eliminated as a result of the addition of the three designed sub-libraries

High1-3 (Supplementary Fig. 14). The predicted IFP_0-480min_ values were converted into summary statistics for cellular Bxb1-sfGFP concentrations relying on calibration curves (Fig. 3d) and the resulting predicted Bxb1-sfGFP values correlated well with their experimentally determined counterparts (Fig. 4d). This indicates that our model reliably predicts cellular protein levels even for unseen sequences. Importantly, except for the overall weakest-performing Ridge Regression, prediction accuracy increased with training set size for all models as reflected by rising confidence (R^2^, percentage of sequences within 2-fold error) and decreasing errors (RMSE, MAE; Fig. 4e). While a general trend towards saturation was observed, no plateau is reached even for the largest training set of 248,451 sequences. This points to the high value of large-scale sequence-function data and emphasizes the potential of uASPIre. Moreover, to go beyond global metrics of accuracy (R^2^, MAE), SAPIENs produces a well-calibrated confidence score for each prediction^54^, which can be used to guide forward engineering of RBSs (Fig. 4f, Supplementary Fig. 15, Methods). Lastly, we observed that our high prediction accuracy is reproducible across the three biological replicates (Supplementary Fig. 16, ML Annex).

**Fig. 5.**
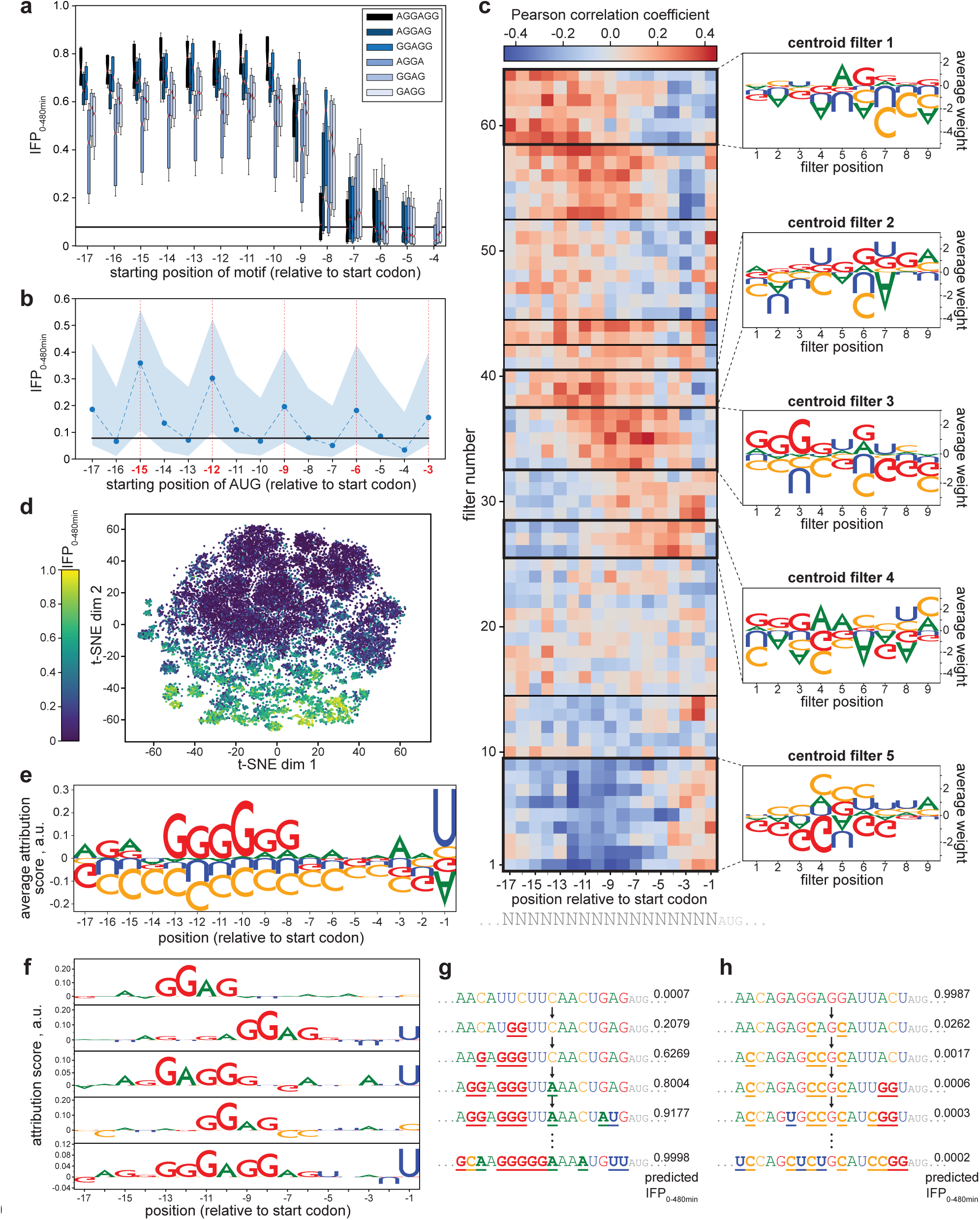
Interpretation of uASPIre data and SAPIENs. **a/b**, Influence of Shine-Dalgarno-like motifs (**a**) and AUG codons (**b**) in the 5’-UTR on the RBS activity as represented by IFP_0-480min_. The black horizontal line corresponds to the median IFP_0-480min_ in the dataset. Boxes in (**a**) contain the interquartile range with median (red box centres) and percentiles 20/80 (whiskers). Circles in (**b**) represent median IFP_0-480min_ with shaded areas comprising percentiles 20/80 and in-frame positions highlighted in red. **c**, Importance of filters of the ResNet model for the prediction of RBS activity. The Pearson correlation between a filter’s activation and the RBS activities of all held-out sequences is displayed per filter and per position for the first convolutional layer of one randomly selected ResNet model from SAPIENs. Five filter stacks with apparent high significance for the model are framed in bold and the average weight per base and position of the corresponding centroid filter is shown to the right. **d**, Visualization of the integrated gradients scores of SAPIENs in a low-dimensional space. T-distributed stochastic neighbour embedding (t-SNE) is applied to the integrated gradient scores of the RBS sequences from the test set. t-SNE dim1/2 are the two dimensions resulting from the t-SNE algorithm. **e**, Impact of bases and positions in the 5’-UTR on the RBS activity. Using an all-zeroes input as baseline, the average attribution score per base and position is displayed as determined for the sequences in the test set. The size of the letters corresponds to their importance score and their orientation to the direction of effect (i.e. upward/downward corresponding to a tendency to increase/decrease IFP_0-480min_). **f**, Attribution of bases and positions to strong RBSs. The strongest 5% of sequences in the test set were distributed into five clusters using k-means algorithm. The displayed motifs are the five medoids of each cluster (i.e. the five individual sequences closest to the respective cluster centroid). **g**/**h**, *In silico* evolution of RBSs. Starting from the sequence with the lowest (**g**) and highest (**h**) predicted IFP_0-480min_ in the test set, pairwise mutations are greedily applied until no further increase (**g**) or decrease (**h**) in IFP_0-480min_ is observed (total of 10 and 8 rounds for (**g**) and (**h**), respectively).

### Identification of Sequence Determinants of RBSs

We analysed the impact of different factors known to influence RBS activity in our data. In contrast to other studies^55^, we did not find significant correlation between mRNA folding energy and RBS activity regardless of the selected sequence window. This may be due to the full randomization of only a short part of the mRNA rendering strong secondary structures unlikely and thus underrepresented. We then assessed the impact of Shine-Dalgarno (SD)-like motifs (i.e. AGGAGG and subsequences thereof) and additional start codons in the 5’-UTR within the N_17_ library. Clearly, SD-like motifs exhibit a strong positive effect on translation, which is lost (or even slightly inverted) if the motif is too close to the translational start (Fig. 5a). Similarly, a positive effect was observed for additional in-frame AUG codons (Fig. 5b) and, to a lesser extent, for GUG and UUG (Supplementary Fig. 17). By contrast, out-of-frame start codons showed no globally consistent tendency but overall favoured translation, in particular for positions -17 to -8. This is likely due to Gs in the start codons facilitating 16S-rRNA binding, which expectedly is most prevalent for GUG (Supplementary Fig. 18) and difficult to disentangle from a genuine start codon effect.

We analysed one ResNet model to gain an understanding about the relative importance of RBS bases and positions by clustering filters of the first convolutional layer according to the correlation between their output features and the RBS activity (Methods). We found that the model had captured translation-promoting (A, G) and translation-reducing (C) effects of bases (Fig. 5c). Moreover, a positioning effect was observable: filters with large positive weight for Gs or negative weight for Us/Cs correlated positively with RBS activity when scanning upstream regions but negatively when closer to the translational start (centroids 1-4). By contrast, filters promoting Us/Cs correlated negatively with RBS strength for most positions (centroid 5).

For further interpretation, we used integrated gradients^56^, an attribution method commonly used for deep learning models (Methods). For the test set, a low-dimensional embedding of SAPIENs integrated gradients scores indicated a clear structure with strong and weak sequences clustering in almost linearly separable fashion (Fig. 5d). Global analysis of the integrated gradients scores revealed that specific positions and bases are particularly indicative for RBS activity (Fig. 5e). Substantiating the observation from Figure 5c, Gs strongly promote translation while Cs appear to be consistently adverse. The translation-promoting effect for Gs is only observable if the distance from the start codon is at least seven bp, while a neutral or even unfavourable effect prevails for other regions. However, no distinct SD-like motif appeared because this global analysis only represents per-base and -position averages. A more targeted analysis obtained by clustering of strong and weak sequences (Fig. 5f, Supplementary Fig. 18, Methods) revealed SD-like motifs with most impactful positions ranging from -13 to -6 and invariance or slight preference for weakly pairing bases (A, U) outside the motif. Hence, our model successfully reconstructed SD-like patterns, notably without any prior knowledge about the process of translation.

Finally, we used SAPIENs to perform *in silico* evolution by greedily applying pairwise mutations to the weakest and strongest sequence in the test set to maximize and minimize predicted IFP_0-480min_, respectively (Methods). Confirming our previous findings, the model systematically mutated U or C to A or G to form SD-like motifs or create in-frame start codons upon increasing RBS strength (Fig. 5g), whilst removing Gs and adding Cs when decreasing it (Fig. 5h). Moreover, evolving a strong sequence (“gain of function”) required more steps than diminishing RBS activity (“loss of function”) due to the sparsity of strong sequences within the search space.

## Discussion

Herein, we introduce a novel method termed uASPIre, which relies on phenotypic recording in DNA to enable experimental generation of large-scale sequence-function data of high quality while significantly reducing experimental effort and minimizing error. Since uASPIre only relies on NGS as experimental readout, an extremely high throughput can be achieved, which scales linearly with the number of obtained sequencing reads and is independent of other technical parameters such as sorting speed and efficiency. As an example, transferring the experimental setup used in this study to currently available benchmark NGS systems (e.g. Illumina NovaSeq 6000) would allow recording of 10^8^ or more sequence-function pairs per experiment, for which further increases are to be expected in the future due to the ongoing development of the NGS field.

Notably, the uASPIre approach requires no sophisticated instruments such as cell sorters or specialized facilities (except for NGS which can be outsourced to service providers) and only standard methods for sample preparation (DNA purification, restriction, ligation). Determination of library composition and its functional characterization are performed concomitantly and in a single device. Compared to previously available methods, this grants experimental practicability and, more importantly, avoids bias and error from multiple devices and processing steps. To this end, neither barcoding nor amplification of DNA after the actual experiment (e.g. via PCR or clonal expansion by growth) are required, both of which are known sources of bias (compare Supplementary Fig. 5 and 6). Variant treatment is fully parallelized throughout the entire workflow from library generation to final readout, which avoids introduction of bias due to long and/or differential processing times, constituting a major advantage over sequential approaches. Importantly, functional information is recorded directly and does not have to be statistically inferred from read distributions, which is a well-known source of experimental error^22^.

The corresponding functional readout (i.e. fraction of modified discriminators) is a quantitative, internally normalized metric for variant comparison. It exhibits high sensitivity and large dynamic range as can be appreciated from its superior ability to resolve differences between variants at the low and high end of the activity range compared to fluorescence measurements with sfGFP (Fig. 2d). Its resolution can be arbitrarily adjusted by adapting the sequencing depth (i.e. number of reads obtained per variant) and could be further enhanced using systems that allow more than two discriminator states^57, 58^. The instantaneous and continuous recording of the functional readout *in situ* avoids the need for immediate measurements during or directly after the cultivation, which are for instance required in the case of transient reporters such as fluorescent proteins. Therefore, the kinetic resolution of uASPIre is only limited by the time required for sampling of the culture, which can be performed in intervals of one minute or less. This is a key feature, since most biological phenomena are highly dynamic and therefore inappropriately depicted by end-point measurements as our data on RBSs show (Figs. 2c and 3b). Crucially, such high-resolution kinetics cannot be achieved in approaches that rely on elaborate and lengthy procedures such as cell sorting.

In this study, we capitalize on these advantages and demonstrate the utility of uASPIre by recording the effect on translation of more than 313,000 RBSs from two libraries with in total over 2.9 million sequence-function pairs. Furthermore, we exploit the resulting high-quality datasets by deep learning to quantitatively predict the behaviour of RBSs. Notably, only the combination of big data obtained through uASPIre and the model SAPIENs facilitated the high predictive performance achieved in this study (Fig. 4e), emphasizing the potential of high-throughput experimentation combined with state-of-the-art deep learning. SAPIENs accurately quantifies the uncertainty of its predictions, which can be used as a practical criterion to pick sequences for which the prediction is most reliable. Additionally, interpretation of the uASPIre data and SAPIENs revealed position-specific sequence motifs in a fully data-driven fashion without requirement for prior knowledge about RBSs.

Importantly, uASPIre is not restricted to specific functional traits of interest and the approach introduced herein can be repurposed to address a wide range of biologically relevant questions. To this end, RBS library characterization should be viewed as an application example and uASPIre can be used to interrogate different types of GREs and the mechanisms of gene regulation on all levels of the central dogma. This is of high significance since gene regulation is of utmost importance for cellular function and impaired regulation of genes is frequently associated with disease. Moreover, we anticipate the utility of uASPIre also beyond the realm of gene regulation arguing that, in principle, any trait of interest, which can be coupled to a gene expression output, could be accessed with the method. For instance, transcriptional or translational biosensors may be used to drive modifier expression in response to certain stimuli or small molecules of interest rendering a plethora of alternative applications accessible. We therefore envision a wide applicability of the uASPIre approach in several research domains including metabolic engineering, genetic circuit design and microbiome research, to name but a few.

## Supporting information

Machine Learning (ML) Annex

## Acknowledgments

The authors kindly thank all members of the Genomics Facility Basel for support with NGS experiments, Prof. Michael C. Jewett for helpful scientific discussions, and Dr. Tania Roberts for experimental advice. This work was supported in part by the SNSF Starting Grant “Significant Pattern Mining” (K.B.) and the NCCR “Molecular Systems Engineering” (Y.B.).

## Author contributions

M.J. and Y.B. conceived the project. M.J., Y.B. and K.B. coordinated the study. K.B. conceived and supervised machine learning analyses. M.J. supervised experimental work. S.H., K.F. and M.J. performed experiments. S.H., K.F., L.P., A.C.G. and M.J. analysed data. A.C.G., L.P. and M.J. developed measures to increase throughput. S.H. and L.P. developed the algorithm for processing of NGS data. L.P. conceived, developed and analysed ML models. C.B. advised design of DNA adapters and NGS. M.J., L.P., and Y.B. wrote the manuscript with input from all authors.

## Methods

### Chemicals and Reagents

Unless stated otherwise, all chemicals and reagents were obtained from Sigma Aldrich (Buchs, Switzerland). Enzymes were obtained from New England Biolabs (Ipswich, MA, USA). Oligonucleotides (Supplementary Tab. 1), custom duplex DNA adapters, synthetic genes and gene fragments were obtained from Integrated DNA Technologies (Leuven, Belgium).

### Cultivation of E. coli

*E. coli* strains were commonly cultivated in lysogeny broth (LB)^1^ supplemented with 10 g L^-1^ D -glucose, 50 mg L^-1^ kanamycin, 50 mg L^-1^ streptomycin and 15 g L^-1^ agar where appropriate. Rhamnose-utilization deficiency was assessed by cultivation of strains in defined mineral medium^2^ supplemented with 0.1 g L^-1^ L-leucine, 0.03 g L^-1^ L-isoleucine, 0.15 g L^-1^ L-valine and 10 g L^-1^ of either D -glucose or L-rhamnose as major carbon source. Shake flask cultures (LB) were inoculated from monoclonal pre-cultures to an initial OD_600_ of 0.05 and cultivated in a shaking incubator (37 °C, 200 rpm). Expression was induced at an OD_600_ of ∼0.5 by addition of 2 g L^-1^ L-rhamnose to the cultures. Microtiter plate cultivations were performed in sterile 96-well plates (flat bottom Nunclon™ Delta Surface, Thermo Fisher Scientific, Waltham, MA, US) containing 200 μL LB per well. Wells were inoculated from monoclonal pre-cultures to an initial OD_600_ of 0.05 and plates were incubated in an Infinite® M1000 PRO plate reader (Tecan Group, Männedorf, Switzerland) at 37 °C without lid (orbital shaking mode, 6 mm amplitude).

### Construction of Plasmids

Plasmids used are listed in Supplementary Table 2. All plasmids constructed in the course of this study are based on pSEVA291^3^. Inserts were created relying on synthetic genes or gene fragments, which were inserted into the vector backbone by conventional restriction-ligation cloning. Maps and sequences of plasmids created in this study can be found in the Supplementary Information (Supplementary Figs. 19-23). A detailed description of library construction procedures is provided below.

### Assessment of Bxb1 Recombination

Bxb1-mediated inversion of the discriminator was assessed by direct fluorescent measurement, counting of red-and non-fluorescent colonies after retransformation of isolated plasmid and Sanger sequencing of the discriminator region. For direct measurement of mCherry fluorescence, 1 mL samples were collected from shake flask cultivations, spun down in a tabletop centrifuge (1 min, 8’000 rcf) and pellets were re-suspended in 1 mL of ice-cold, sterile-filtered phosphate-buffered saline^1^ (PBS). To ensure full chromophore maturation, samples were incubated at 4 °C overnight before measurement in an Infinite® M1000 PRO plate reader (Tecan Group, Männedorf, Switzerland) in 96-well plates (flat bottom Nunclon™ Delta Surface, Thermo Fisher Scientific, 200 μL per well). Cell-specific mCherry fluorescence was determined by dividing the red fluorescence signal (excitation at λ_Ex_ = 587 nm, emission at λ_Em_ = 610 nm) by the OD_600_ for each sample. For microtiter plate cultivations, fluorescence of mCherry and sfGFP (λ_Ex_ = 485 nm, λ_Em_ = 535 nm) was directly measured in the culture broth and normalized for OD_600_. To assess the dynamics of Bxb1-mediated recombination directly on the DNA level, plasmid DNA was extracted from shake flask culture samples and the state of the discriminator was determined by Sanger sequencing. Furthermore, 50 ng of extracted plasmid DNA were used to re-transform *E. coli* TOP10 and the transformation mixture was plated on LB agar containing 50 mg L^-1^ kanamycin and 10 g L^-1^ D-glucose to shut down transcription from *P_Rha_*. After overnight incubation at 37 °C, plates were stored at 4 °C for maturation of the mCherry chromophore. Afterwards, colonies (at least 275) were manually counted to determine the ratio of clones that had received a plasmid copy of pASPIre1 with a flipped (red colonies) or unflipped (white colonies) discriminator upon transformation, respectively.

### Construction of Knockout Strains

All *E. coli* strains used in this study are listed in Supplementary Table 2. Knockout of the genomic *rhaA* gene (L-rhamnose isomerase) in parent strain *E. coli* TOP10 was achieved using the method described by Datsenko and Wanner^4^. Primers 1 and 2 were used to generate the required linear DNA fragment containing the kanamycin resistance gene (*kan^R^*) from plasmid pKD13^4^ flanked by sequences homologous to the target locus.

### Library Generation

RBS libraries were generated via PCR on template plasmid pASPIre3 using forward primer 3 and degenerate reverse primers 4, 5, 6, and 7 to diversify the respective RBS region. The resulting PCR products were digested with *Pst*I and *Sac*I (37 °C, 4 h) and ligated into pASPIre3 pre-treated with the same restriction enzymes. The ligation mixtures were purified and used for electroporation of *E. coli* TOP10 Δ*rhaA*. Transformants were plated on several plates of LB agar (10 g L^-1^ D-glucose, 50 mg L^-1^ kanamycin, 50 mg L^-1^ streptomycin) in order to estimate library size by colony counting and facilitate pooling of the different libraries in defined ratios. After overnight incubation (37 °C), 5 mL of LB were added to the plates and colonies were scraped off. The resulting cell suspensions were pooled and sterile glycerol was added to a final concentration of 15% (v/v). Last, OD_600_ of the cell suspensions was determined before library stocks were snap-frozen and stored at -80 °C until further use.

### Library Cultivation and Sampling

Library stocks were thawed on ice and used to inoculate 600 mL pre-warmed LB (50 mg L^-1^ kanamycin) to an initial OD_600_ of 0.05 in 5 L baffled cultivation flasks. Cultures were incubated (37 °C, 200 rpm) until an OD_600_ of ∼0.5 was reached and 2 g L^-1^ L-rhamnose were added to induce Bxb1-mediated recombination. Samples taken throughout the cultivation were immediately mixed with an excess of ice-cold, sterile PBS for rapid cooling and then centrifuged (10 min, 4’000 rcf, 4 °C), and cell pellets were frozen on dry ice until extraction of plasmid DNA was performed using a commercial kit (ZymoPURE Miniprep Kit, Zymo Research) and stored at -20 °C until further use.

### NGS Sample Preparation

Plasmid DNA isolated from culture samples was digested with *Nco*I and *Sac*I (37 °C, 4 h). After, target fragments (308 bp) containing both RBS and the *attP/R* site were purified by agarose gel electrophoresis and sample-specific combinations of customized, indexed DNA duplexes (Supplementary Tab. 3) were ligated to the sticky ends of the target fragment. For the PCR-amplified sample (Supplementary Fig. 4), NGS fragments were generated using primers 8 and 9 to specifically amplify the target region and add required overhangs for Illumina sequencing to both ends. The resulting linear DNA fragments were purified by agarose gel electrophoresis and concentration of the target was determined using capillary electrophoresis (12-capillary Fragment Analyzer, Advanced Analytical/Agilent). Afterwards, indexed samples were pooled according to their determined concentrations to adjust equal molarity for all samples and the pooled sample was subjected to NGS.

### NGS

NGS was performed using an Illumina NextSeq 500 platform and a High Output kit v2.5 (75 cycles, PE 33/51) using ∼20% genomic PhiX library as spike-in to increase sequence diversity. Primary data analyses were done with Illumina RTA version 2.4.11 and bcl2fastq v2.20.0.422.

### Computational Scripts and Datasets

An annotated script for the processing of NGS data (see below) used in this study is available under: https://github.com/JeschekLab/uASPIre. Raw and processed NGS data will be made available upon final publication. A detailed description of machine learning (ML) models is provided in the separate ML Annex. Code describing how to define and fit the SAPIENs model as well as the resulting parameters of the fitted model will be made available upon final publication.

### Processing of NGS Data

The algorithms for processing of NGS data for this project were written in bash and python and are available under https://github.com/JeschekLab/uASPIre. Briefly, forward and reverse reads retrieved from fastq files were paired and all reads with more than six consecutive unidentified nucleotides were removed. Afterwards, target fragments were selected by a 10-bp constant region (GAGCTCGCAT, max. 3 mismatches) and sequences from different samples were deconvoluted by their unique combination of two 6-bp indices (Supplementary Tab. 3). Next, the discriminator state was determined by searching for the presence of an *attP* or *attR* site corresponding to the sequences G**GGTTTG**T**ACCGTAC**AC or G**CCCGGA**T**GATCCTG**AC, respectively (max. 3 mismatches, differential bases highlighted in bold). RBS sequences were determined by retrieving the 17 nucleotides upstream of the *bxb1* start codon. Finally, variants with mismatches in the *bxb1* CDS in more than 8% of reads were removed to exclude off-target mutations.

### Construction of Internal-standard RBSs and Recording of Calibration Curves

The internal-standard RBSs used in this study are listed in Supplementary Table 4. RBSs R1-R22 were selected from the proof-of-concept library with the goal to span the entire range of observed RBS activities. First, kinetic profiles were cropped at 720 min and only RBSs with at least 100 reads per sample were used which resulted in a set of high-quality profiles for approximately 9,500 RBSs. Afterwards, profiles were grouped according to their dynamic behaviour relying on k-medoid clustering^5^ (k = 25) and the RBS corresponding to the centre of each cluster was selected as representative internal-standard RBS (Supplementary Fig. 24). Additionally, three weak (R23-R25) and four strong (R26-R29) RBSs handpicked from the initial library as well as two strong RBSs (R30, R31) designed using the RBS calculator^6^ were included. These RBSs were individually introduced into pASPIre3 by conventional cloning procedures to obtain derivatives that carry the respective RBS sequence controlling Bxb1-sfGFP translation. Activity of these RBSs was assessed by recording of the cell-specific fluorescence of three biological replicates of each variant in individual shake flask cultivations. 100 mL of pre-warmed LB (50 mg L^-1^ kanamycin) in 1 L baffled shake flasks were inoculated from overnight pre-cultures to an initial OD_600_ of 0.05. Bxb1-sfGFP expression was induced at an OD_600_ of ∼0.5 by adding 2 g L^-1^ L-rhamnose. Samples taken throughout the cultivation were immediately mixed with an excess of ice-cold, sterile PBS for rapid cooling and then centrifuged (10 min, 4,000 rcf, 4 °C). Cell pellets were re-suspended in PBS and suspensions were stored overnight in micro centrifuge tubes at 4 °C for sfGFP maturation. Fluorescence (λ_Ex_ = 485 nm, λ_Em_ = 535 nm) and OD_600_ were measured in technical triplicates in 96-well plates (Corning 96-well Clear Bottom Black Polystyrol, 200 µL per well) in a TECAN Infinite® M1000 PRO plate reader. Curves of cell-specific sfGFP fluorescence were obtained by normalizing the blanked fluorescence signal for the blanked OD_600_ measurements and subtracting the cell-specific background fluorescence of an sfGFP-less variant (empty vector control) which was included for every cultivation batch. Furthermore, a dilution series of fluorescein was included in every 96-well plate to compensate for variations of the fluorescent readout over time.

### Correlation of Bxb1-mediated Recombination with Cellular Bxb1-sfGFP Levels

In order to convert Bxb1-catalysed discriminator flipping into cellular Bxb1 concentrations, we compared the recorded cellular fluorescence profiles for the 31 internal-standard RBSs with their corresponding flipping profiles as recorded by NGS. To this end, we sought to i) establish a combination of summary statistics which exhibit a high degree of correlation between the two measured quantities across the entire range of RBS strengths, ii) identify the best (potentially non-linear) fit between the two summary statistics, and iii) ensure that a high degree of diversity is maintained for the representation of the discriminator flipping across the entire set of sequences in the dataset. We used integral-based (i.e. area under the curve) summary statistics for the flipping profiles and slope-based representations (i.e. slope of the linear fit) for the fluorescence profiles (Supplementary Fig. 12a,b). For the flipping statistics we also quantified the diversity of each representation by estimating the differential entropy^7^ of its probability density (Supplementary Fig. 12c). For each type of summary statistic, we additionally treated the time ranges over which both the fluorescence and flipping summaries are computed as additional hyperparameters to be optimized. Moreover, for each pair of candidate summary statistics, we evaluated linear, log-linear and generalized logistic fits. We quantified the quality of each pair of summary statistics using the resulting R^2^ of the fit as evaluated using leave-one-out cross-validation on the pool of 31 internal-standard RBSs in order to compensate for potential effects of overfitting in the analysis. Moreover, the standard deviation of each summary statistic for fluorescence was computed for all internal-standard RBSs relying on the three biological replicates. Further details on the evaluation of summary statistics are provided in the ML Annex.

### Optimization of Sampling Time Points

Sampling times were optimized using the high-quality kinetic profiles obtained from the proof-of-concept RBS library (see previous section for definition of high-quality profiles). To avoid biases towards the initial sampling schedule, we first represented the profile *p* of each RBS by an approximation with a logistic function 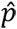 imputed at 5 min intervals, which was fixed as the minimal time difference between two samples (Supplementary Fig. 25a). In cases where logistic approximation was not possible (i.e. failed parameter optimization), an exponential decay function was used. Formally, RBS *i* is represented by its approximation 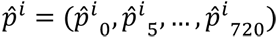. Afterwards, optimal sampling times were greedily selected while fixing the first and last sample at 0 min and 720 min after induction. The goal of the optimization was to find a set *S* of optimal sampling times (initially *S* = (0, 720)) which allows to reconstruct 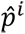 such that a linear approximation using time points in S is as close to 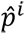 as possible. Given the set of possible sampling times *T* = {0, 5, 10, …, 720} and the subset *I* of RBS profiles on which the sampling times should be inferred, the greedy optimization finds the next optimal sampling time point *s*\* from *T* as follows:

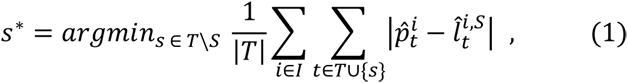

where 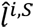 corresponds to the linear approximation of 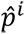 using only sampling times in *S* (Supplementary Fig. 25b). In other words, *s*\* is the time point that (i) is not part of the sampling schedule *S* yet and (ii) results in the smallest cumulative reconstruction error over all RBS profiles in *I*. Subsequently, *S* is augmented by *s*\*, and equation (1) is evaluated to find the next optimal *s*\*, until *S* contains the desired number of sampling times. Finally, the quality of the optimal sampling schedule for every RBS *i* is evaluated by computing the approximation error *r^i^* between the observed profile *p^i^*, and its linear interpolation at the optimal time points *S*, termed *l^i,S^* (Supplementary Fig. 25c):

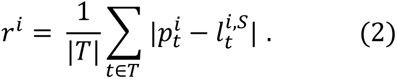

This optimization was performed for different sets of RBS profiles *I*: we first sorted RBSs by their observed strength (i.e. difference in the fraction of flipped discriminators between first and last sample) and optimized sampling for the top 5%, 10%, 25%, 50% and 100%, respectively. This strategy was chosen to compensate for the strong bias towards weak RBSs in the initial library. Afterwards, we computed the cumulative approximation error on the entire library (top-100%) for the sampling schedules optimized on the different subsets. We found that for seven or more samples the difference in approximation error between subsets became indistinguishable and hence chose the sampling times inferred on the top-10% of profiles for the following experiments.

### Optimization of NGS Loading

To increase the throughput of uASPIre, we analysed the data from the proof-of-concept (poc) experiment, which contained kinetic data of approximately 10,000 RBS variants. We sought to estimate an optimal number of variants to be loaded into NGS in order to retrieve a maximized number of variants with high quality data (i.e. above different minimal read count thresholds *θ*). For this simulation, we assumed that the limiting factor is the NGS throughput and that the maximal number of valid reads (i.e. reads that pass the pre-processing pipeline quality constraints) retrieved by NGS is constant across experiments under the same experimental conditions. This simulation is based on the idea that increasing the number of RBS variants reduces the coverage and *vice versa*, as the maximum number of valid reads is constant. For the distribution of read counts, we assumed that it follows a log-normal distribution and that its variance is independent of the coverage. The proof-of-concept dataset is composed of approximately 2×10^8^ valid reads, which are spread among *n_t_* = 18 time points and *n_poc_* = 10,427 variants with an average coverage of *cov* ∼ 1000 reads per variant per time point. If the coverage of the small dataset is reduced by a factor of *r_c_* > 1, and the number of time points by a factor of *r_t_* > 1, the total number of variants that could be loaded into NGS without loss would be *n_input_ (r_c_*, *r_t_)* = *n_poc_*×*r_c_*×*r_t_*, by conservation of the maximal number of valid reads. However, out of these *n_input_ (r_c_*, *r_t_)* variants, only *n_output_ (θ, r_c_, r_t_)* < *n_input_ (r_c_*, *r_t_)* would pass the quality control as enforced by the minimal read threshold *θ*. To simulate the effect of the minimal read threshold, we downsampled the read counts of the proof-of-concept dataset by a factor *r_c_* and applied to it the minimal read threshold *θ* resulting in a number of variants above threshold *n_simul_ (θ, r_c_)* < *n_poc_*. The estimated final number of variants is therefore *n_output_ (θ, r_c_, r_t_)* = *n_simul_ (θ, r_c_)*× *r_c_* × *r_t_*. Figure 2e illustrates the estimation of the number of variants for *r_t_* = *n_t_* / 9, several minimal read thresholds *θ* and several downsampling factors *r_c_*.

### RBS Library Design

Initial efforts for training a convolutional neural network (CNN)^8^ based on the proof-of-concept dataset resulted in a systematic underestimation of RBS strength, in particular for strong RBSs. This is likely due to the library being skewed towards weak sequences as a result of the full randomization of the 17 bases upstream of the Bxb1 start codon (Supplementary Fig. 7). To overcome this, three libraries (High1-3) presumably enriched in moderate-to-strong RBSs were designed *in silico* based on the proof-of-concept dataset and added to a fully randomized library (N_17_). Libraries High1 and High2 were designed using position probability matrices (PPMs), 2D matrices in which each element represents the proportion of times a nucleotide occurs at a given position in the sequence. To this end, RBSs from the proof-of-concept dataset were grouped into 10 linearly distributed bins according to a proxy for the normalized integral of their flipping profile (IFP_trz_, contained in [0, 1]), for each of which a PPM was computed. The IFP_trz_ was computed using the trapezoidal rule on the flipping profiles. Degenerate RBS sequences for High1 and High2 were designed with the goal to obtain PPMs that most closely resemble (minimal mean-squared error) the PPMs of the highest and second highest bin, respectively. Library High3 was designed using a genetic algorithm on the basis of predictions from an initial CNN trained on the proof-of-concept dataset. The RBS sequences from the three highest IFP_trz_ bins were randomly mutated for 200 iterations (1-2 mutations per sequence and iteration). Only sequences for which the predicted IFP_trz_ was increased due to the mutations were propagated to the next iteration. At the end of this process, we calculated the PPM of the resulting pool of sequences with high predicted IFP_trz_, randomly selected 20,000 sequences from this PPM, and computed the predicted IFP_trz_ distribution for this sub-sample. Finally, the degenerate RBS sequence of High3 was obtained by greedily minimizing the Kolmogorov-Smirnov distance between the predicted IFP_trz_ distribution of the sub-sample and the predicted IFP_trz_ distribution for the respective degenerate candidate RBS sequence. For further details regarding the computation of the IFP_trz_, the CNN and the genetic algorithm please refer to the ML Annex.

### Normalization of Biological Replicates

In order to facilitate comparison of biological replicates we capitalized on the 31 internal-standard RBSs. These serve as internal references spanning a large range of RBS activities and allow to compensate for potential batch effects and other systematic biases between replicates. Formally, for each of the 31 internal-standard RBSs, we denote by *x* and *y* the measured normalized integral of the flipping profile (IFP) for the biological replicate to be normalized and the reference replicate, respectively. We fit either a polynomial function of degree two, *f*: [0, 1] → ℝ with *f*(*x*) = *I* + *A_x_* + *B_x_*^2^, or its inverse *f*(*x*) = *g*^−1^(*x*) with *g*(*z*) = *I* + *A_z_* + *B_z_*^2^, such that the mean squared error between *f*(*x*) and *y* is minimized across the 31 measurement pairs. Moreover, we impose the following constraints on the parameters of *f*: first, RBSs that show no activity in one replicate should remain inactive in the other replicates (*f*(0) = 0). Second, RBSs whose discriminators are entirely flipped before induction in one replicate should exhibit that behavior in the other replicates (*f*(1) = 1). Third, the ranking of RBSs according to their strength should be preserved across replicates (*f* is monotonically non-decreasing in [0, 1]). It should be noted that, empirically, these assumptions appear to hold across the three biological replicates in this study. Imposing the first two constraints above reduces the number of free parameters of the polynomial function from three to one, resulting in the family of functions parametrized by *A*: *f*(*x*) = *A_x_* + (1 − *A*)*x*^2^. Moreover, the third constraint translates into the following bounds on the set of allowed values for the free parameter *A*: 0 ≤ *A* ≤ 2. This procedure was carried out for each pair of biological replicates. The quality of the resulting fits was then evaluated on the full datasets, excluding the 31 internal-standard RBSs that were used to optimize *A*.

### Machine Learning Core Model

We fitted the flipping profile of each RBS with a generalized logistic function (ML Annex), integrating the fitted kinetic curves between the time points at 0 and 480 minutes and normalized the integral value by dividing by 480 (minutes). The resulting normalized integral value (range between 0 and 1; IFP_0-480min_) was used as a descriptor of RBS behaviour and was selected as an exemplary target for prediction since it exhibits high correlation with cellular Bxb1-sfGFP levels and a high diversity across the RBS libraries (Supplementary Fig. 12). Initially, we defined a set of preliminary candidate deep-learning architectures for a predictive model according to standard practices^9-11^. These included convolutional neural networks (CNNs) with and without residual blocks, as well as multilayer perceptrons. These architectures were assessed as part of the hyperparameter selection process, which indicated superior performance of the CNN with residual blocks (ResNet)^12, 13^ for this particular application, resulting in a model with three residual blocks of two convolutional layers and two sets of two fully-connected layers. We applied three main variations to the ResNet model in order to improve predictive accuracy and additionally provide a measure for predictive uncertainty. First, we chose the negative log-likelihood, which is a proper scoring rule, as the training criterion to achieve better uncertainty estimates^14^. The predicted IFP_0-480min_ was modelled using a beta distribution, as it provides a flexible distribution with support in the interval [0, 1]. Second, the last two fully-connected layers in the network were modified to output two values instead of one, thereby allowing to independently parametrize the two shape parameters of the predictive beta distribution for each input sequence. Equivalently, as the first two moments of the beta distribution are functions of the shape parameters, we were able to retrieve the mean and the standard deviation of the predictive distribution for each input sequence. Third, we used an ensemble of N = 2×5 ResNet models^14^, each trained separately with a different random initialization of network parameters, a random order of training sequences during stochastic gradient-based optimization and different architecture and optimizer hyperparameters. This third variation helped increase predictive accuracy and capture epistemic uncertainty. The final model, SAPIENs, is an ensemble composed of five ResNet models with three residual blocks of two convolutional layers, composed of 64 filters of sizes 9 and 1, respectively, followed by two sets of two fully connected layers with 64 and 1 units respectively (weight decay parameter: 10^-6^, learning rate: 0.01) and five ResNet models with three residual blocks of two convolutional layers, composed of 512 filters of sizes 10 and 1, respectively, followed by two sets of two fully connected layers with 64 and 1 units respectively (weight decay parameter: 10^-6^, learning rate: 0.001). In all cases, we kept a held-out test set and split the remaining dataset into a training and a validation set while keeping the same proportion of strong RBSs as defined by the 15th percentile of the IFP_0-480min_ distribution and softplus activation functions for the two output layers. We used batch-normalization^15^ followed by LeakyReLU activation functions^16^ between each layer. For optimization, we used the Adam optimizer^17^. The model was implemented in Keras with the Tensorflow^18^ backend. All hyperparameters (number of filters and layers, filters sizes, number of units of the fully-connected layers, weight decay, learning rate, batch size) were selected with random search^19^ on the basis of their performance on the validation set. Additional details about the neural network can be found in the ML Annex.

### Uncertainty Estimation

The measured IFP_0-480min_ for each RBS was modelled as a draw from a beta distribution. The mean and variance of this distribution estimated by the ResNet model (see above) correspond to the predicted IFP_0-480min_ value and an indication of the aleatoric uncertainty of prediction, respectively. To complement this aleatoric estimate with an estimate of epistemic uncertainty, we first used an ensemble of N=5 ResNet models with identical architecture and optimizer hyperparameters but different random parameter initialization and ordering of the input sequences. The uncertainty estimate is therefore given by the standard deviation of the mixture of N=5 beta distributions (Supplementary Fig. 15a-c). Furthermore, we extended this ensemble strategy at a later stage by also including M different configurations for the higher level hyperparameters, such as architecture and optimizer hyperparameters, with five ResNet models per configuration, resulting in a total of N=M×5 ResNet models in the ensemble. Finally, a number of configurations M=2 was fixed as a trade-off between predictive performance and computational complexity (Supplementary Fig. 15d). The reliability diagram for this final ResNet ensemble (SAPIENs, N=2×5) showed well-calibrated uncertainty estimates (Fig. 4f) indicating that the uncertainty of each predicted target value seems to be accounted for. This is confirmed by the fact that the mean absolute error is positively correlated with the predicted standard deviations (Supplementary Fig. 15e). Both these results suggest that the predicted standard deviations can be used as scores to evaluate the quality of each individual prediction.

### Minimal Read Number Threshold

A minimal threshold for the number of NGS reads per RBS was determined as a quality control criterion for both training and test sets. Increasing this threshold is expected to trade off two opposite effects since it increases the average quality of the data leading to a decrease in the underlying aleatoric uncertainty but at the same time reduces the dataset size available for training, which generally lowers predictive performance. To this end, we first defined six filtered datasets obtained by keeping only sequences with at least 10, 15, 20, 30, 40 or 50 reads per sampling time point. Then, we randomly split each filtered dataset into training, validation and test sets as described above and made sure that for each split the high-quality training, validation and test sets were contained in the lower quality training, validation and test sets, respectively. Moreover, a test set was held out for the following prediction experiments. In order to identify an optimal lower read count threshold, we trained a single ResNet model for 150 epochs. We randomized the search for hyperparameters^19^ (see above) used the same 150 sets of hyperparameters for each filtered training dataset and determined the coefficient of determination on the validation set. Hence, the minimal threshold was effectively treated as a hyperparameter. This analysis indicated that a minimal threshold of 20 reads per time point was optimal for predictive performance, which saturated for lower thresholds despite the increase in overall dataset size (Supplementary Fig. 9a). We kept this training/validation/test split (“Split0”) for the following prediction experiments. Finally, we confirmed that these conclusions were not an artefact of the random split of the original dataset by repeating this analysis using five different training, validation and test set splits (Supplementary Fig. 9b).

### Evaluation and Benchmarking of the Prediction Model

Using “Split0”, we evaluated our model in more detail. Importantly, this implies that the test set had not been used in previous experiments in order to avoid overfitting. First, we used random search for selecting the best combination among 150 sets of hyperparameters on the validation set (see above), let SAPIENs run for 300 epochs and used an early stopping criterion on the validation set to avoid overfitting by selecting the epoch with the best validation R^2^ (Fig. 4c). To compare our single ResNet models and SAPIENs to different available ML approaches, we trained different models (Fig. 4b,e) on the training set and tuned their hyperparameters by optimizing predictive performance on a subset of the validation set of “Split0”. The single ResNet and SAPIENs models were trained for a maximum of 150 epochs, using early stopping. A total of 100 randomly generated model architectures with 1-3 residual blocks were considered. Hyperparameters tuned for the other models were regularization strength for Ridge Regression^20^, number of neighbours K for k-Nearest Neighbours^21^, number of trees for Random Forests^22^, and maximum depth and learning rate for Gradient Tree Boosting^23^, the later also benefited from early stopping in the validation set. The impact of the training set size on predictive performance (Fig. 4e) was evaluated by training the different models on different smaller datasets, while ensuring that the training and validation sets were contained in the training and validation sets of higher sample size experiments (i.e. nested training and validation sets). Hyperparameters for all models were optimized independently for each training set size on the corresponding validation set. The effect of adding designed sub-libraries to increase the fraction of stronger RBSs in the bulk library (Fig. 3e) was further analysed to evaluate a potential gain in predictive performance for the intermediate and strong sequences (Supplementary Fig. 14). To this end, we performed cross-analyses with the fully degenerate sub-library (N_17_) and the bulk library (N_17_+High1-3). We trained on N_17_ and predicted on unseen subsets of N_17_ and N_17_+High1-3, and trained N_17_+High1-3 and predicted on unseen subsets of N_17_ and N_17_+High1-3 (Supplementary Fig. 14a). In another set of analyses, we omitted each of the enriched sub-libraries while training by moving them to the test sets and evaluated the corresponding effect (Supplementary Fig. 14b). We trained a single ResNet model for 300 epochs for computational considerations and we used early stopping in the validation set. Hyperparameters were tuned independently for each dataset and selected from 150 random configurations in the corresponding validation set. All analyses were done with the same training and validation set sizes. Comparative analyses were performed with the same test set.

### Evaluation of Sequence Motifs and Model Interpretations

We analysed the fully degenerate sub-library (N_17_) in order to measure the impact of the position of known motifs of influence on the RBS activity, such as start-codons (AUG, UUG, GUG) or the consensus Shine-Dalgarno sequence (AGGAGG and subsequences). To this end, for each position, for each group of RBSs that presented the motif of interest at the given position, we calculated simple statistics (median, interquartile ranges, 20/80 percentiles) on the target IFP_0-480min_ of the sequences in the group (Fig. 5a,b and Supplementary Fig. 17). We excluded from these groups RBSs that contained at least one start codon other than the one at the position of interest. We also analysed the filters of the first convolutional layer (excluding the first skip connection) of a ResNet model of the ensemble chosen at random (Fig. 5c). To this end, the effect of each filter was evaluated by calculating Pearson’s correlation coefficient between the filter activations at each position and the flipping integral for all sequences in the test set. Consequently, each filter is represented by a vector of correlations of size 17, which corresponds to the number of positions at which the filter influence is estimated. Finally, the filter representations are then clustered in twelve groups with a complete linkage clustering method using Hamming distance as the underlying metric for comparing individual sequences in order to group filters of similar influence. The integrated gradients^24^ method assigns attribution scores to each base and position by computing the linear path integral between the sequence of interest and a baseline sequence chosen *a priori*. The attribution scores measure the effect of individual bases on the predicted IFP_0-480min_, relative to a baseline. We applied the integrated gradients method to SAPIENs and chose a “blank” one-hot encoded sequence as a neutral baseline (i.e. an all-zeros array). We first used a dimension reduction method, the t-distributed stochastic neighbour embedding (tSNE) method, to visualize how sequences behave in a low dimensional space (perplexity=12, early exaggeration=30) (Fig. 5d). We also averaged the attribution scores of all sequences in the test set, per base and per position, to get a better understanding of the important positions and bases, which contribute either to a high RBS activity or to a low one (Fig. 5e). Finally, in order to account for non-linearities between positions and to understand the drivers of very strong or very weak sequences, we selected the top 5% and the bottom 5% sequences in the test set after removing outlier sequences and clustered each pool with k-means according to their attribution score profiles into five clusters. The medoids of these five clusters are displayed for the strong (Fig. 5f) and weak RBSs (Supplementary Fig. 18). For *in silico* evolution, we selected the weakest (respectively strongest) sequence in the test set and aimed to mutate it progressively to a sequence presenting a maximum (respectively minimum) attainable RBS activity as predicted by SAPIENs (Fig. 5g,h). To do so, we considered all sequences that could result from applying one or two mutations to the current sequence and kept the strongest (respectively weakest) one in each round until no candidate exhibited a change in predicted IFP_0-480min_ in the desired direction.

