## Supplementary material for "Large-scale DNA-based phenotypic recording and deep learning enable highly accurate sequence-function mapping": Methods

RBS $i$ is represented by its approximation $\hat{p}^i = (\hat{p}^i{}_0, \hat{p}^i{}_5, \ldots, \hat{p}^i{}_{720})$. Afterwards, optimal sampling times were greedily selected while fixing the first and last sample at 0 min and 720 min after induction. The goal of the optimization was to find a set $S$ of optimal sampling times (initially $S = (0, 720)$) which allows to reconstruct $\hat{p}^i$ such that a linear approximation using time points in S is as close to $\hat{p}^i$ as possible. Given the set of possible sampling times $T = \{0, 5, 10, \ldots, 720\}$ and the subset $I$ of RBS profiles on which the sampling times should be inferred, the greedy optimization finds the next optimal sampling time point $s^*$ from $T$ as follows:

$$s^* = argmin_{s \in T \setminus S} \frac{1}{|T|} \sum_{i \in I} \sum_{t \in T \cup \{s\}} \left| \hat{p}_t^i - \hat{l}_t^{i,S} \right| \ , \qquad (1)$$

where $\hat{l}^{i,S}$ corresponds to the linear approximation of $\hat{p}^i$ using only sampling times in $S$ (Supplementary Fig. 25b). In other words, $s^*$ is the time point that (i) is not part of the sampling schedule $S$ yet and (ii) results in the smallest cumulative reconstruction error over all RBS profiles in $I$. Subsequently, $S$ is augmented by $s^*$, and equation (1) is evaluated to find the next optimal $s^*$, until $S$ contains the desired number of sampling times. Finally, the quality of the optimal sampling schedule for every RBS $i$ is evaluated by computing the approximation error $r^i$ between the observed profile $p^i$, and its linear interpolation at the optimal time points $S$, termed $l^{i,S}$ (Supplementary Fig. 25c):

*References (Methods)*


1 Sambrook, J. F. & Russell, D. W. *Molecular cloning: a laboratory manual* (Cold Spring Harbor Laboratory, 3rd edition, 2001).

2 Jeschek, M. *et al.* Biotin-independent strains of Escherichia coli for enhanced streptavidin production. *Metab. Eng.* **40**, 33-40 (2017).

3 Martinez-Garcia, E., Aparicio, T., Goni-Moreno, A., Fraile, S. & de Lorenzo, V. SEVA 2.0: an update of the Standard European Vector Architecture for de-/re-construction of bacterial functionalities. *Nucleic Acids Res.* **43**, D1183-D1189 (2015).

4 Datsenko, K. A. & Wanner, B. L. One-step inactivation of chromosomal genes in Escherichia coli K-12 using PCR products. *Proc. Natl. Acad. Sci. U.S.A.* **97**, 6640-6645 (2000).

5 Hastie, T., Tibshirani, R. & Friedman, J. H. *The elements of statistical learning: data mining, inference, and prediction (Springer, New York, 2001).*

6 Salis, H. M., Mirsky, E. A. & Voigt, C. A. Automated design of synthetic ribosome binding sites to control protein expression. *Nat. Biotechnol.* **27**, 946-950 (2009).

7 Perez-Cruz, F. Estimation of information theoretic measures for continuous random variables. *Advances in Neural Information Processing Systems*, 1257-1264 (2009).

8 LeCun, Y. *et al.* Backpropagation applied to handwritten zip code recognition. *Neural Comput.* **1**, 541-551 (1989).

9 Alipanahi, B., Delong, A., Weirauch, M. T. & Frey, B. J. Predicting the sequence specificities of DNA- and RNA-binding proteins by deep learning. *Nat. Biotechnol.* **33**, 831-838 (2015).

10 Zeng, H., Edwards, M. D., Liu, G. & Gifford, D. K. Convolutional neural network architectures for predicting DNA-protein binding. *Bioinformatics* **32**, i121-i127 (2016).

11 Kelley, D. R. *et al.* Sequential regulatory activity prediction across chromosomes with convolutional neural networks. *Genome Res.* **28**, 739-750 (2018).

12 He, K. M., Zhang, X. Y., Ren, S. Q. & Sun, J. Deep residual learning for image recognition. *Proceedings of the IEEE conference on computer vision and pattern recognition*, 770-778 (2016).

13 Xie, S., Girshick, R., Dollár, P., Tu, Z. & He, K. Aggregated residual transformations for deep neural networks. *Proceedings of the IEEE conference on computer vision and pattern recognition*, 1492-1500 (2017).

14 Lakshminarayanan, B., Pritzel, A. & Blundell, C. Simple and scalable predictive uncertainty estimation using deep ensembles. *Advances in Neural Information Processing Systems* **30**, 6402-6413 (2017).

15 Ioffe, S. S., C. Batch normalization: accelerating deep network training by reducing internal covariate shift. *Proceedings of the 32nd International Conference on Machine Learning* **37**, 448-456 (2015).

16 Maas, A. L., Awni Y. Hannun, and Andrew Y. Ng. Rectifier nonlinearities improve neural network acoustic models. *Proceedings of the 30thInternational Conference on Machine Learning* **28** (2013).

17 Kingma, D. P. a. B., J. ADAM: a method for stochastic optimization. *ICLR* (2015).

18 Abadi, M. *et al.* TensorFlow: a system for large-scale machine learning. *Proceedings of the 12th USENIX Symposium on Operating Systems Design and Implementation* (2016).

19 Bergstra, J. & Bengio, Y. Random search for hyper-parameter optimization. *Journal of Machine Learning Research*, 281-305 (2012).

20 Hastie, T., Tibshirani, R. & Friedman, J. H. *The elements of statistical learning: data mining, inference, and prediction (Springer, New York, 2001).*

21 Altman, N. S. An introduction to kernel and nearest-neighbor nonparametric regression. A*merican Statistician* **46**, 175–185 (1992).

22 Breiman, L. Random forests. *Mach. Learn.* **45**, 5-32 (2001).

23 Friedman, J. H. Greedy function approximation: a gradient boosting machine. *Ann. Stat.* **29**, 1189-1232 (2001).

24 Sundararajan, M., Taly, A. & Yan, Q. Axiomatic attribution for deep networks. *Proceedings of the 34th International Conference on Machine Learning* **70,** 3319-3328 (2017).
