## Supplementary Information for "Large-scale DNA-based phenotypic recording and deep learning enable highly accurate sequence-function mapping"

† Equal contribution

### Supplementary Figures

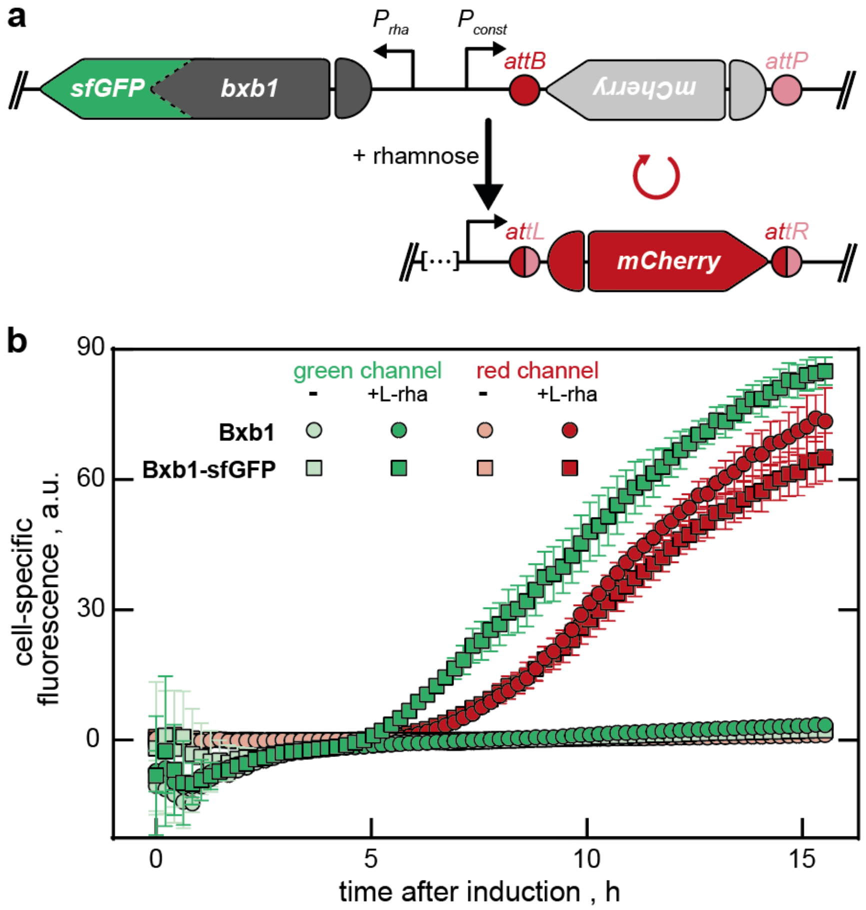


**Supplementary Fig. 1 | Recombination activity of the Bxb1-sfGFP fusion. A**, In plasmid pASPIre2, a Bxb1-sfGFP fusion controlled by the rhamnose-inducible promoter $P_{rha}$ is used as modifier to invert an mCherry CDS into the right orientation to a constitutive promoter $P_{const}$ to activate *mCherry* expression. *attB*/*P* and *attL*/*R*: Bxb1 attachment sites before and after recombination. **b**, The Bxb1-sfGFP fusion exhibits similar recombination activity as the GFP-less variant Bxb1 as indicated by the increase in red fluorescence after induction with 0.2% rhamnose (+L-rha). The appearance of the fusion protein can be tracked relying on sfGFP fluorescence. Microtiter plate cultivation of *E. coli* TOP10 Δ*rhaA*. Data points represent mean values of four independent culture wells with error bars indicating standard deviation.

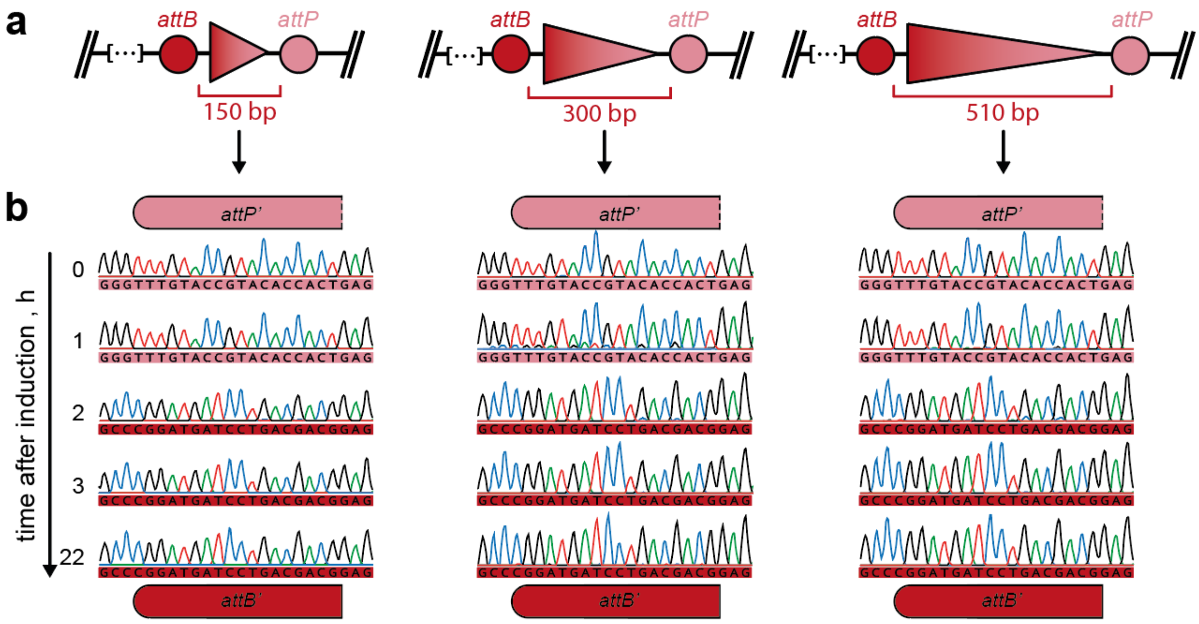


**Supplementary Fig. 2 | Evaluation of silent discriminators with different spacer lengths. a**, DNA spacers of different lengths are introduced between Bxb1 attachment sites *attB* and *attP* (plasmids pASPIre3, pASPIre4 and pASPIre5 corresponding to spacer lengths of 150 bp, 300 bp and 510 bp, respectively). **b**, Bxb1-mediated discriminator inversion in shake flask cultivations of *E. coli* TOP10 as analyzed by Sanger sequencing. The displayed sequence window corresponds to the left half site of *attB* and *attR* before and after recombination, respectively. Since no significant impact on the kinetics of BxB1 recombination was observed in the tested range of spacer lengths the shortest discriminator (150 bp spacer, pASPIre3) was selected for all following experiments.

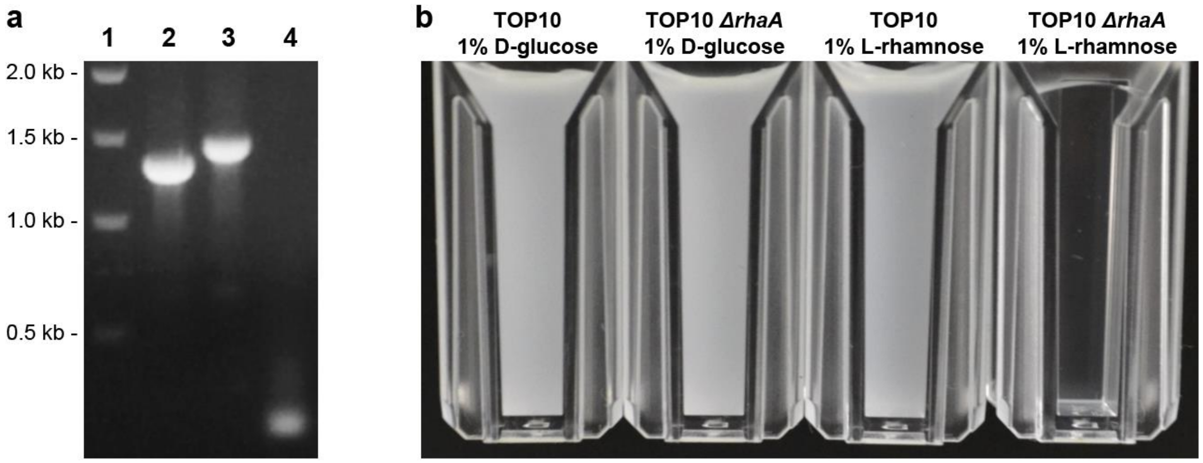


**Supplementary Fig. 3 | Construction of a rhamnose utilization-deficient strain of *E. coli*. a**, Knockout of the *rhaA* gene was verified by analytical PCR of the genomic locus using primers 10 and 11. Lane 1: DNA ladder, lane 2: *E. coli* TOP10 (parent strain), lane 3: *E. coli* TOP10 *rhaA::kan*$^R$, lane 4: *E. coli* TOP10 Δ*rhaA* (i.e. strain after removal of the *kan*$^R$ cassette). **b**, Deficiency in rhamnose utilization was confirmed by testing growth of *E. coli* TOP10 Δ*rhaA* in defined mineral medium supplemented with 1% of l-rhamnose as sole carbon source.

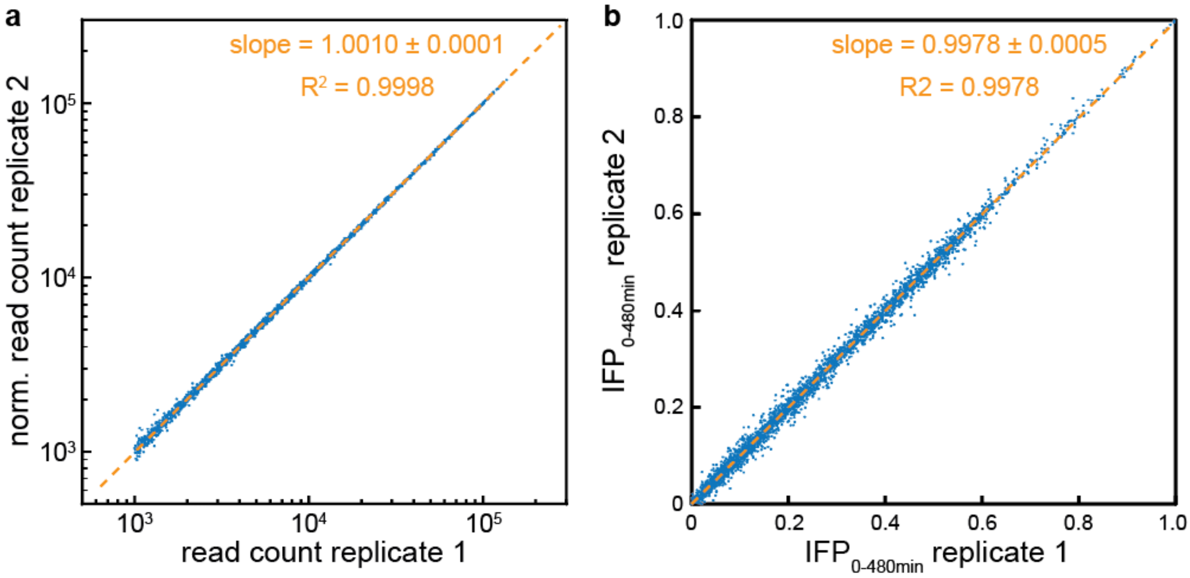


**Supplementary Fig. 4 | Technical reproducibility of uASPIre.** Data points represent the read count (**a**) and the integral of the flipping profiles between 0 and 480 minutes after induction ($IFP_{0\text{-}480min}$, **b**) of each RBS from the proof-of-concept library as retrieved from two independent NGS runs (technical NGS replicates of the identical sample). For this analysis, a global minimal threshold of 1,000 reads per RBS was applied (10,419 variants above threshold). To facilitate comparison between the two replicates, the read counts for replicate 2 in **a** were normalized by a correction factor that accounts for the different sum of all reads in the runs: $(norm.read\ count\ r2) = \frac{(\text{sum of all reads r1})}{(\text{sum of all reads r2})} \cdot (absolute\ read\ count\ r2)$.

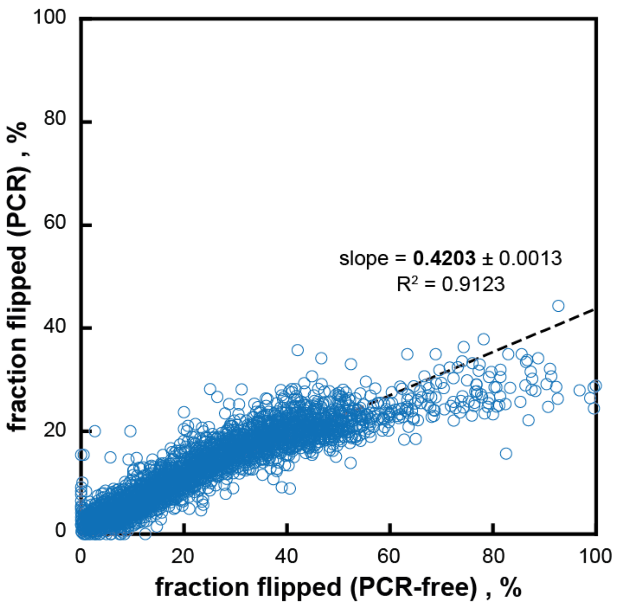


**Supplementary Fig. 5 | Comparison of PCR-based and PCR-free sample preparation.** Percentage of flipped discriminators was determined for a culture sample from the proof-of-concept RBS library (compare Fig. 2c) taken three hours after induction. NGS sample preparation was performed by two different procedures, direct ligation of adapters to the target fragment (“PCR-free”, horizontal axis) and PCR-amplification of target fragments (“PCR”, vertical axis). PCR-amplification introduces a strong, non-linear bias in favor of unflipped discriminators which also leads to a significant reduction of the apparent dynamic measurement window. Details of the different sample preparation procedures can be found in the Methods section.

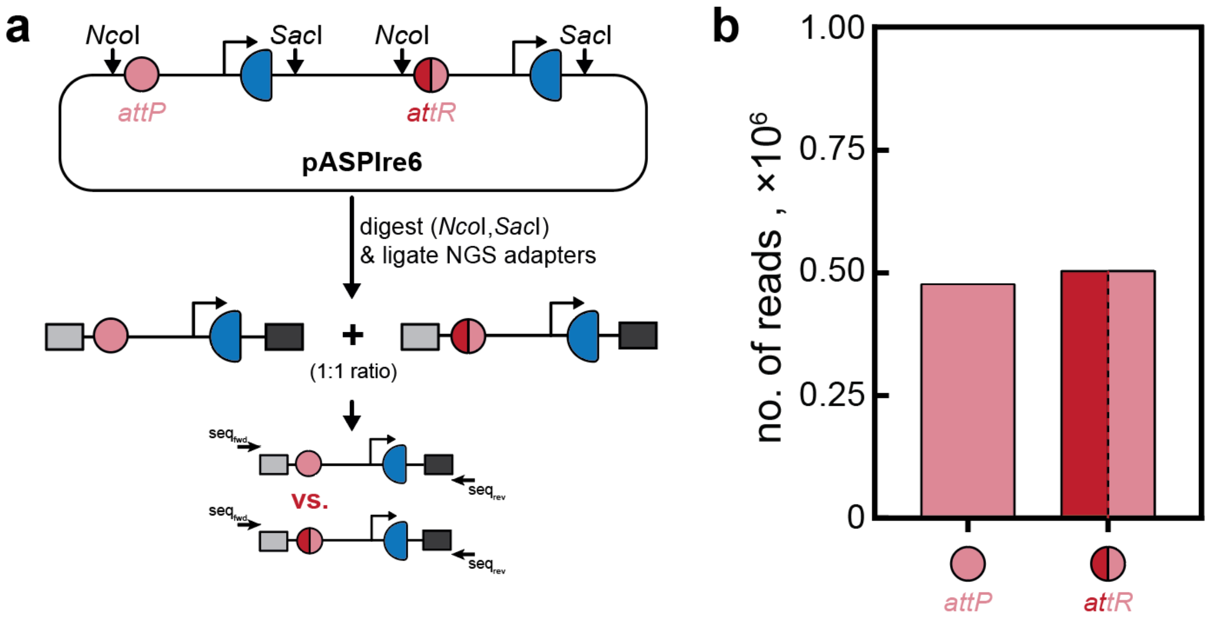


**Supplementary Fig. 6 | Evaluation of bias introduced during PCR-free sample preparation. a**, Plasmid pASPIre6 contains a copy of both the unflipped and flipped discriminator from pASPIre3, each of which is flanked by the same restriction sites (*Nco*I and *Sac*I) used during the developed uASPIre workflow. Restriction digest of pASPIre6 should result in a 1:1 ratio of unflipped and flipped discriminator fragments which was analyzed by applying the PCR-free uASPIre sample preparation and subsequent NGS. **b**, Number of read counts for unflipped and flipped discriminators (corresponding to *attP* and *attR* sites, respectively) as obtained from PCR-free preparation of the restriction products of pASPIre6 according to the uASPIre workflow.

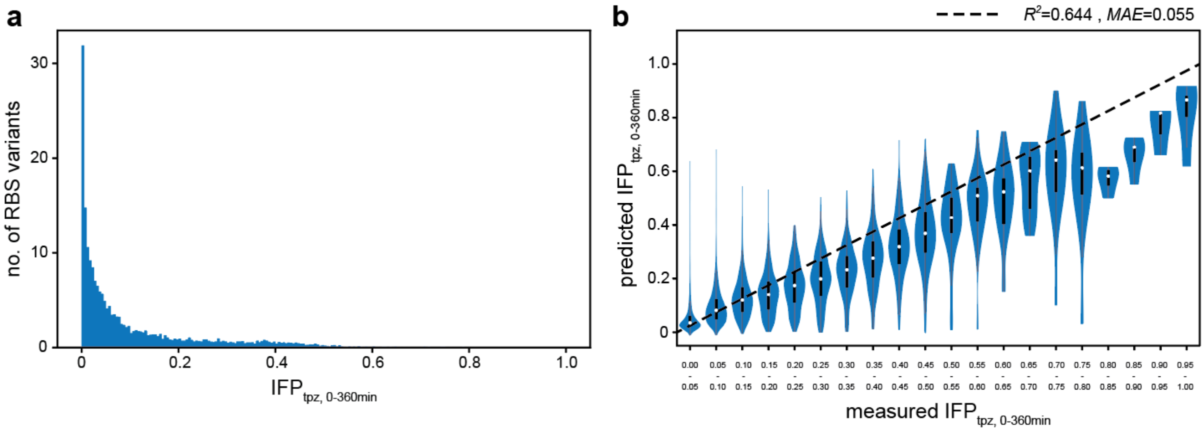


**Supplementary Fig. 7 | Bias in the initial RBS library. a**, The proof-of-concept RBS library with 17 consecutive, fully randomized bases ($N_{17}$) upstream of the Bxb1-sfGFP start codon shows a strong skew towards weak RBSs as represented by the integral of the flipping profile between 0 and 360 minutes after induction approximated using the trapezoidal rule ($IFP_{tpz,\ 0\text{-}360min}$). **b**, Initial predictions obtained with a convolutional neural network model (5-fold cross-validation) trained on the proof-of-concept RBS library. Coefficient of determination ($R^2$) and mean absolute error (MAE) are obtained based on predictions on held-out data.

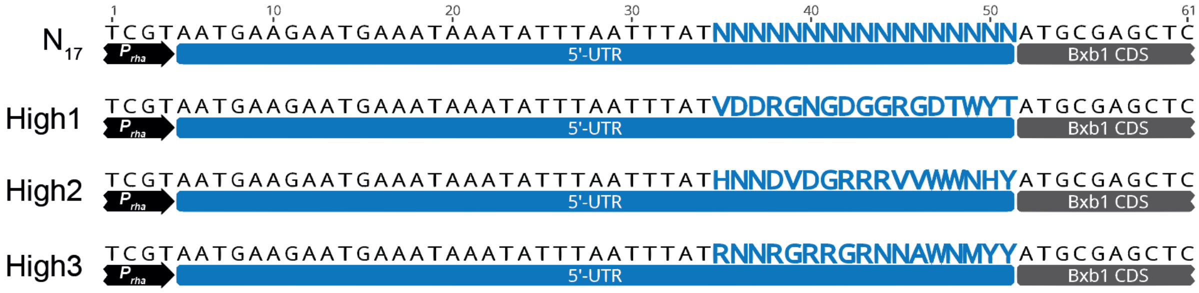


**Supplementary Fig. 8 | RBS libraries constructed in this study.** 17 consecutive base pairs upstream of the *bxb1* start codon were randomized. A fully degenerate library ($N_{17}$) as well as three libraries with reduced skew towards weak RBSs (High1, High2, High3) were constructed. Details about library design and cloning can be found in the Methods section. A map of plasmid pASPIre3, which served as a starting point to construct the displayed libraries, is shown in Supplementary Figure 21.

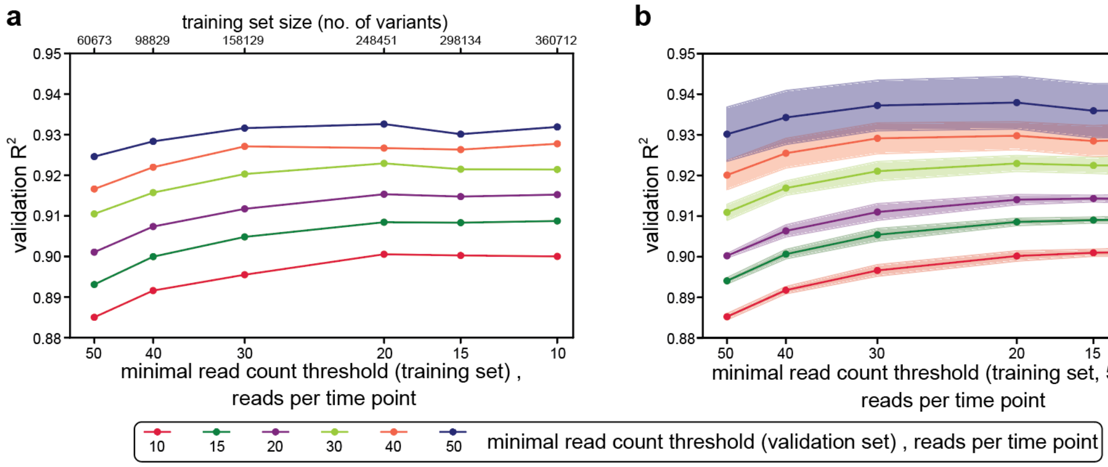


**Supplementary Fig. 9 | Effects of the minimal read threshold on the predictive performance of the ResNet. a**, ResNet models were trained on different subsets of the training data corresponding to different minimal read count thresholds, and were evaluated in terms of the coefficient of determination $R^2$ in different subsets of the validation set also corresponding to different minimal read count thresholds. The data training/validation/test split ("Split0") corresponds to the one we use to obtain the results in Figure 4. **b**, The experiment shown in (**a**) was repeated four times for different random splits to assess robustness. Data points in (**b**) represent the average of five random splits with two-standard-deviation intervals shown as shaded areas. A detailed description of the threshold is provided in the Methods section.

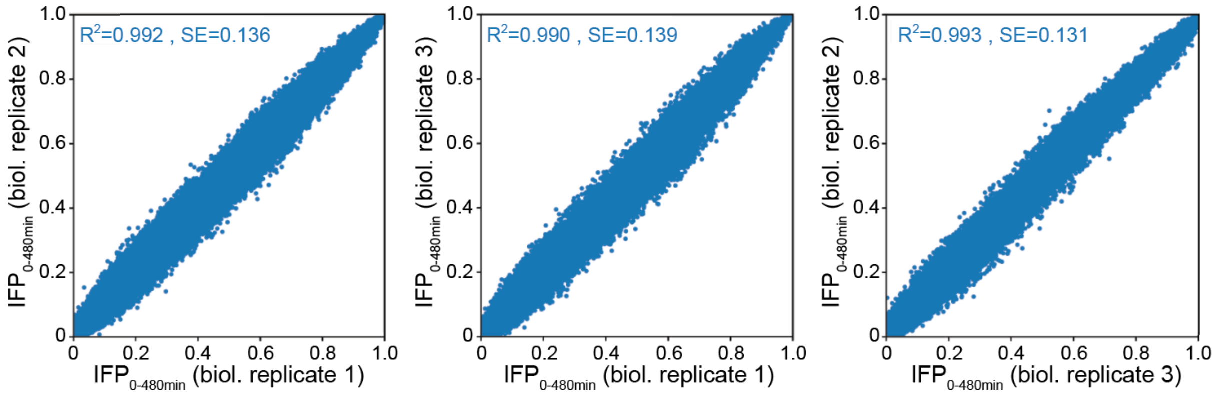


**Supplementary Fig. 10 | Biological replicates.** A total of three independent biological replicates (i.e. individual shake flask cultivations) of the second, larger RBS library were subjected to the uASPIre workflow. Relying on the spiked-in 31 internal-standard RBSs, a normalization curve was constructed and used to normalize the $IFP_{0\text{-}480min}$ values between replicates (Methods). SE: standard error.

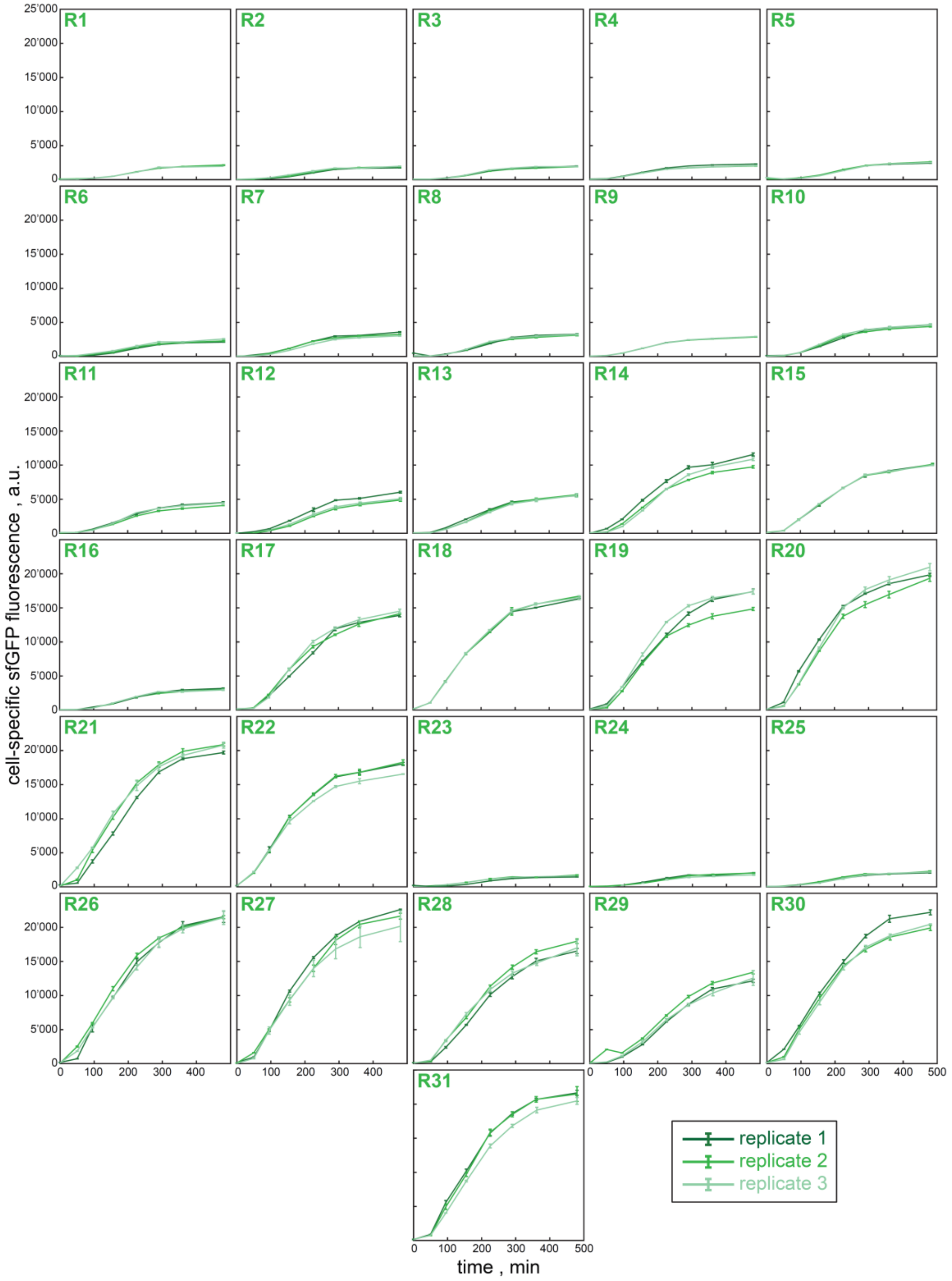


**Supplementary Fig. 11 | Time-resolved Bxb1-sfGFP production for internal-standard RBSs R1-R31.** For each internal-standard RBS cell-specific fluorescence was recorded in three independent shake flask cultivations (biological replicates). Data points are depicted as a function of the time after induction and represent the average of three technical replicates with standard deviation indicated by error bars.

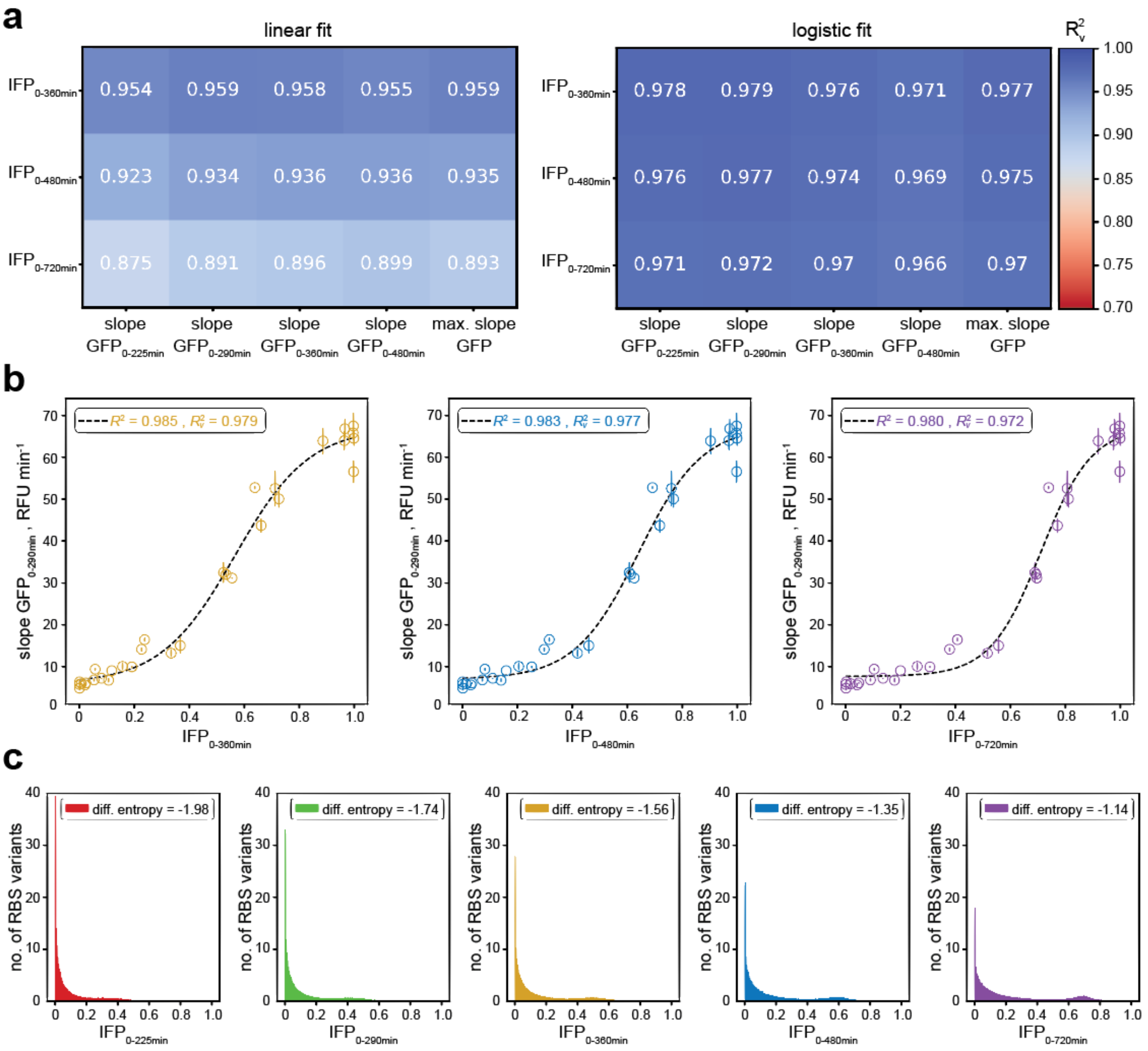


**Supplementary Fig. 12 | Identification of optimal parameters to correlate Bxb1-mediated recombination with cellular Bxb1-sfGFP levels. a**, Coefficient of determination after leave-one-out cross validation (loo-CV) ($R_v^2$) between different slope- and integral-based summary statistics for cell-specific fluorescence and the flipping profiles of the 31 internal-standard RBSs using linear and logistic fits. Note that slope-based summary statistics for the flipping profiles failed to deliver robust fits ($R_v^2$ consistently below 0.5) and were therefore not included in this figure. **b**, Selected logistic fits involving the integral of the flipping profile (IFP) for different time spans and the slope of the cell-specific fluorescence curve between 0 and 290 min after induction. The standard deviation of three biological replicates for the fluorescence profiles is indicated by vertical error bars and coefficients of determination without ($R^2$) and with loo-CV ($R_v^2$) are displayed. **c**, IFP distribution across the entire larger RBS library for different integration intervals. The differential entropies of the respective IFP probability densities are indicated. $IFP_{0\text{-}t\ min}$: normalized integral of the flipping profile between 0 and *t* min after induction; slope $GFP_{0\text{-}t\ min}$: slope of the cell-specific fluorescence curve between 0 and *t* min after induction; max. slope GFP: maximum slope (minimum three timepoints) of the cell-specific fluorescence curve.

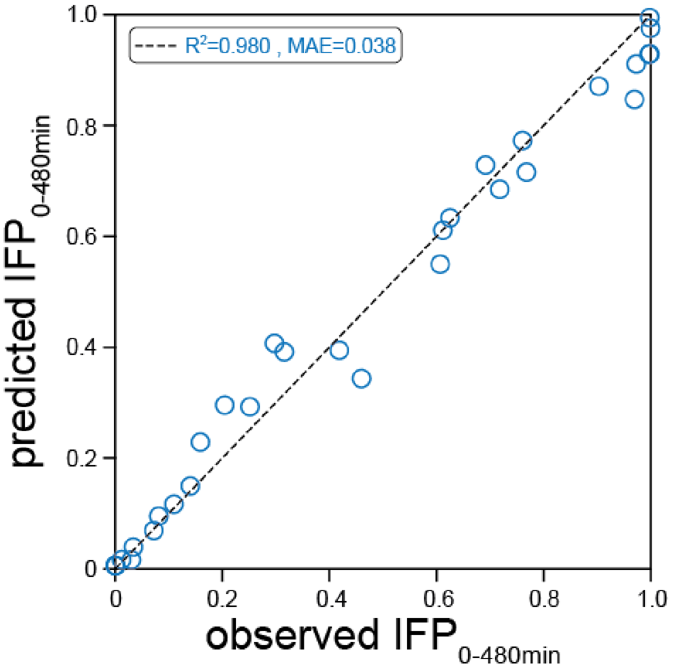


**Supplementary Fig. 13 | Correlation of predicted and observed $IFP_{0\text{-}480min}$ for the 31 internal-standard RBS used in this study.** The $IFP_{0\text{-}480min}$ values for the 31 internal-standard RBSs as predicted by SAPIENs are highly correlated with the corresponding values experimentally determined by uASPIre, which were held out during training.

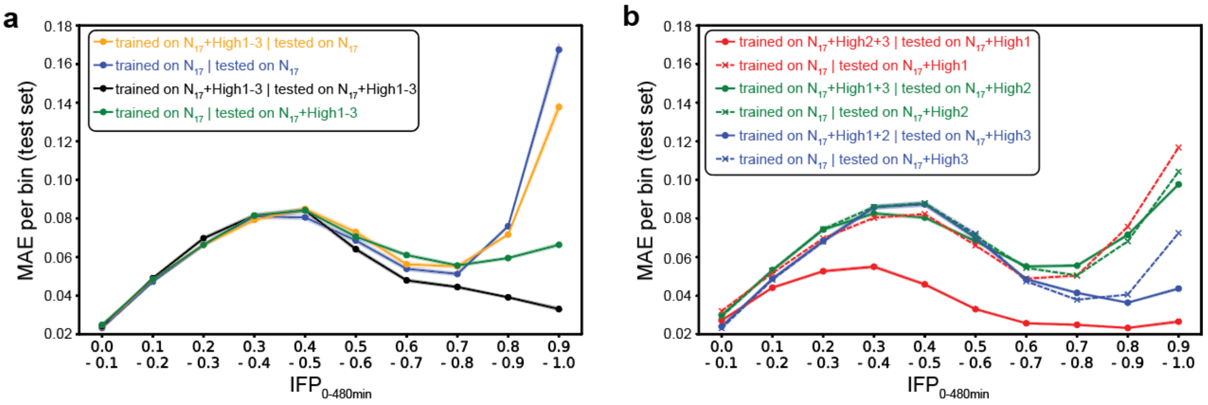


**Supplementary Fig. 14 | Effects of designed sub-libraries on the prediction accuracy of the ResNet model. a/b**, The mean absolute error (MAE) is evaluated for different bins of the experimentally determined $IFP_{0\text{-}480min}$ value and several combinations of training and test sets composed of the fully degenerate RBS library ($N_{17}$) and the designed libraries (High1-3). 95% confidence intervals are indicated by shaded areas.

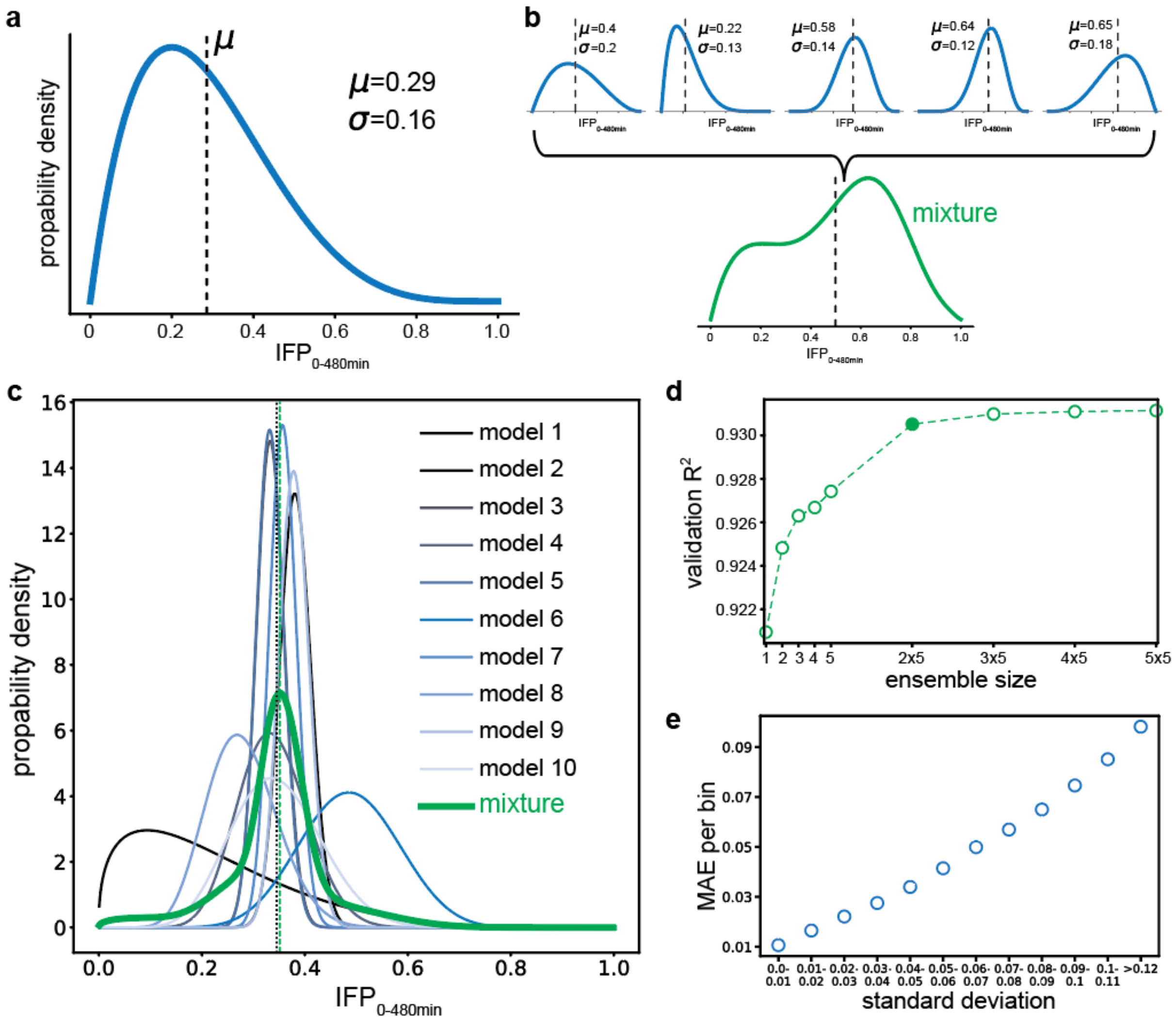


**Supplementary Fig. 15 | Uncertainty estimation of the prediction. a**, Each single ResNet models IFP$_{0\text{-}480min}$ for each RBS sequence as a beta distribution whose mean μ and standard deviation σ can be computed. While μ corresponds to the predicted IFP$_{0\text{-}480min}$ value, σ represents a measure of the uncertainty of the prediction. **b**, Combining multiple ResNet models into an ensemble allows modeling IFP$_{0\text{-}480min}$ as a mixture of beta distributions, for each RBS sequence. **c**, Example of the predicted mixture of beta distributions (in green) of an ensemble of ten models for a given RBS sequence. The ensemble achieves a better prediction than the individual models as can be appreciated from its mean μ (green dashed line) which is in close proximity to the experimentally determined ground truth IFP$_{0\text{-}480min}$ value (black dotted line). **d**, The validation coefficient of determination $R^2$ increases with the number of ensemble members. An ensemble size of 2x5 (filled circle), corresponding to two hyperparameter configurations with five independently trained ResNets per configuration, was selected as a trade-off between acuracy and computational demand. **e**, The per-bin mean absolute error (MAE) is strongly correlated with the predicted uncertainty as measured by the predicted standard deviation for the 2x5 ResNet ensemble.

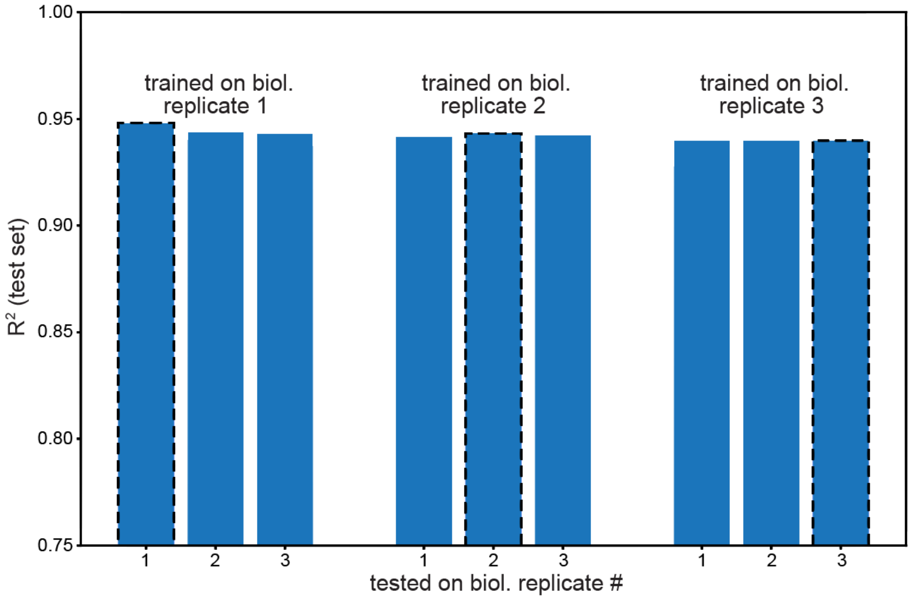


**Supplementary Fig. 16 | Cross-training between biological replicates.** Three instances of the model are trained and validated independently on each biological replicate. Afterwards, they are tested on test labels from the same (within replicate performance, outlined columns) and the other two replicates (cross-replicate performance). For cross-replicate analysis data from the other two replicates were normalized using internal-standard RBSs (Supplementary Fig. 10, Methods). Further details about the cross-training analysis are available in the ML Annex.

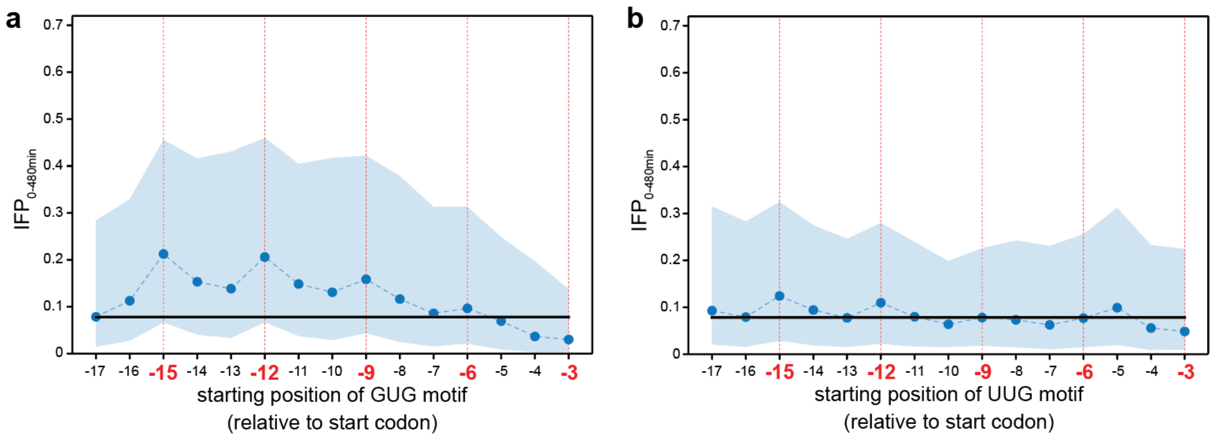


**Supplementary Fig. 17 | Influence of GUG (a) and UUG (b) codons in the 5'-UTR on the RBS strength.** Black horizontal line corresponds to the median $IFP_{0\text{-}480min}$ in the dataset. Circles represent median $IFP_{0\text{-}480min}$ with shaded areas containing percentiles 20/80. In-frame positions are highlighted in red.

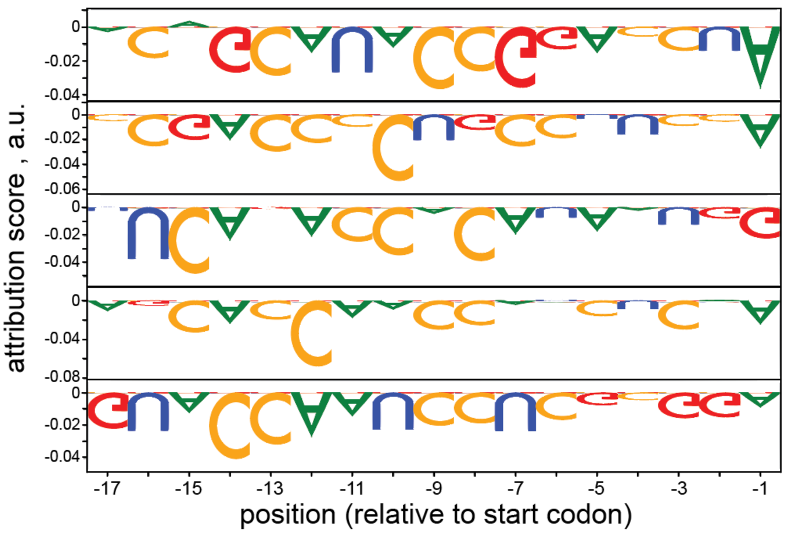


**Supplementary Fig. 18 | Attribution of bases and positions in the 5'-UTR to weak RBSs.** The weakest 5% of RBSs in the test set were distributed into five clusters using k-means algorithm. The displayed motifs are the five medoids of each cluster (i.e. the five individual sequences closest to the respective cluster centroid).

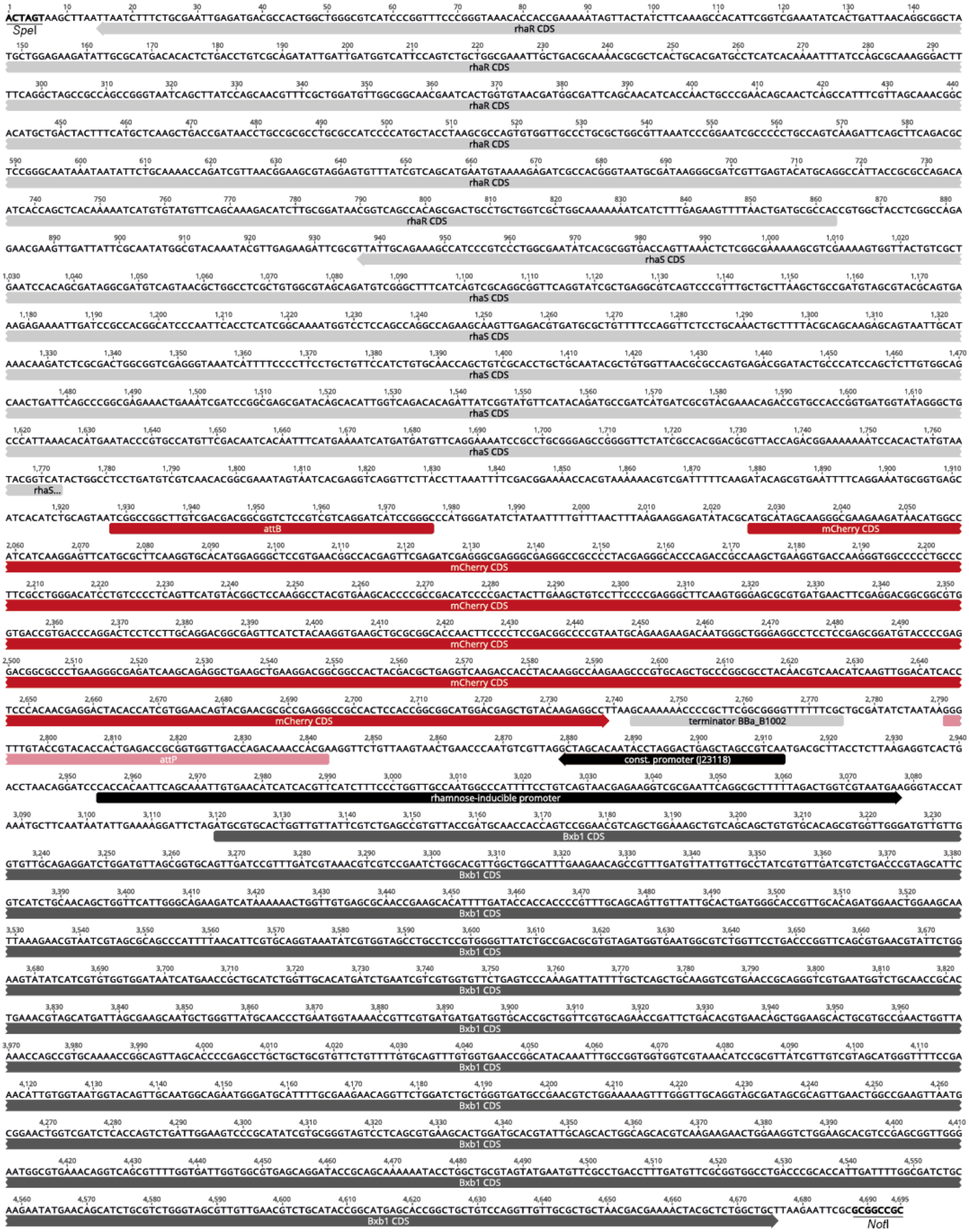


**Supplementary Fig. 19 | Plasmid map of pASPIre1.** The CDS (including 35 bp upstream of the start codon as designed in an earlier study[1] was placed between Bxb1 attachment sites *attB* and *attP*. Upon activation of Bxb1, this cassette is inverted and mCherry is transcribed via a constitutive promoter (J23118, Anderson collection). The CDS of Bxb1 was derived from an earlier study[2]. *rhaR*/*rhaS*: transcriptional activators for the rhamnose-inducible promoter (*E. coli*). For clarity, the plasmid backbone derived from pSEVA291[3] (kanR, pBR322 ori) was omitted. The displayed sequence can be found as plain text below.

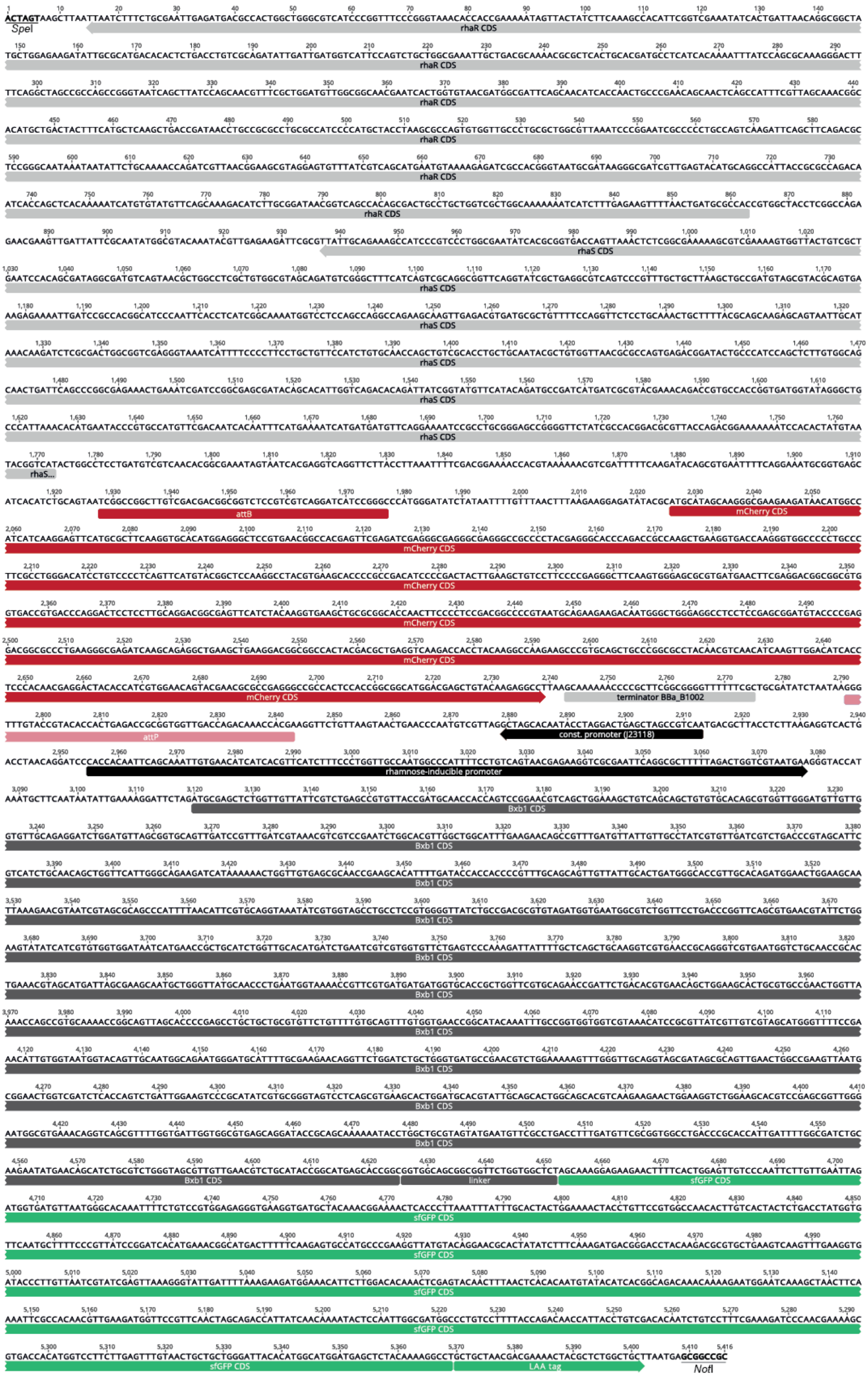


**Supplementary Fig. 20 | Plasmid map of pASPIre2.** This plasmid was derived from pASPIre1 by replacing the *bxb1* gene with a *bxb1-sfGFP* fusion. The CDS of *sfGFP* was derived from a previous study[1]. LAA tag: protein degradation tag. For clarity, the plasmid backbone derived from pSEVA291[3] (*kanR*, pBR322 *ori*) was omitted. The displayed sequence can be found as plain text below.

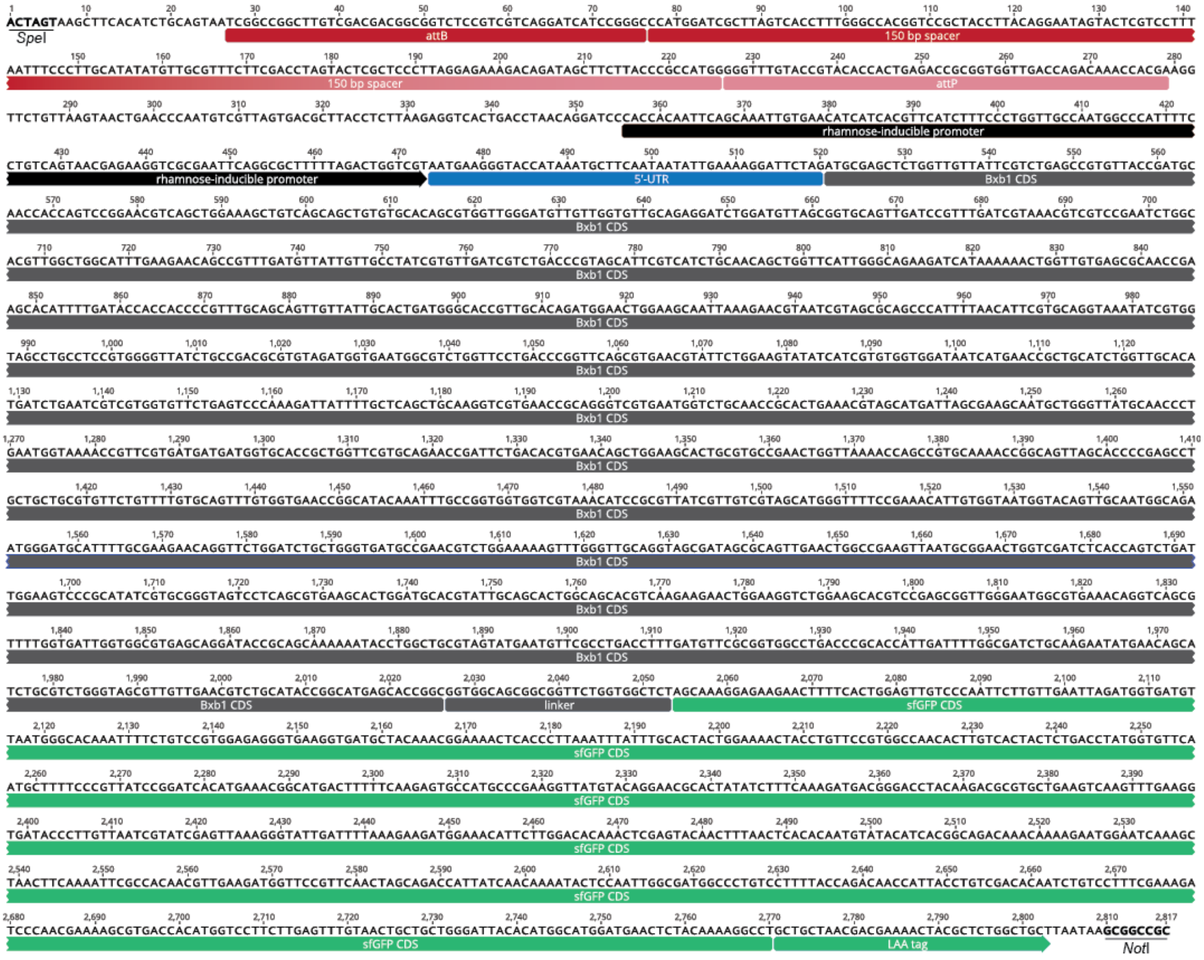


**Supplementary Fig. 21 | Plasmid map of pASPIre3.** This plasmid was derived from pASPIre2 by replacing the mCherry discriminator with a silent 150 bp spacer flanked by Bxb1 attachment sites *attB*/*P*. Plasmids pASPIre4 and pASPIre5 contain 300 bp and 510 bp spacers, respectively (Supplementary Fig. 22), but are otherwise identical with this pASPIre3. For clarity, the plasmid backbone derived from pSEVA291[3] (*kanR*, pBR322 ori) was omitted. The displayed sequence can be found as plain text below.

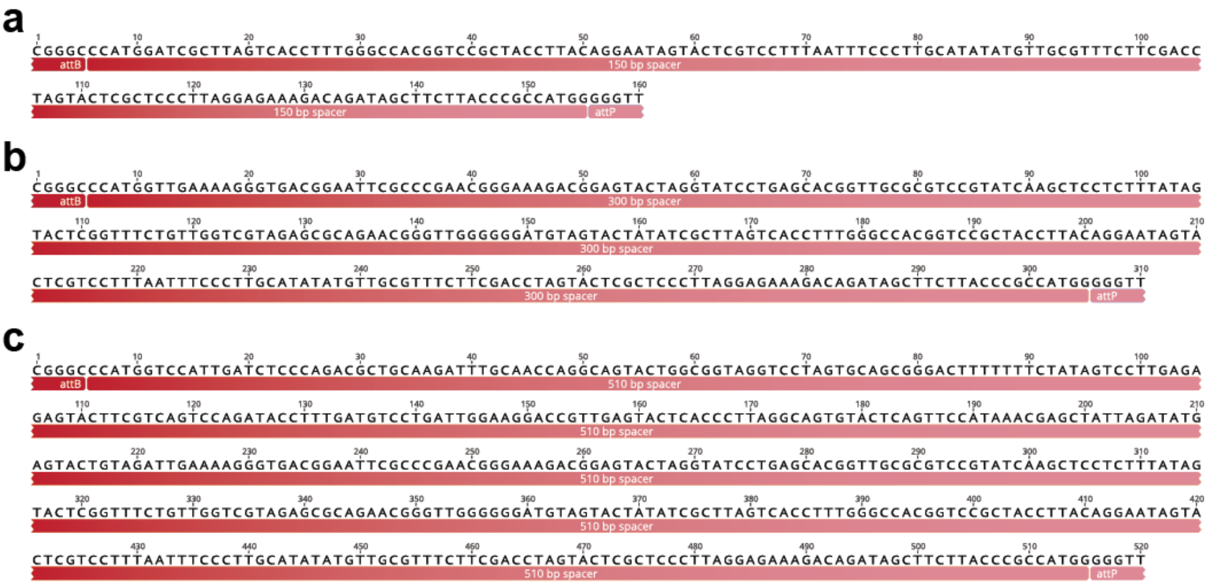


**Supplementary Fig. 22 | Discriminators of plasmids pASPIre3 (a), pASPIre4 (b) and pASPIre5 (c).** Silent (non-coding) DNA spacers were designed using the "Random DNA generator" (Maduro Lab, UC Riverside) at a fixed GC-content of 50%.

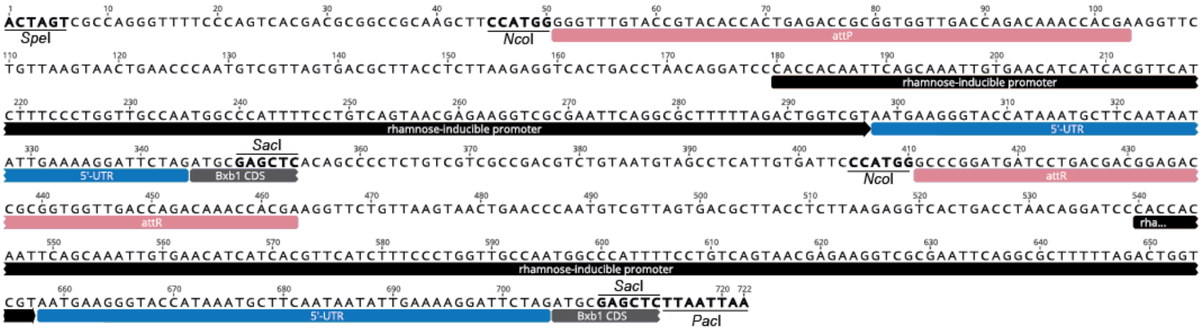


**Supplementary Fig. 23 | Plasmid map of pASPIre6.** This plasmid contains a copy of both the unflipped and flipped discriminator from pASPIre3, each of which is flanked by the same restriction sites (*Nco*I and *Sac*I) used during the developed uASPIre workflow. For clarity, the plasmid backbone derived from pSEVA291[3] (*kanR*, pBR322 *ori*) was omitted. The displayed sequence can be found as plain text below.

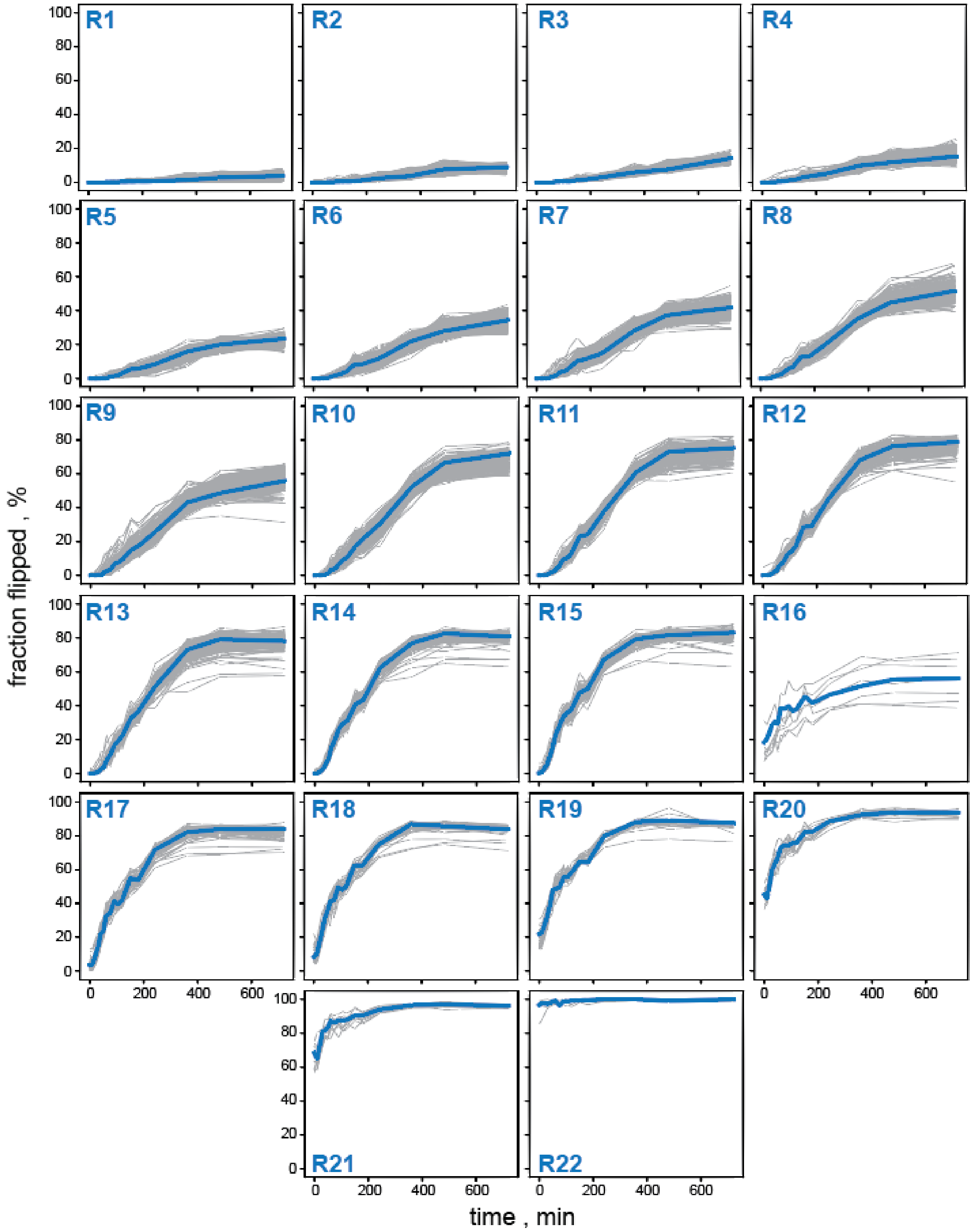


**Supplementary Fig. 24 | Clustering of RBSs from the proof-of-concept library.** RBSs were clustered according to the observed behavior in Bxb1-mediated discriminator flipping as described in the Methods section. The RBS in the center of each cluster (R1-R22, highlighted in blue) was selected as representative and used as internal-standard RBS to record calibration curves. Sequences corresponding to the cluster representatives are listed in Supplementary Table 4.

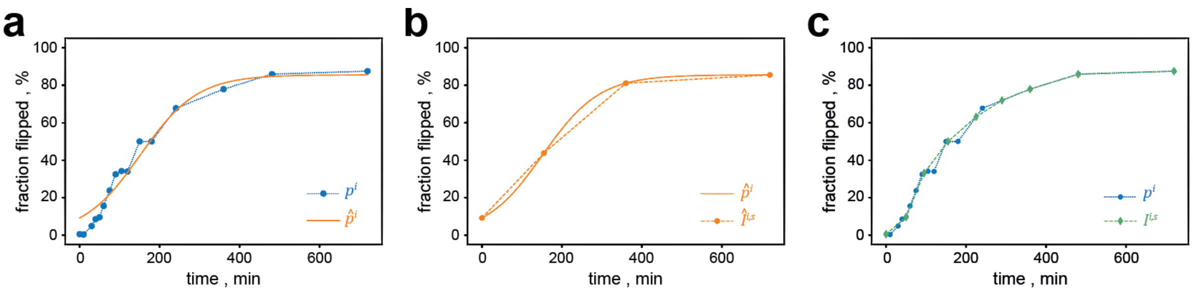


**Supplementary Fig. 25 | Schematic representation of approximations and errors used during optimization of the sampling schedule. a**, The measured kinetic profile $p^i$ of RBS $i$ is represented by a corresponding approximation $\hat{p}^i$. **b**, Approximation $\hat{p}^i$ is reduced to four samples S=(0,155,360,720) and the corresponding linear interpolation $\hat{l}^{i,S}$. Sampling points $s^*$ are consecutively chosen to minimize the cumulative reconstruction error (i.e. the area between the two curves $\hat{p}^i$ and $\hat{l}^{i,S}$) averaged over all RBSs $i \in I$. **c**, Comparison of measured kinetic profile $p^i$ and the corresponding profile after optimization, $l^{i,S}$. The approximation error $r^i$ is computed as in equation (2) and constitutes an approximation to the area between the two curves. A detailed description of the optimization is provided in the Methods section.

### Supplementary Tables

**Supplementary Tab. 1 Primers used in this study.**

| No. | Sequence (5' to 3') |
|---|---|
| 1 | GGCGGTGGGGATTGATGTCGAGGAGGCGCTGCGCCAATTCCGGGGATCCGTCGACC |
| 2 | CCACGCGGCAATGCGGTTGATAGAGGCATCGAAGAAGTGTAGGCTGGAGCTGCTTC |
| 3 | ATCTGCAGTAATCGGCCG |
| 4 | ATAGAGCTCGCATNNNNNNNNNNNNNNNNNATAAATTAAATATTTATTTCATTCTTCATTACGACCAGTCTAAAAAGC |
| 5 | TATATAGAGCTCGCATARWAHCYCCHCNCYHHBATAAATTAAATATTTATTTCATTCTTCATTACGACCAGTCTAAAAAGC |
| 6 | TATATAGAGCTCGCATRDNWWBBYYYCHBHNNDATAAATTAAATATTTATTTCATTCTTCATTACGACCAGTCTAAAAAGC |
| 7 | TATATAGAGCTCGCATRRKNWTNNYCYYCYNNYATAAATTAAATATTTATTTCATTCTTCATTACGACCAGTCTAAAAAGC |
| 8 | AATGATACGGCGACCACCGAGATCTACACTCTTTCCCTACACGACGCTCTTCCGATCTTATCACGACGACGGCGGTCTC |
| 9 | CAAGCAGAAGACGGCATACGAGATGTGACTGGAGTTCAGACGTGTGCTCTTCCGATCTTATCACGATAACAACCAGAGCTCGC |
| 10 | ACGACAGACAAAGGAGCTTTGC |
| 11 | TCGCGCCGCATCCGGCAG |

**Supplementary Tab. 2 | *E. coli* strains and plasmids used in this study.**

| Strain or plasmid | Genotype/Description | Source/Reference |
|---|---|---|
| <u>*E. coli* strains</u> | | |
| TOP10 | $F^-$ *mcrA Δ(mrr-hsdRMS-mcrBC) φ80lacZΔM15 ΔlacX74 nupG recA1 araD139 Δ(ara-leu)7697 galE15 galK16 rpsL*($Str^R$) *endA1* $\lambda^-$; general cloning strain | Thermo Fisher Scientific, Reinach, Switzerland |
| TOP10 *rhaA::kan$^R$* | TOP10 derivative carrying an insertional knockout of the L-rhamnose isomerase gene *rhaA*; deficient in rhamnose utilization | This study |
| TOP10 Δ*rhaA* | Derivative of TOP10 *rhaA::kan$^R$* with kanamycin resistance cassette removed by FRT recombination[4]; deficient in rhamnose utilization | This study |
| <u>Plasmids</u> | | |
| pSEVA291 | Plasmid backbone used for pASPIre plasmid series containing a kanamycin resistance cassette, a pBBR322 replicon and a multiple cloning site | Martinez-Garcia *et al.* (2015)[3] |
| pASPIre1 | Derivative of pSEVA291[3] carrying the *bxb1* gene[2] under the control of a rhamnose-inducible promoter and *mCherry* flanked by *attB*/*P* sites | This study |
| pASPIre2 | Derivative of pASPIre1 carrying a *bxb1-sfGFP* translational fusion in place of *bxb1* | This study |
| pASPIre3 | Derivative of pASPIre2 lacking *mCherry* but containing a silent 150 bp DNA spacer flanked by *attB*/*P* sites; used to create RBS libraries | This study |
| pASPIre4 | Derivative of pASPIre2 lacking *mCherry* but containing a silent 300 bp DNA spacer flanked by *attB*/*P* sites | This study |
| pASPIre5 | Derivative of pASPIre2 lacking *mCherry* but containing a silent 510 bp DNA spacer flanked by *attB*/*P* sites | This study |
| pASPIre6 | Derivative of pSEVA291 containing a copy of both the unflipped and flipped discriminator from pASPIre3 flanked by *Nco*I and *Sac*I sites | This study |
| pKD13 | PCR template for the generation of the linear DNA fragment used to delete the chromosomal *rhaA* gene | Datsenko & Wanner (2000)[4] |
| pASPIre3_R1-R31 | Derivatives of pASPIre3 containing internal-standard RBSs R1-R31 controlling the translation of the Bxb1-sfGFP fusion | This study |

**Supplementary Tab. 3 | Customized duplex DNA adapters for Illumina sequencing used in this study.** Blue: Flow cell binding site, green: Illumina sequencing primer binding site, red: customized index, P: phosphorylation. All duplexes were obtained from Integrated DNA Technologies (Leuven, Belgium).

<table>
<tr><th>No.</th><th>Sequence</th></tr>
<tr><td>Left1</td><td>5′-AATGATACGGCGACCACCGAGATCTACACTCTTTCCCTACACGACGCTCTTCCGATCTTATCACGC-3′<br>3′-TTACTATGCCGCTGGTGGCTCTAGATGTGAGAAAGGGATGTGCTGCGAGAAGGCTAGAATAGTGCGGTAC-P-5′</td></tr>
<tr><td>Left2</td><td>5′-AATGATACGGCGACCACCGAGATCTACACTCTTTCCCTACACGACGCTCTTCCGATCTATCGATGTC-3′<br>3′-TTACTATGCCGCTGGTGGCTCTAGATGTGAGAAAGGGATGTGCTGCGAGAAGGCTAGATAGCTACAGGTAC-P-5′</td></tr>
<tr><td>Left3</td><td>5′-AATGATACGGCGACCACCGAGATCTACACTCTTTCCCTACACGACGCTCTTCCGATCTGATCTTGTAC-3′<br>3′-TTACTATGCCGCTGGTGGCTCTAGATGTGAGAAAGGGATGTGCTGCGAGAAGGCTAGACTAGAACATGGTAC-P-5′</td></tr>
<tr><td>Left4</td><td>5′-AATGATACGGCGACCACCGAGATCTACACTCTTTCCCTACACGACGCTCTTCCGATCTCGATGCCAATC-3′<br>3′-TTACTATGCCGCTGGTGGCTCTAGATGTGAGAAAGGGATGTGCTGCGAGAAGGCTAGAGCTACGGTTAGGTAC-P-5′</td></tr>
<tr><td>Left5</td><td>5′-AATGATACGGCGACCACCGAGATCTACACTCTTTCCCTACACGACGCTCTTCCGATCTTCGATACAGTGC-3′<br>3′-TTACTATGCCGCTGGTGGCTCTAGATGTGAGAAAGGGATGTGCTGCGAGAAGGCTAGAAGCTATGTCACGGTAC-P-5′</td></tr>
<tr><td>Left6</td><td>5′-AATGATACGGCGACCACCGAGATCTACACTCTTTCCCTACACGACGCTCTTCCGATCTATCGATACTTGAC-3′<br>3′-TTACTATGCCGCTGGTGGCTCTAGATGTGAGAAAGGGATGTGCTGCGAGAAGGCTAGATAGCTATGAACTGGTAC-P-5′</td></tr>
<tr><td>Right1</td><td>5′-P-CCGTGATAAGATCGGAAGAGCACACGTCTGAACTCCAGTCACATCTCGTATGCCGTCTTCTGCTTG-3′<br>3′-TCGAGGCACTATTCTAGCCTTCTCGTGTGCAGACTTGAGGTCAGTGTAGAGCATACGGCAGAAGACGAAC-5′</td></tr>
<tr><td>Right2</td><td>5′-P-CACATCGATAGATCGGAAGAGCACACGTCTGAACTCCAGTCACATCTCGTATGCCGTCTTCTGCTTG-3′<br>3′-TCGAGTGTAGCTATCTAGCCTTCTCGTGTGCAGACTTGAGGTCAGTGTAGAGCATACGGCAGAAGACGAAC-5′</td></tr>
<tr><td>Right3</td><td>5′-P-CTACAAGATCAGATCGGAAGAGCACACGTCTGAACTCCAGTCACATCTCGTATGCCGTCTTCTGCTTG-3′<br>3′-TCGAGATGTTCTAGTCTAGCCTTCTCGTGTGCAGACTTGAGGTCAGTGTAGAGCATACGGCAGAAGACGAAC-5′</td></tr>
<tr><td>Right4</td><td>5′-P-CATTGGCATCGAGATCGGAAGAGCACACGTCTGAACTCCAGTCACATCTCGTATGCCGTCTTCTGCTTG-3′<br>3′-TCGAGTAACCGTAGCTCTAGCCTTCTCGTGTGCAGACTTGAGGTCAGTGTAGAGCATACGGCAGAAGACGAAC-5′</td></tr>
<tr><td>Right5</td><td>5′-P-CCACTGTATCGAAGATCGGAAGAGCACACGTCTGAACTCCAGTCACATCTCGTATGCCGTCTTCTGCTTG-3′<br>3′-TCGAGGTGACATAGCTTCTAGCCTTCTCGTGTGCAGACTTGAGGTCAGTGTAGAGCATACGGCAGAAGACGAAC-5′</td></tr>
<tr><td>Right6</td><td>5′-P-CTCAAGTATCGATAGATCGGAAGAGCACACGTCTGAACTCCAGTCACATCTCGTATGCCGTCTTCTGCTTG-3′<br>3′-TCGAGAGTTCATAGCTATCTAGCCTTCTCGTGTGCAGACTTGAGGTCAGTGTAGAGCATACGGCAGAAGACGAAC-5′</td></tr>
</table>

**Supplementary Tab. 4 | Internal-standard RBSs used in this study.** RBS sequences controlling translation of Bxb1-sfGFP are shown from transcriptional start base (+1) to the *bxb1* start codon with the variable 17-bp region highlighted in bold. Internal-standard RBSs were introduced into pASPIre3 and the resulting derivatives (pASPIre3_R1-R31) were used for the recording of calibration curves.

| No. | RBS sequence (5' to 3') |
|---|---|
| R1 | AAUGAAGAAUGAAAUAAAUAUUUAAUUUAU**AUUGUACGAUAGGUUUA**AUG |
| R2 | AAUGAAGAAUGAAAUAAAUAUUUAAUUUAU**AUCAUCGCAAGCUGCGU**AUG |
| R3 | AAUGAAGAAUGAAAUAAAUAUUUAAUUUAU**GUUAGCGCCACUGAAAG**AUG |
| R4 | AAUGAAGAAUGAAAUAAAUAUUUAAUUUAU**ACGUGUGUGUCGGAAAG**AUG |
| R5 | AAUGAAGAAUGAAAUAAAUAUUUAAUUUAU**CGGACAACGGAUACAUC**AUG |
| R6 | AAUGAAGAAUGAAAUAAAUAUUUAAUUUAU**UUACUAUCAGUAGUUAU**AUG |
| R7 | AAUGAAGAAUGAAAUAAAUAUUUAAUUUAU**GACAUGACGAGUAUGCC**AUG |
| R8 | AAUGAAGAAUGAAAUAAAUAUUUAAUUUAU**CCUUAUUGGGAAGCUGA**AUG |
| R9 | AAUGAAGAAUGAAAUAAAUAUUUAAUUUAU**ACUCUGGAUGUAAUGUG**AUG |
| R10 | AAUGAAGAAUGAAAUAAAUAUUUAAUUUAU**AACACCCGAGGGUUUAG**AUG |
| R11 | AAUGAAGAAUGAAAUAAAUAUUUAAUUUAU**CUCAGAAAGGCUAAGAC**AUG |
| R12 | AAUGAAGAAUGAAAUAAAUAUUUAAUUUAU**UGAAAGCGGGGUUCCUA**AUG |
| R13 | AAUGAAGAAUGAAAUAAAUAUUUAAUUUAU**CAAAUCAGGAAGAGACG**AUG |
| R14 | AAUGAAGAAUGAAAUAAAUAUUUAAUUUAU**GUAUGAAGAUACCCUAC**AUG |
| R15 | AAUGAAGAAUGAAAUAAAUAUUUAAUUUAU**UAAGGAUACUUACGCAC**AUG |
| R16 | AAUGAAGAAUGAAAUAAAUAUUUAAUUUAU**GGGCAGAACCUUAGGAG**AUG |
| R17 | AAUGAAGAAUGAAAUAAAUAUUUAAUUUAU**CACUAAGAGAGUAGAUU**AUG |
| R18 | AAUGAAGAAUGAAAUAAAUAUUUAAUUUAU**AAUGUGUGGGGAGAAUU**AUG |
| R19 | AAUGAAGAAUGAAAUAAAUAUUUAAUUUAU**UUUAGAGGAACGAACAU**AUG |
| R20 | AAUGAAGAAUGAAAUAAAUAUUUAAUUUAU**GAGAGGAGUAAGUGAUG**AUG |
| R21 | AAUGAAGAAUGAAAUAAAUAUUUAAUUUAU**CGAGGAGGUGCUAUUAU**AUG |
| R22 | AAUGAAGAAUGAAAUAAAUAUUUAAUUUAU**GGAGGAGAUUUAUAAUG**AUG |
| R23 | AAUGAAGAAUGAAAUAAAUAUUUAAUUUAU**GCUUUCUCGUUAUAUUA**AUG |
| R24 | AAUGAAGAAUGAAAUAAAUAUUUAAUUUAU**GCCUGAUCGGCAACGUA**AUG |
| R25 | AAUGAAGAAUGAAAUAAAUAUUUAAUUUAU**UAUACAGCUAUCCGGGA**AUG |
| R26 | AAUGAAGAAUGAAAUAAAUAUUUAAUUUAU**GUAAGGAGGCAAUUACC**AUG |
| R27 | AAUGAAGAAUGAAAUAAAUAUUUAAUUUAU**GACAGGAGGAACUAGAU**AUG |
| R28 | AAUGAAGAAUGAAAUAAAUAUUUAAUUUAU**ACGGGGGGAUUGUGACA**AUG |
| R29 | AAUGAAGAAUGAAAUAAAUAUUUAAUUUAU**GCGUGAGGGCGAAUAUU**AUG |
| R30 | AAUGAAGAAUGAAAUAAAUAUUUAAUUUAU**AUUAAGGAGGUUUUUUU**AUG |
| R31 | AAUGAAGAAUGAAAUAAAUAUUUAAUUUAU**CACUAAGGAGGUUUUUU**AUG |

### Plasmid Sequences

Please note that all plasmids are based on pSEVA291 and the sequence displayed as plain text below corresponds only to the part inserted into the original multiple cloning site as flanked by restriction sites for *Spe*I and *Not*I/*Pac*I.

### pASPIre1

**ACTAGT**AAGCTTAATTAATCTTTCTGCGAATTGAGATGACGCCACTGGCTGGGCGTCATCCCGGTTTCCCGGGTAAACACCACCGAAAAATAGT
TACTATCTTCAAAGCCACATTCGGTCGAAATATCACTGATTAACAGGCGGCTATGCTGGAGAAGATATTGCGCATGACACACTCTGACCTGTCG
CAGATATTGATTGATGGTCATTCCAGTCTGCTGGCGAAATTGCTGACGCAAAACGCGCTCACTGCACGATGCCTCATCACAAAATTTATCCAGC
GCAAAGGGACTTTTCAGGCTAGCCGCCAGCCGGGTAATCAGCTTATCCAGCAACGTTTCGCTGGATGTTGGCGGCAACGAATCACTGGTGTAAC
GATGGCGATTCAGCAACATCACCAACTGCCCGAACAGCAACTCAGCCATTTCGTTAGCAAACGGCACATGCTGACTACTTTCATGCTCAAGCTG
ACCGATAACCTGCCGCGCCTGCGCCATCCCCATGCTACCTAAGCGCCAGTGTGGTTGCCCTGCGCTGGCGTTAAATCCCGGAATCGCCCCCTGC
CAGTCAAGATTCAGCTTCAGACGCTCCGGGCAATAAATAATATTCTGCAAAACCAGATCGTTAACGGAAGCGTAGGAGTGTTTATCGTCAGCAT
GAATGTAAAAGAGATCGCCACGGGTAATGCGATAAGGGCGATCGTTGAGTACATGCAGGCCATTACCGCGCCAGACAATCACCAGCTCACAAAA
ATCATGTGTATGTTCAGCAAAGACATCTTGCGGATAACGGTCAGCCACAGCGACTGCCTGCTGGTCGCTGGCAAAAAAATCATCTTTGAGAAGT
TTTAACTGATGCGCCACCGTGGCTACCTCGGCCAGAGAACGAAGTTGATTATTCGCAATATGGCGTACAAATACGTTGAGAAGATTCGCGTTAT
TGCAGAAAGCCATCCCGTCCCTGGCGAATATCACGCGGTGACCAGTTAAACTCTCGGCGAAAAAGCGTCGAAAAGTGGTTACTGTCGCTGAATC
CACAGCGATAGGCGATGTCAGTAACGCTGGCCTCGCTGTGGCGTAGCAGATGTCGGGCTTTCATCAGTCGCAGGCGGTTCAGGTATCGCTGAGG
CGTCAGTCCCGTTTGCTGCTTAAGCTGCCGATGTAGCGTACGCAGTGAAAGAGAAAATTGATCCGCCACGGCATCCCAATTCACCTCATCGGCA
AAATGGTCCTCCAGCCAGGCCAGAAGCAAGTTGAGACGTGATGCGCTGTTTTCCAGGTTCTCCTGCAAACTGCTTTTACGCAGCAAGAGCAGTA
ATTGCATAAACAAGATCTCGCGACTGGCGGTCGAGGGTAAATCATTTTCCCCTTCCTGCTGTTCCATCTGTGCAACCAGCTGTCGCACCTGCTG
CAATACGCTGTGGTTAACGCGCCAGTGAGACGGATACTGCCCATCCAGCTCTTGTGGCAGCAACTGATTCAGCCCGGCGAGAAACTGAAATCGA
TCCGGCGAGCGATACAGCACATTGGTCAGACACAGATTATCGGTATGTTCATACAGATGCCGATCATGATCGCGTACGAAACAGACCGTGCCAC
CGGTGATGGTATAGGGCTGCCCATTAAACACATGAATACCCGTGCCATGTTCGACAATCACAATTTCATGAAAATCATGATGATGTTCAGGAAA
ATCCGCCTGCGGGAGCCGGGGTTCTATCGCCACGGACGCGTTACCAGACGGAAAAAAATCCACACTATGTAATACGGTCATACTGGCCTCCTGA
TGTCGTCAACACGGCGAAATAGTAATCACGAGGTCAGGTTCTTACCTTAAATTTTCGACGGAAAACCACGTAAAAAACGTCGATTTTTCAAGAT
ACAGCGTGAATTTTCAGGAAATGCGGTGAGCATCACATCTGCAGTAATCGGCCGGCTTGTCGACGACGGCGGTCTCCGTCGTCAGGATCATCCG
GGCCCATGGGATATCTATAATTTTGTTTAACTTTAAGAAGGAGATATACGCATGCATAGCAAGGGCGAAGAAGATAACATGGCCATCATCAAGG
AGTTCATGCGCTTCAAGGTGCACATGGAGGGCTCCGTGAACGGCCACGAGTTCGAGATCGAGGGCGAGGGCGAGGGCCGCCCCTACGAGGGCAC
CCAGACCGCCAAGCTGAAGGTGACCAAGGGTGGCCCCCTGCCCTTCGCCTGGGACATCCTGTCCCCTCAGTTCATGTACGGCTCCAAGGCCTAC
GTGAAGCACCCCGCCGACATCCCCGACTACTTGAAGCTGTCCTTCCCCGAGGGCTTCAAGTGGGAGCGCGTGATGAACTTCGAGGACGGCGGCG
TGGTGACCGTGACCCAGGACTCCTCCTTGCAGGACGGCGAGTTCATCTACAAGGTGAAGCTGCGCGGCACCAACTTCCCCTCCGACGGCCCCGT
AATGCAGAAGAAGACAATGGGCTGGGAGGCCTCCTCCGAGCGGATGTACCCCGAGGACGGCGCCCTGAAGGGCGAGATCAAGCAGAGGCTGAAG
CTGAAGGACGGCGGCCACTACGACGCTGAGGTCAAGACCACCTACAAGGCCAAGAAGCCCGTGCAGCTGCCCGGCGCCTACAACGTCAACATCA
AGTTGGACATCACCTCCCACAACGAGGACTACACCATCGTGGAACAGTACGAACGCGCCGAGGGCCGCCACTCCACCGGCGGCATGGACGAGCT
GTACAAGAGGCCTTAAGCAAAAAACCCCGCTTCGGCGGGGTTTTTTCGCTGCGATATCTAATAAGGGTTTGTACCGTACACCACTGAGACCGCG
GTGGTTGACCAGACAAACCACGAAGGTTCTGTTAAGTAACTGAACCCAATGTCGTTAGGCTAGCACAATACCTAGGACTGAGCTAGCCGTCAAT
GACGCTTACCTCTTAAGAGGTCACTGACCTAACAGGATCCCACCACAATTCAGCAAATTGTGAACATCATCACGTTCATCTTTCCCTGGTTGCC
AATGGCCCATTTTCCTGTCAGTAACGAGAAGGTCGCGAATTCAGGCGCTTTTTAGACTGGTCGTAATGAAGGGTACCATAAATGCTTCAATAAT
ATTGAAAAGGATTCTAGATGCGTGCACTGGTTGTTATTCGTCTGAGCCGTGTTACCGATGCAACCACCAGTCCGGAACGTCAGCTGGAAAGCTG
TCAGCAGCTGTGTGCACAGCGTGGTTGGGATGTTGTTGGTGTTGCAGAGGATCTGGATGTTAGCGGTGCAGTTGATCCGTTTGATCGTAAACGT
CGTCCGAATCTGGCACGTTGGCTGGCATTTGAAGAACAGCCGTTTGATGTTATTGTTGCCTATCGTGTTGATCGTCTGACCCGTAGCATTCGTC
ATCTGCAACAGCTGGTTCATTGGGCAGAAGATCATAAAAAACTGGTTGTGAGCGCAACCGAAGCACATTTTGATACCACCACCCCGTTTGCAGC
AGTTGTTATTGCACTGATGGGCACCGTTGCACAGATGGAACTGGAAGCAATTAAAGAACGTAATCGTAGCGCAGCCCATTTTAACATTCGTGCA
GGTAAATATCGTGGTAGCCTGCCTCCGTGGGGTTATCTGCCGACGCGTGTAGATGGTGAATGGCGTCTGGTTCCTGACCCGGTTCAGCGTGAAC
GTATTCTGGAAGTATATCATCGTGTGGTGGATAATCATGAACCGCTGCATCTGGTTGCACATGATCTGAATCGTCGTGGTGTTCTGAGTCCCAA
AGATTATTTTGCTCAGCTGCAAGGTCGTGAACCGCAGGGTCGTGAATGGTCTGCAACCGCACTGAAACGTAGCATGATTAGCGAAGCAATGCTG
GGTTATGCAACCCTGAATGGTAAAACCGTTCGTGATGATGATGGTGCACCGCTGGTTCGTGCAGAACCGATTCTGACACGTGAACAGCTGGAAG
CACTGCGTGCCGAACTGGTTAAAACCAGCCGTGCAAAACCGGCAGTTAGCACCCCGAGCCTGCTGCTGCGTGTTCTGTTTTGTGCAGTTTGTGG
TGAACCGGCATACAAATTTGCCGGTGGTGGTCGTAAACATCCGCGTTATCGTTGTCGTAGCATGGGTTTTCCGAAACATTGTGGTAATGGTACA
GTTGCAATGGCAGAATGGGATGCATTTTGCGAAGAACAGGTTCTGGATCTGCTGGGTGATGCCGAACGTCTGGAAAAAGTTTGGGTTGCAGGTA
GCGATAGCGCAGTTGAACTGGCCGAAGTTAATGCGGAACTGGTCGATCTCACCAGTCTGATTGGAAGTCCCGCATATCGTGCGGGTAGTCCTCA
GCGTGAAGCACTGGATGCACGTATTGCAGCACTGGCAGCACGTCAAGAAGAACTGGAAGGTCTGGAAGCACGTCCGAGCGGTTGGGAATGGCGT
GAAACAGGTCAGCGTTTTGGTGATTGGTGGCGTGAGCAGGATACCGCAGCAAAAAATACCTGGCTGCGTAGTATGAATGTTCGCCTGACCTTTG
ATGTTCGCGGTGGCCTGACCCGCACCATTGATTTTGGCGATCTGCAAGAATATGAACAGCATCTGCGTCTGGGTAGCGTTGTTGAACGTCTGCA
TACCGGCATGAGCACCGGCTGCTGTCCAGGTTGTTGCGCTGCTAACGACGAAAACTACGCTCTGGCTGCTTAAGAATTCGC**GCGGCCGC.**

**pASPIre2**

**ACTAGT**AAGCTTAATTAATCTTTCTGCGAATTGAGATGACGCCACTGGCTGGGCGTCATCCCGGTTTCCCGGGTAAACACCACCGAAAAATAGT
TACTATCTTCAAAGCCACATTCGGTCGAAATATCACTGATTAACAGGCGGCTATGCTGGAGAAGATATTGCGCATGACACACTCTGACCTGTCG
CAGATATTGATTGATGGTCATTCCAGTCTGCTGGCGAAATTGCTGACGCAAAACGCGCTCACTGCACGATGCCTCATCACAAAATTTATCCAGC
GCAAAGGGACTTTTCAGGCTAGCCGCCAGCCGGGTAATCAGCTTATCCAGCAACGTTTCGCTGGATGTTGGCGGCAACGAATCACTGGTGTAAC
GATGGCGATTCAGCAACATCACCAACTGCCCGAACAGCAACTCAGCCATTTCGTTAGCAAACGGCACATGCTGACTACTTTCATGCTCAAGCTG
ACCGATAACCTGCCGCGCCTGCGCCATCCCCATGCTACCTAAGCGCCAGTGTGGTTGCCCTGCGCTGGCGTTAAATCCCGGAATCGCCCCCTGC
CAGTCAAGATTCAGCTTCAGACGCTCCGGGCAATAAATAATATTCTGCAAAACCAGATCGTTAACGGAAGCGTAGGAGTGTTTATCGTCAGCAT
GAATGTAAAAGAGATCGCCACGGGTAATGCGATAAGGGCGATCGTTGAGTACATGCAGGCCATTACCGCGCCAGACAATCACCAGCTCACAAAA
ATCATGTGTATGTTCAGCAAAGACATCTTGCGGATAACGGTCAGCCACAGCGACTGCCTGCTGGTCGCTGGCAAAAAAATCATCTTTGAGAAGT
TTTAACTGATGCGCCACCGTGGCTACCTCGGCCAGAGAACGAAGTTGATTATTCGCAATATGGCGTACAAATACGTTGAGAAGATTCGCGTTAT
TGCAGAAAGCCATCCCGTCCCTGGCGAATATCACGCGGTGACCAGTTAAACTCTCGGCGAAAAAGCGTCGAAAAGTGGTTACTGTCGCTGAATC
CACAGCGATAGGCGATGTCAGTAACGCTGGCCTCGCTGTGGCGTAGCAGATGTCGGGCTTTCATCAGTCGCAGGCGGTTCAGGTATCGCTGAGG
CGTCAGTCCCGTTTGCTGCTTAAGCTGCCGATGTAGCGTACGCAGTGAAAGAGAAAATTGATCCGCCACGGCATCCCAATTCACCTCATCGGCA
AAATGGTCCTCCAGCCAGGCCAGAAGCAAGTTGAGACGTGATGCGCTGTTTTCCAGGTTCTCCTGCAAACTGCTTTTACGCAGCAAGAGCAGTA
ATTGCATAAACAAGATCTCGCGACTGGCGGTCGAGGGTAAATCATTTTCCCCTTCCTGCTGTTCCATCTGTGCAACCAGCTGTCGCACCTGCTG
CAATACGCTGTGGTTAACGCGCCAGTGAGACGGATACTGCCCATCCAGCTCTTGTGGCAGCAACTGATTCAGCCCGGCGAGAAACTGAAATCGA
TCCGGCGAGCGATACAGCACATTGGTCAGACACAGATTATCGGTATGTTCATACAGATGCCGATCATGATCGCGTACGAAACAGACCGTGCCAC
CGGTGATGGTATAGGGCTGCCCATTAAACACATGAATACCCGTGCCATGTTCGACAATCACAATTTCATGAAAATCATGATGATGTTCAGGAAA
ATCCGCCTGCGGGAGCCGGGGTTCTATCGCCACGGACGCGTTACCAGACGGAAAAAAATCCACACTATGTAATACGGTCATACTGGCCTCCTGA
TGTCGTCAACACGGCGAAATAGTAATCACGAGGTCAGGTTCTTACCTTAAATTTTCGACGGAAAACCACGTAAAAAACGTCGATTTTTCAAGAT
ACAGCGTGAATTTTCAGGAAATGCGGTGAGCATCACATCTGCAGTAATCGGCCGGCTTGTCGACGACGGCGGTCTCCGTCGTCAGGATCATCCG
GGCCCATGGGATATCTATAATTTTGTTTAACTTTAAGAAGGAGATATACGCATGCATAGCAAGGGCGAAGAAGATAACATGGCCATCATCAAGG
AGTTCATGCGCTTCAAGGTGCACATGGAGGGCTCCGTGAACGGCCACGAGTTCGAGATCGAGGGCGAGGGCGAGGGCCGCCCCTACGAGGGCAC
CCAGACCGCCAAGCTGAAGGTGACCAAGGGTGGCCCCCTGCCCTTCGCCTGGGACATCCTGTCCCCTCAGTTCATGTACGGCTCCAAGGCCTAC
GTGAAGCACCCCGCCGACATCCCCGACTACTTGAAGCTGTCCTTCCCCGAGGGCTTCAAGTGGGAGCGCGTGATGAACTTCGAGGACGGCGGCG
TGGTGACCGTGACCCAGGACTCCTCCTTGCAGGACGGCGAGTTCATCTACAAGGTGAAGCTGCGCGGCACCAACTTCCCCTCCGACGGCCCCGT
AATGCAGAAGAAGACAATGGGCTGGGAGGCCTCCTCCGAGCGGATGTACCCCGAGGACGGCGCCCTGAAGGGCGAGATCAAGCAGAGGCTGAAG
CTGAAGGACGGCGGCCACTACGACGCTGAGGTCAAGACCACCTACAAGGCCAAGAAGCCCGTGCAGCTGCCCGGCGCCTACAACGTCAACATCA
AGTTGGACATCACCTCCCACAACGAGGACTACACCATCGTGGAACAGTACGAACGCGCCGAGGGCCGCCACTCCACCGGCGGCATGGACGAGCT
GTACAAGAGGCCTTAAGCAAAAAACCCCGCTTCGGCGGGGTTTTTTCGCTGCGATATCTAATAAGGGTTTGTACCGTACACCACTGAGACCGCG
GTGGTTGACCAGACAAACCACGAAGGTTCTGTTAAGTAACTGAACCCAATGTCGTTAGGCTAGCACAATACCTAGGACTGAGCTAGCCGTCAAT
GACGCTTACCTCTTAAGAGGTCACTGACCTAACAGGATCCCACCACAATTCAGCAAATTGTGAACATCATCACGTTCATCTTTCCCTGGTTGCC
AATGGCCCATTTTCCTGTCAGTAACGAGAAGGTCGCGAATTCAGGCGCTTTTTAGACTGGTCGTAATGAAGGGTACCATAAATGCTTCAATAAT
ATTGAAAAGGATTCTAGATGCGAGCTCTGGTTGTTATTCGTCTGAGCCGTGTTACCGATGCAACCACCAGTCCGGAACGTCAGCTGGAAAGCTG
TCAGCAGCTGTGTGCACAGCGTGGTTGGGATGTTGTTGGTGTTGCAGAGGATCTGGATGTTAGCGGTGCAGTTGATCCGTTTGATCGTAAACGT
CGTCCGAATCTGGCACGTTGGCTGGCATTTGAAGAACAGCCGTTTGATGTTATTGTTGCCTATCGTGTTGATCGTCTGACCCGTAGCATTCGTC
ATCTGCAACAGCTGGTTCATTGGGCAGAAGATCATAAAAAACTGGTTGTGAGCGCAACCGAAGCACATTTTGATACCACCACCCCGTTTGCAGC
AGTTGTTATTGCACTGATGGGCACCGTTGCACAGATGGAACTGGAAGCAATTAAAGAACGTAATCGTAGCGCAGCCCATTTTAACATTCGTGCA
GGTAAATATCGTGGTAGCCTGCCTCCGTGGGGTTATCTGCCGACGCGTGTAGATGGTGAATGGCGTCTGGTTCCTGACCCGGTTCAGCGTGAAC
GTATTCTGGAAGTATATCATCGTGTGGTGGATAATCATGAACCGCTGCATCTGGTTGCACATGATCTGAATCGTCGTGGTGTTCTGAGTCCCAA
AGATTATTTTGCTCAGCTGCAAGGTCGTGAACCGCAGGGTCGTGAATGGTCTGCAACCGCACTGAAACGTAGCATGATTAGCGAAGCAATGCTG
GGTTATGCAACCCTGAATGGTAAAACCGTTCGTGATGATGATGGTGCACCGCTGGTTCGTGCAGAACCGATTCTGACACGTGAACAGCTGGAAG
CACTGCGTGCCGAACTGGTTAAAACCAGCCGTGCAAAACCGGCAGTTAGCACCCCGAGCCTGCTGCTGCGTGTTCTGTTTTGTGCAGTTTGTGG
TGAACCGGCATACAAATTTGCCGGTGGTGGTCGTAAACATCCGCGTTATCGTTGTCGTAGCATGGGTTTTCCGAAACATTGTGGTAATGGTACA
GTTGCAATGGCAGAATGGGATGCATTTTGCGAAGAACAGGTTCTGGATCTGCTGGGTGATGCCGAACGTCTGGAAAAAGTTTGGGTTGCAGGTA
GCGATAGCGCAGTTGAACTGGCCGAAGTTAATGCGGAACTGGTCGATCTCACCAGTCTGATTGGAAGTCCCGCATATCGTGCGGGTAGTCCTCA
GCGTGAAGCACTGGATGCACGTATTGCAGCACTGGCAGCACGTCAAGAAGAACTGGAAGGTCTGGAAGCACGTCCGAGCGGTTGGGAATGGCGT
GAAACAGGTCAGCGTTTTGGTGATTGGTGGCGTGAGCAGGATACCGCAGCAAAAAATACCTGGCTGCGTAGTATGAATGTTCGCCTGACCTTTG
ATGTTCGCGGTGGCCTGACCCGCACCATTGATTTTGGCGATCTGCAAGAATATGAACAGCATCTGCGTCTGGGTAGCGTTGTTGAACGTCTGCA
TACCGGCATGAGCACCGGCGGTGGCAGCGGCGGTTCTGGTGGCTCTAGCAAAGGAGAAGAACTTTTCACTGGAGTTGTCCCAATTCTTGTTGAA
TTAGATGGTGATGTTAATGGGCACAAATTTTCTGTCCGTGGAGAGGGTGAAGGTGATGCTACAAACGGAAAACTCACCCTTAAATTTATTTGCA
CTACTGGAAAACTACCTGTTCCGTGGCCAACACTTGTCACTACTCTGACCTATGGTGTTCAATGCTTTTCCCGTTATCCGGATCACATGAAACG
GCATGACTTTTTCAAGAGTGCCATGCCCGAAGGTTATGTACAGGAACGCACTATATCTTTCAAAGATGACGGGACCTACAAGACGCGTGCTGAA
GTCAAGTTTGAAGGTGATACCCTTGTTAATCGTATCGAGTTAAAGGGTATTGATTTTAAAGAAGATGGAAACATTCTTGGACACAAACTCGAGT
ACAACTTTAACTCACACAATGTATACATCACGGCAGACAAACAAAAGAATGGAATCAAAGCTAACTTCAAAATTCGCCACAACGTTGAAGATGG
TTCCGTTCAACTAGCAGACCATTATCAACAAAATACTCCAATTGGCGATGGCCCTGTCCTTTTACCAGACAACCATTACCTGTCGACACAATCT
GTCCTTTCGAAAGATCCCAACGAAAAGCGTGACCACATGGTCCTTCTTGAGTTTGTAACTGCTGCTGGGATTACACATGGCATGGATGAGCTCT
ACAAAAGGCCTGCTGCTAACGACGAAAACTACGCTCTGGCTGCTTAATGA**GCGGCCGC**.

**pASPIre3**

**ACTAGT**AAGCTTCACATCTGCAGTAATCGGCCGGCTTGTCGACGACGGCGGTCTCCGTCGTCAGGATCATCCGGGCCCATGGATCGCTTAGTCA
CCTTTGGGCCACGGTCCGCTACCTTACAGGAATAGTACTCGTCCTTTAATTTCCCTTGCATATATGTTGCGTTTCTTCGACCTAGTACTCGCTC
CCTTAGGAGAAAGACAGATAGCTTCTTACCCGCCATGGGGGTTTGTACCGTACACCACTGAGACCGCGGTGGTTGACCAGACAAACCACGAAGG
TTCTGTTAAGTAACTGAACCCAATGTCGTTAGTGACGCTTACCTCTTAAGAGGTCACTGACCTAACAGGATCCCACCACAATTCAGCAAATTGT
GAACATCATCACGTTCATCTTTCCCTGGTTGCCAATGGCCCATTTTCCTGTCAGTAACGAGAAGGTCGCGAATTCAGGCGCTTTTTAGACTGGT
CGTAATGAAGGGTACCATAAATGCTTCAATAATATTGAAAAGGATTCTAGATGCGAGCTCTGGTTGTTATTCGTCTGAGCCGTGTTACCGATGC
AACCACCAGTCCGGAACGTCAGCTGGAAAGCTGTCAGCAGCTGTGTGCACAGCGTGGTTGGGATGTTGTTGGTGTTGCAGAGGATCTGGATGTT
AGCGGTGCAGTTGATCCGTTTGATCGTAAACGTCGTCCGAATCTGGCACGTTGGCTGGCATTTGAAGAACAGCCGTTTGATGTTATTGTTGCCT
ATCGTGTTGATCGTCTGACCCGTAGCATTCGTCATCTGCAACAGCTGGTTCATTGGGCAGAAGATCATAAAAAACTGGTTGTGAGCGCAACCGA
AGCACATTTTGATACCACCACCCCGTTTGCAGCAGTTGTTATTGCACTGATGGGCACCGTTGCACAGATGGAACTGGAAGCAATTAAAGAACGT
AATCGTAGCGCAGCCCATTTTAACATTCGTGCAGGTAAATATCGTGGTAGCCTGCCTCCGTGGGGTTATCTGCCGACGCGTGTAGATGGTGAAT
GGCGTCTGGTTCCTGACCCGGTTCAGCGTGAACGTATTCTGGAAGTATATCATCGTGTGGTGGATAATCATGAACCGCTGCATCTGGTTGCACA
TGATCTGAATCGTCGTGGTGTTCTGAGTCCCAAAGATTATTTTGCTCAGCTGCAAGGTCGTGAACCGCAGGGTCGTGAATGGTCTGCAACCGCA
CTGAAACGTAGCATGATTAGCGAAGCAATGCTGGGTTATGCAACCCTGAATGGTAAAACCGTTCGTGATGATGATGGTGCACCGCTGGTTCGTG
CAGAACCGATTCTGACACGTGAACAGCTGGAAGCACTGCGTGCCGAACTGGTTAAAACCAGCCGTGCAAAACCGGCAGTTAGCACCCCGAGCCT
GCTGCTGCGTGTTCTGTTTTGTGCAGTTTGTGGTGAACCGGCATACAAATTTGCCGGTGGTGGTCGTAAACATCCGCGTTATCGTTGTCGTAGC
ATGGGTTTTCCGAAACATTGTGGTAATGGTACAGTTGCAATGGCAGAATGGGATGCATTTTGCGAAGAACAGGTTCTGGATCTGCTGGGTGATG
CCGAACGTCTGGAAAAAGTTTGGGTTGCAGGTAGCGATAGCGCAGTTGAACTGGCCGAAGTTAATGCGGAACTGGTCGATCTCACCAGTCTGAT
TGGAAGTCCCGCATATCGTGCGGGTAGTCCTCAGCGTGAAGCACTGGATGCACGTATTGCAGCACTGGCAGCACGTCAAGAAGAACTGGAAGGT
CTGGAAGCACGTCCGAGCGGTTGGGAATGGCGTGAAACAGGTCAGCGTTTTGGTGATTGGTGGCGTGAGCAGGATACCGCAGCAAAAAATACCT
GGCTGCGTAGTATGAATGTTCGCCTGACCTTTGATGTTCGCGGTGGCCTGACCCGCACCATTGATTTTGGCGATCTGCAAGAATATGAACAGCA
TCTGCGTCTGGGTAGCGTTGTTGAACGTCTGCATACCGGCATGAGCACCGGCGGTGGCAGCGGCGGTTCTGGTGGCTCTAGCAAAGGAGAAGAA
CTTTTCACTGGAGTTGTCCCAATTCTTGTTGAATTAGATGGTGATGTTAATGGGCACAAATTTTCTGTCCGTGGAGAGGGTGAAGGTGATGCTA
CAAACGGAAAACTCACCCTTAAATTTATTTGCACTACTGGAAAACTACCTGTTCCGTGGCCAACACTTGTCACTACTCTGACCTATGGTGTTCA
ATGCTTTTCCCGTTATCCGGATCACATGAAACGGCATGACTTTTTCAAGAGTGCCATGCCCGAAGGTTATGTACAGGAACGCACTATATCTTTC
AAAGATGACGGGACCTACAAGACGCGTGCTGAAGTCAAGTTTGAAGGTGATACCCTTGTTAATCGTATCGAGTTAAAGGGTATTGATTTTAAAG
AAGATGGAAACATTCTTGGACACAAACTCGAGTACAACTTTAACTCACACAATGTATACATCACGGCAGACAAACAAAAGAATGGAATCAAAGC
TAACTTCAAAATTCGCCACAACGTTGAAGATGGTTCCGTTCAACTAGCAGACCATTATCAACAAAATACTCCAATTGGCGATGGCCCTGTCCTT
TTACCAGACAACCATTACCTGTCGACACAATCTGTCCTTTCGAAAGATCCCAACGAAAAGCGTGACCACATGGTCCTTCTTGAGTTTGTAACTG
CTGCTGGGATTACACATGGCATGGATGAACTCTACAAAAGGCCTGCTGCTAACGACGAAAACTACGCTCTGGCTGCTTAATAA**GCGGCCGC.**

### pASPIre6

**ACTAGT**CGCCAGGGTTTTCCCAGTCACGACGCGGCCGCAAGCTTCCATGGGGGTTTGTACCGTACACCACTGAGACCGCGGTGGTTGACCAGAC
AAACCACGAAGGTTCTGTTAAGTAACTGAACCCAATGTCGTTAGTGACGCTTACCTCTTAAGAGGTCACTGACCTAACAGGATCCCACCACAAT
TCAGCAAATTGTGAACATCATCACGTTCATCTTTCCCTGGTTGCCAATGGCCCATTTTCCTGTCAGTAACGAGAAGGTCGCGAATTCAGGCGCT
TTTTAGACTGGTCGTAATGAAGGGTACCATAAATGCTTCAATAATATTGAAAAGGATTCTAGATGCGAGCTCACAGCCCCTCTGTCGTCGCCGA
CGTCTGTAATGTAGCCTCATTGTGATTCCCATGGGCCCGGATGATCCTGACGACGGAGACCGCGGTGGTTGACCAGACAAACCACGAAGGTTCT
GTTAAGTAACTGAACCCAATGTCGTTAGTGACGCTTACCTCTTAAGAGGTCACTGACCTAACAGGATCCCACCACAATTCAGCAAATTGTGAAC
ATCATCACGTTCATCTTTCCCTGGTTGCCAATGGCCCATTTTCCTGTCAGTAACGAGAAGGTCGCGAATTCAGGCGCTTTTTAGACTGGTCGTA
ATGAAGGGTACCATAAATGCTTCAATAATATTGAAAAGGATTCTAGATGCGAGCTC**TTAATTAA**.

### References (Supplementary Information)


1. Jeschek, M., Gerngross, D. & Panke, S. Rationally reduced libraries for combinatorial pathway optimization minimizing experimental effort. *Nat. Commun.* **7**, 11163 (2016).
2. Bonnet, J., Subsoontorn, P. & Endy, D. Rewritable digital data storage in live cells via engineered control of recombination directionality. *Proc. Natl. Acad. Sci. U.S.A.* **109**, 8884-8889 (2012).
3. Martinez-Garcia, E., Aparicio, T., Goni-Moreno, A., Fraile, S. & de Lorenzo, V. SEVA 2.0: an update of the Standard European Vector Architecture for de-/re-construction of bacterial functionalities. *Nucleic Acids Res.* **43**, D1183-D1189 (2015).
4. Datsenko, K. A. & Wanner, B. L. One-step inactivation of chromosomal genes in Escherichia coli K-12 using PCR products. *Proc. Natl. Acad. Sci. U.S.A.* **97**, 6640-6645 (2000).
