## Supplementary material for "Large-scale DNA-based phenotypic recording and deep learning enable highly accurate sequence-function mapping": Machine Learning (ML) Annex

### Machine Learning Annex to:

† Equal contribution

### A. Details of the deep learning model

#### 1. Notation

In the following paragraphs, lower-case italic letters such as $z$ describe scalar values, lower-case bold italic letters $\boldsymbol{z}$ represent vectors (1D array), upper-case bold italic letters $\boldsymbol{Z}$ symbolize matrices (2D array) and upper-case bold letters $\mathbf{Z}$ stand for tensors (3D array). Sizes of arrays are noted as follows: $l$ for a vector which has $l$ elements along its first dimension, $l \times c$ for a 2D array which has $l$ rows and $c$ columns; and $l \times c \times p$ for a 3D array that contains $p$ $l \times c$ 2D arrays.

Other useful notations are shown below:

| Notation | Meaning |
|---|---|
| $z$ | scalar |
| $\boldsymbol{z}$ | vector (1D array) |
| $(z_1, \dots, z_l)$ | specification of the vector components for a vector of size $l$ |
| $z_i$ | scalar element of the vector $\boldsymbol{z}$ |
| $\boldsymbol{Z}$ | matrix (2D array) |
| $\boldsymbol{Z}_{i,.}$ | $i$'th row of matrix $\boldsymbol{Z}$ |
| $\boldsymbol{Z}_{.,i}$ | $i$'th column of the matrix $\boldsymbol{Z}$ |
| $Z_{i,j}$ | scalar element of the matrix $\boldsymbol{Z}$ |
| $\mathbf{Z}$ | tensor (3D array) |
| $\mathbf{Z}_{i,:}$ | 2D array slice of the tensor $\mathbf{Z}$ for which the first dimension is fixed |
| $\mathbf{Z}_{.,i,.}$ | 2D array slice of the tensor $\mathbf{Z}$ for which the second dimension is fixed |
| $\mathbf{Z}_{:,i}$ | 2D array slice of the tensor $\mathbf{Z}$ for which the third dimension is fixed |
| $\mathbf{Z}_{i,j,.}$ | vector slice of the tensor $\mathbf{Z}$ for which the first and second dimensions are fixed |
| $\mathbf{Z}_{i,.,j}$ | vector slice of the tensor $\mathbf{Z}$ for which the first and third dimensions are fixed |
| $\mathbf{Z}_{.,i,j}$ | vector slice of the tensor $\mathbf{Z}$ for which the second and third dimensions are fixed |
| $\mathrm{Z}_{i,j,k}$ | scalar element of the tensor $\mathbf{Z}$ |
| $(\boldsymbol{X}, y)$ | a datapoint or sample-target pair, with sample $\boldsymbol{X}$ and its ground truth target value $y$ |
| $(\mathbf{X}, \boldsymbol{y})$ or $D$ | the entire dataset |
| $\mathbf{W}$ or $\mathbf{W}_\mathrm{m}$ | parameters of the model |

| | |
|---|---|
| $p_{\mathbf{W}}(y\|\boldsymbol{X})$ | the likelihood of the target $y$ given sample $\boldsymbol{X}$ and the weights $\mathbf{W}$ |
| $q(y,\boldsymbol{X})$ | the (unknown) ground truth distribution of the datapoints $(\boldsymbol{X},y) \in (\mathbf{X},\boldsymbol{y})$ |
| $q(y\|\boldsymbol{X})$ | the (unknown) ground truth conditional distribution of the target $y$ given sample $\boldsymbol{X}$ |
| $r(\mathbf{W}\|D)$ | posterior probability of the parameters $\mathbf{W}$ given the dataset $D$ |
| $\mu \longmapsto \delta(\mu-\theta)$ | Dirac delta function centered in $\theta$ |

#### 2. Glossary and main machine learning concepts

##### *a. Main machine learning concepts*

A **parameter** or **weight** of a model is a variable that is optimized by fitting the model to the data, often by minimizing, with a gradient-based method (see below), a function of the difference between the model output and some ground-truth values.

The action of **learning** in machine learning corresponds to fixing the weights of a model by fitting the model to the data. In machine learning, we can alternatively talk about **training** a model.

**Supervised learning** is a task in machine learning that consists of learning a function that maps an input to an output based on available input-output pairs. Both input and output can take several data types, such as unstructured data types (i.e. scalars, vectors, arrays) or structured data types (i.e. graphs, permutations).

A **target** or **output target** is the scalar (or other more complex data type) variable that characterizes an input sample $\boldsymbol{X}$. In general, the aim of machine learning models is to predict the target given the corresponding input sample, which is why it is also called the output target. For each input sample, the **ground truth value** of the target can either be available, for example measured experimentally, or not available.

A **gradient-based method** is an iterative algorithm to solve optimization problems, i.e. minimizing or maximizing a quantity with respect to some parameters. Its particularity is that the search directions are given by the gradient of the quantity to optimize with respect to the parameters.

A **cost function** is, in one of its most common forms, composed of two terms, a) an **empirical risk** or **expected loss**, which is a quantity that is correlated to the difference between the predicted output targets and their ground truth values for several input samples and b) a **regularization term**, which increases with the absolute values of the parameters of the model. The cost function is a **training**

**criterion** that is minimized, often with a gradient-based method, to find the best fit (i.e. best parameters) of the model to the training data, model that should at the same time be able to generalize to unseen data thanks to the regularization term.

A **hyperparameter** is a parameter or a function whose value or type is set before learning the weights of the model. Hyperparameters are different from the weights of the model and in general regroup a) parameters that cannot be learned using a gradient-based method or b) parameters that can lead the model optimization to a bad minimum if it were considered as a common parameter, for example by making the model perfectly fit to the training data but not able to generalize in the test data or c) functions that are part of the model design and that cannot be learned with a gradient-based method. The hyperparameters control aspects of the model behavior such as: time, memory cost, representational capacity of the model, degree of regularization of the model, and, more generally, ability of the model to infer correct results for seen and unseen inputs.

In machine learning, a **predictive model** or **predictor** is a model that uses observations about samples and their target values to be able to guess the target value of any unseen input sample.

The ability of the model to predict correctly the targets of new examples is known as **generalization**.

##### *b. An overview of machine learning*

A key step in the development of a machine learning model is to separate the available data into non-overlapping **training**, **validation** and **test datasets**, prior to model fitting. In supervised learning, as it is the case in this manuscript, each dataset consists of a collection of tuples of samples and their respective ground-truth targets. The three datasets have distinct roles, which can be roughly described as follows.

The **training set** is utilized to fit the model, that is, it is the dataset on which the parameters of the model are optimized. Often, including in this manuscript, this is done by minimizing a **cost function**, which typically combines an aggregated measure of the discrepancy between ground-truth targets and model predictions in the training set (**empirical risk**) with a penalty for excessive model complexity (**regularization**). Many alternative ways to design such a cost function have been explored in the literature, and each choice will generally lead to a different fitted model.

The main purpose of the **validation set** is to provide a way to evaluate the performance of fitted models in an unbiased manner, thereby allowing to choose between multiple candidate fitted models. Since the different candidate models might differ in their complexity, for example, by having a different number of parameters or by having been trained by minimizing cost functions that give different importance to the model complexity penalty, their predictive performance can only be meaningfully compared using sample-target pairs other than those used for model fitting. That is, model selection cannot be made on the basis of predictive performance on the training set, as this criterion would be irremediably biased in favor of the most complex models. In this manuscript, different candidate models will be obtained by choosing different **hyperparameters**. Roughly, these correspond to variables that control high-level aspects of the model such as how many parameters it has, what type of computations it performs or which cost function is used to fit its parameters. In this context, we will henceforth refer to the process of selecting between different models as **hyperparameter search**. In summary, hyperparameter search is performed by a) fitting a different model to the training set for each candidate hyperparameter combination; b) evaluating an aggregated measure of the discrepancy between ground-truth targets and model predictions in the validation set, typically tailored to the specific application at hand and c) selecting the model (and its corresponding hyperparameter setting) that achieves the lowest error, according to the metric of choice, in the validation set.

Finally, the **test set** is used to evaluate the predictive performance of the trained model selected according to the procedure described above. It is crucial to ensure that the test set has no overlap not only with the training set, but also with the validation set. Otherwise, contributions to the overall error due to imperfect model selection would not be properly accounted for.

In situations where data is scarce, random variability arising due to the allocation of different sample-target pairs to either the training, validation or test sets can become a concern. In those cases, however, the small sample size might imply it is computationally feasible to repeat the process for multiple random splits to obtain estimates of the expected predictive performance and of its standard deviation due to different splits.

##### *c. Definitions of common array operations in machine learning and deep learning*

An **affine transformation** can be seen as the extension of a linear function (in $\mathbb{R}$) to multiple dimensions space $\mathbb{R}^n$. It is a composition of a translation (i.e. transformation that moves every point in the same direction without deformation) and a linear map. For example, in $\mathbb{R}$, an affine transformation is a linear function which can be written as $f: \mathbb{R} \longrightarrow \mathbb{R}$, $f(x) = ax + b$, where $x$ is the input scalar, the translation is represented by the addition of the intercept $b$ and the linear map by the multiplication of the slope parameter $a$ with $x$. In the two-dimensional case, i.e. in $\mathbb{R}^2$, an affine transformation $g: \mathbb{R}^2 \longrightarrow \mathbb{R}^2$ can be written as $g(x) = Ax + b$. In this equation, $x$ is a datapoint in $\mathbb{R}^2$, $g(x)$ the output of the affine transformation, the translation is represented by the addition of a vector $b$ and the linear map by a matrix multiplication with $A$. Following the same principles, an affine transformation generalizes to multiple dimensions and consists informally of an array multiplication of the input followed by an addition.

A **normalization operation** is an operation that changes the scale of several variables to often bring them to a common scale. For example, the normalized variables can be centered on zero and have a standard deviation equal to one. In the context of deep learning models, the normalization most often happens across samples, either to individual array elements or across elements as well.

An **elementwise operation** is an operation that occurs to each scalar value (i.e. element) of an array.

A **kernel transformation** is a function that maps a space to another one. Typically, this function is applied to the input data and maps the input data to a different space in order to represent the input data differently. For example, it can transform arrays in $\mathbb{R}^2$ into arrays in $\mathbb{R}^n$. We call **representation** the output of the mapping function. Often, this transformation is implicit in machine learning models, i.e. we do not need to calculate the representation to build a predictive model as the latter can depend on the inner product between representations instead of depending on the representations directly.

##### *d. Other keywords*

In this appendix, we refer to (**sequence) motif** to describe a nucleotide pattern.

A **mixture distribution** of a random variable is a weighted average of probability distributions of other random variables. The probability density of a mixture is called **mixture density**.

#### 3. Deep learning model composition

When applied to genomic sequences and for many subfields of other applications (computer vision, natural language processing), a deep learning model is usually composed of a **sequential stack of modules**, which are functions that gradually transform an input array into an output array. These modules are called **layers**. A layer can be generally thought of as being itself composed of a series of operations which are sequentially applied to its input. In this work, we consider a layer to be composed of up to three stages, a) the first stage is an **affine transformation applied** to the input array of the layer, followed by b) an optional second stage which consists of a **normalization operation**, and c) a third stage which consists of **elementwise operations**, most often non-linear, which are applied to every element of the output of the second stage and are called **activation functions**. The composition of a layer is depicted in Figure 1. A layer is therefore the composition of an affine transformation, followed by optional normalization operations and (non-linear) elementwise operations. The operation that is computed in the first stage defines the type of layer. The inputs as well as the outputs of a layer are 1D, 2D or 3D arrays. The dimensionality of an output may differ from that of an input.

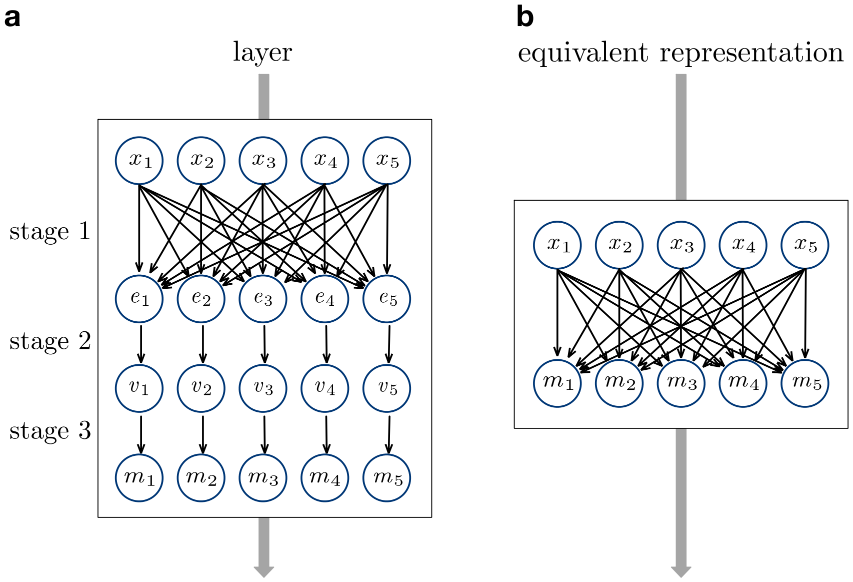

***Figure 1: Layer Representation. a**, Representation of the three stages that compose a layer. The output of the layer* $\boldsymbol{m}$ *is a function of the input* $\boldsymbol{x}$*. The representations of stages 1 and 2 can differ depending on the type of affine transformation and of normalization operation. Dependence between different inputs* $\boldsymbol{x}$ *can be introduced in stage 2 (not represented here).* ***b**, Often, only the first stage affine operation of a layer is represented. The dependence between different inputs is rarely represented.*

For example,

Figure **2** illustrates a model with two layers. In this model, we assume that the layers do not include a normalization stage. The model takes a vector $\boldsymbol{x}$ as input and transforms it into an output scalar $y$. The first function, $f_1$, is the first layer of the model that transforms the input vector $\boldsymbol{x}$ into the vector $\boldsymbol{m}$, such that $\boldsymbol{m} = f_1(\boldsymbol{x})$. $f_1$ can be written as $f_1 = h \circ g$, where $g$ is an affine transformation and $h$ an elementwise function. In a standard representation of deep learning architecture, it is usual to represent $f_1$ as one function and not separate it into $h$ and $g$, as shown in

Figure **3**. The second function $f_2$ corresponds to the second layer of the model and also applies an affine transformation followed by an elementwise operation on its input $\boldsymbol{m}$. The function $f_2$ transforms $\boldsymbol{m}$ into an output scalar $y$, such that $y = f_2(\boldsymbol{m})$. The model can be summarized as the composition of the two functions, $f_1$ and $f_2$, such that $y = f_2(\boldsymbol{m}) = f_2 \circ f_1(\boldsymbol{x})$. While the figure shows a network that takes a vector as input and generates a scalar, deep learning models do not have theoretical constraints in the dimensionality of their input and output.

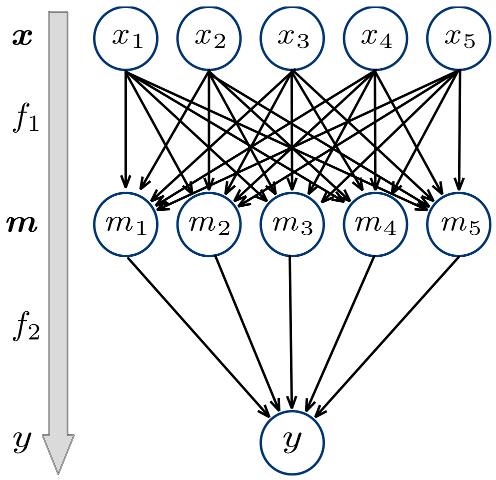

***Figure 2: Example of a Two-Layer Fully-Connected Neural Network.*** *In this example,* $\boldsymbol{x}$ *is a vector composed of 5 scalars, as well as* $\boldsymbol{m}$*. The input* $\boldsymbol{x}$ *is first transformed to the array* $\boldsymbol{m}$ *through a first function* $f_1$ *and then into* $y$ *through* $f_2$*. Each scalar in* $\boldsymbol{m}$ *is a function of scalars of the input* $\boldsymbol{x}$*. The output scalar* $y$ *is a function of scalars of* $\boldsymbol{m}$*. Fully-connected layers will be explained in Section 6.c.*

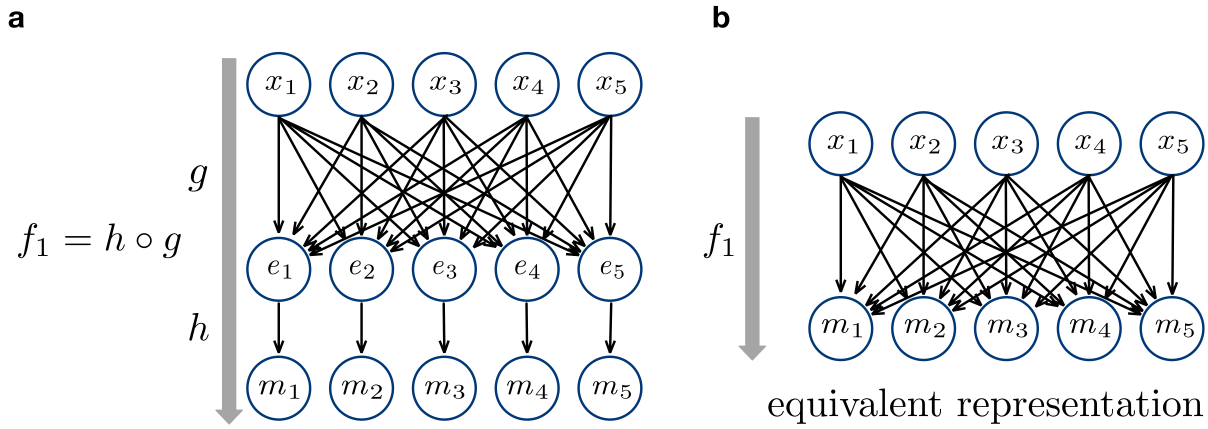

***Figure 3: Layer Representation for two Stages. a****, Illustration of the composition* $f_1 = g \circ h$*, with the affine operation* $g$ *and the elementwise operation* $h$*.* ***b****, Often, and in this manuscript, only the operations of the first stage of the layer are represented.*

The **depth** of the model is given by the number of layers that are stacked between the input and the output. The parameters of the layer functions are called **weights**; most often they are randomly initialized and are learned during training. The goal of the learning process is to determine the weights that collectively minimize a function of the difference between the model outputs, which are dependent on the sequence inputs, and their respective ground truth values, measured experimentally or otherwise known independently of the model (see Section A.9).

**Implementation:** Our core ResNet model is composed of typical layers such as **convolutional** (see Section A.6.b) and **fully-connected** (see Section A.6.c) ones, that include common activation functions such as variants of the **ReLU function** (see Section A.6.f) and **normalization operations** (see Section

A.6.g). Additional simpler components of the model architecture introduced in this manuscript, such as **skip-connections** (see Section A.6.d) and **flattened operations** (see Section A.6.e) are also described.

#### 4. Workflow

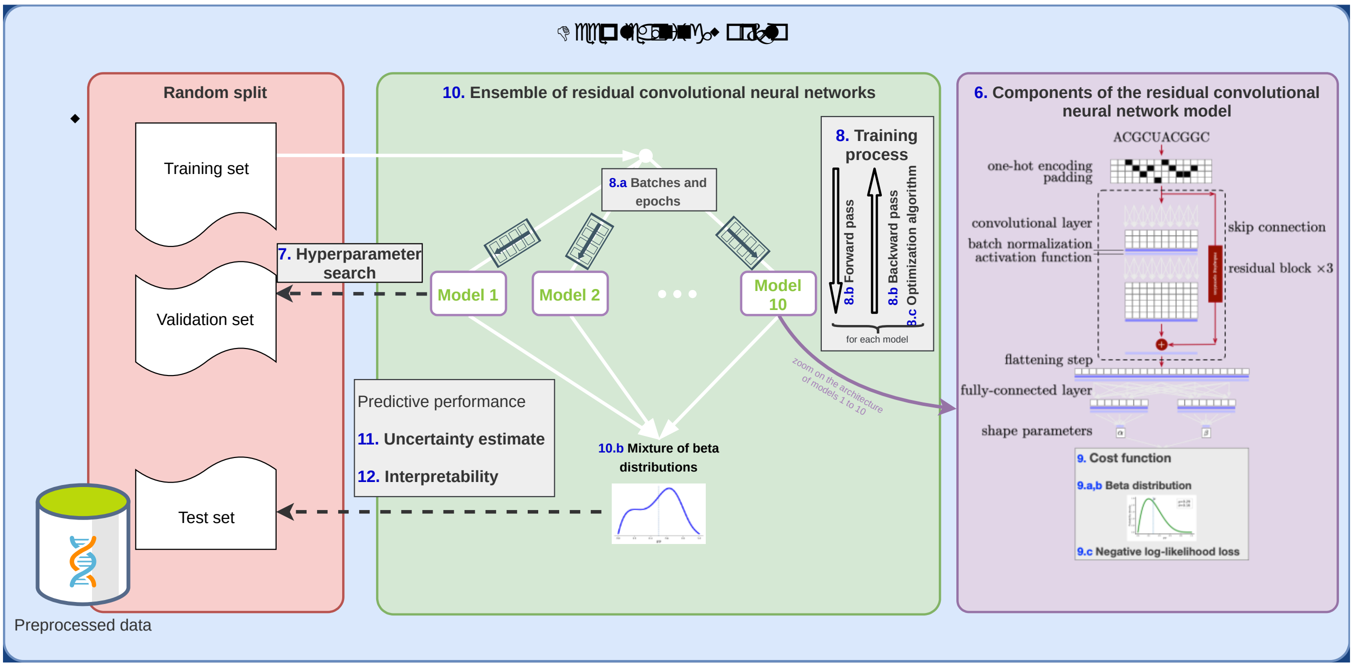

***Figure 4: SAPIENs Workflow.*** *The different components of the model SAPIENs (Sequence-Activity Prediction In Ensemble of Networks), presented in the main manuscript, are referred to according to their respective title numbering in this appendix.*

#### 5. Dataset and main objective of the machine learning model

The dataset is composed of hundreds of thousands of RNA sequences whose activity has been quantified. More specifically, the RNA sequences are ribosome binding sites (RBS) in *E. Coli* (Salis, 2011) and the term activity refers to their ability to control the translation of a protein from the mRNA transcript. In our biological experiments, the activity of the ribosome binding site is estimated by the normalized integral of the flipping profiles ($IFP_{0\text{-}480min}$), profiles that are experimentally measured.

We build a machine learning model, termed SAPIENs (Sequence-Activity Prediction In Ensemble of Networks), in order to predict the activity (as estimated experimentally) of any RBS sequence. To do so, we use an ensemble (see Section A.10) of ten **residual convolutional neural network** (ResNet) (He, 2016; Xie, 2017). We train the model on a pool of sequences – activity pairs, with the objective to predict the RBS activity, approximated by the $IFP_{0\text{-}480min}$ values. The output target is therefore the variable that corresponds to the $IFP_{0\text{-}480min}$ values, we know its ground truth values for the RBS sequences that are present in the dataset, as measured by the $IFP_{0\text{-}480min}$ values, but not for the other RBS sequences. In order to obtain a good predictive performance, SAPIENs learns to automatically detect sequence motifs, and rules to combine them. At the end, the performance of the model is evaluated by comparing its output to the ground truth values of the targets.

Sections A.6 to A.9 describe the fundamental constituents of a single ResNet model and of its training process. Section A.10 introduces SAPIENs as an ensemble of ten ResNet models. Section A.11 presents how well-calibrated uncertainty estimates are obtained. Section A.12 gives an introduction to an integrated gradient attribution method, used in deep learning to attribute the predictions to specific elements of the input.

#### 6. Components of the residual convolutional neural network model

The ResNet model that we present in the main manuscript comprises several components identified in the rightmost box of the workflow in

Figure **4** and described below in this section.

##### *a. One-hot encoded input*

Notation:

In this section, for 2D and 3D input or output arrays, we refer to:

- the first dimension as the **spatial dimension**, the number of rows as the number of **positions**, corresponding to the position of an RNA base along the RNA sequence.
- the second dimension as the **channel dimension,** each column as a **channel**, corresponding to the nature of an RNA base (A/C/G/U).
- the third dimension as the **sample dimension**, corresponding to different RNA sequences.

The **input** of the model is an RNA sequence $\boldsymbol{s} = (s_1, \dots, s_l)$, of length $l$, where each element $s_i$, $i \in \{1, \dots, l\}$, represents a base $\in \{A, C, G, U\}$ (Alipanahi, 2015). However, the neural network requires a numerical input, therefore the input sequences must be transformed into numerical arrays.

**Implementation:** As done in most genomic applications of deep learning, the sequence of bases $\boldsymbol{s}$ is **one-hot encoded** into an array of $c = 4$ binary **channels** of length $l$, one for each base: the A-channel, the C-channel, the G-channel and the U-channel. As illustrated in Figure 5, **A**s are represented by a one in the first column and zeros in the other columns, **C**s are represented by a one in the second column and a zero in the other three, **G**s are represented by a one in the third column and a zero elsewhere and **U**s are represented by a one in the fourth column and a zero in the other columns. The size of the one-hot encoded array is therefore $l \times c = l \times 4$. Figure 5 shows how the sequence AUCGGCU is one-hot encoded.

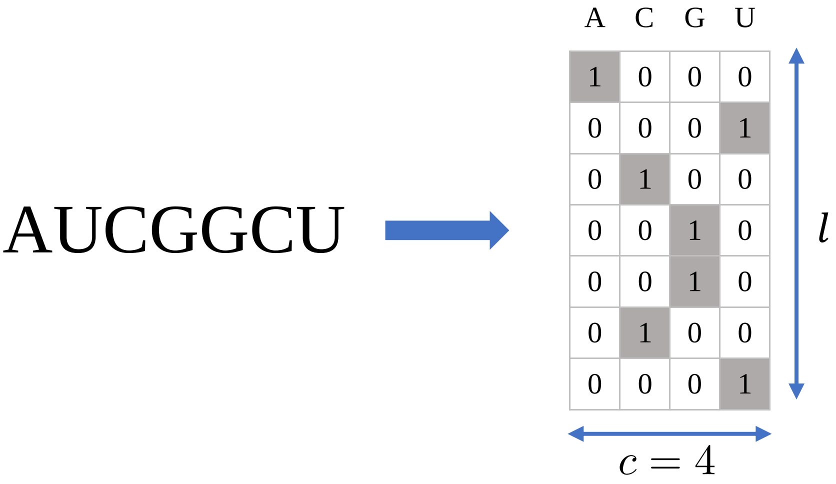

***Figure 5: One-Hot Encoding***. *The genomic sequence AUCGGCU is one-hot encoded. The resulting array is a 2D matrix that has as many rows as the genomic sequence is long (l) and that has 4 channels (c).* ***A****s are represented by a one in the first column and zeros in the other columns,* ***C****s are represented by a one in the second column and a zero in the other three,* ***G****s are represented by a one in the third column and a zero elsewhere and* ***U****s are represented by a one in the fourth column and a zero in the other columns.*

output of the same length $l$ along the spatial dimension (see Section A.6.b). As we will see, the number of all-zeros rows that are added depends on hyperparameters of the convolutional layer that is applied to the input.

The padded one-hot encoded array serves as **input** to the first layer of the neural network.

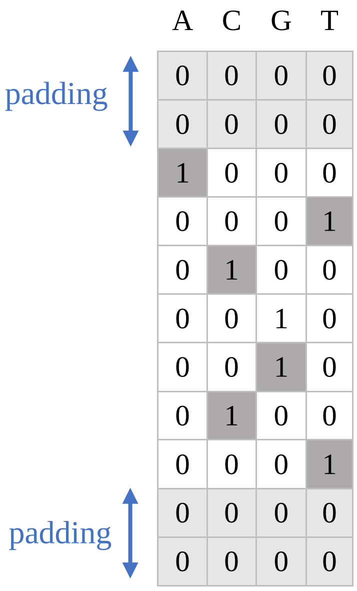

***Figure 6: Padding.*** *The one-hot encoded array is extended on the spatial dimension by two rows on both sides. The extension is filled with zeros.*

##### *b. Convolutional layer*

Notation:

In this section, for 2D and 3D input or output arrays, we refer to:

- the first dimension as the **spatial dimension**, the number of rows as the number of **positions**. For the input array, the first dimension corresponds to the position of an RNA base along the RNA sequence and this relation can be extended to the next convolutional layers outputs.
- the second dimension as the **channel dimension,** each column as a **channel**. For the input array, the second dimension corresponds to the nature of an RNA base (A/C/G/U).
- the third dimension as the **sample dimension**, corresponding to different samples, here RNA sequences.

*First stage of a convolutional layer*

The **convolution** (Goodfellow, 2016; Alipanahi, 2015) is a vector-to-vector operation that applies the same function $f$ to sliding windows covering subvectors of the input and generates a vector output. By extension, convolutions can also be applied to arrays, in which case the function $f$ is applied to subarrays. The function $f$ is a Frobenius inner product, whose variable is a subarray and whose parameters are defined in an object called the **convolution matrix**. Given an input subarray $\boldsymbol{X}$ of size $l' \times c$ and a convolution matrix $\boldsymbol{W}$ of the same size as $\boldsymbol{X}$, the resulting scalar $u$ of the Frobenius inner product can be written as:

$$u = \sum_{i=1}^{l'} \sum_{j=1}^{c} W_{i,j} X_{i,j} \quad (1)$$

In practice, the Frobenius inner product is slightly modified and a **bias term** $b$ is added to its scalar output.

$$u = \sum_{i=1}^{l'} \sum_{j=1}^{c} W_{i,j} X_{i,j} + b \quad (2)$$

The convolution[1] operation consists of applying this Frobenius inner product to sliding windows covering subarrays of the input that have the same dimensions as $\boldsymbol{W}$ (the sliding windows are shifted by a predefined number of elements at a time in one or several dimensions). Each convolution operation generates an output array. For example, let $\boldsymbol{X}$ be an input matrix, of size $l \times c$, $\boldsymbol{W}$ a convolution matrix

[1] The term that describes more accurately the type of operation that is used in convolutional layers (excluding the bias term) is a cross-correlation. However, we follow the conventions in the deep learning literature and use the term convolution instead of cross-correlation.

of size $l' \times c'$ (with $l' \leq l$ and $c' \leq c$) and $b$ a scalar term. Let's assume that the sliding windows are shifted by only one element at a time (i.e. **stride** of 1). The 2D output array $\boldsymbol{U}$ will then be of size $(l - l' + 1) \times (c - c' + 1)$ and each element $U_{k,m}$ can be written as:

$$U_{k,m} = \sum_{i=1}^{l'} \sum_{j=1}^{c'} W_{i,j} X_{i+k,j+m} + b \quad (3)$$

We can notice that $\boldsymbol{W}$ and $b$ are independent of the coordinates of the input array the inner products are applied to.

Convolutional layers are typically composed of several of these convolution operations applied to the same input array and computed in parallel. The output arrays of each convolution are stacked together into a larger array that is the output of the convolutional layer. Additionally, in most cases, convolutional layers are either 1-dimensional (1d), 2-dimensional (2d) or 3-dimensional (3d) depending on the type of convolution operations that is used (1d, 2d or 3d). Informally, a 1-dimensional convolution slides along one axis only and generates a vector output, a 2-dimensional convolution slides along two axes and generates a 2D array output (as presented Equation (3)) and a 3-dimensional convolution slides along three axes and generates a 3D array output.

The parameters of the convolution matrices as well as the bias terms are known as **weights**. The convolution matrices are alternatively termed as **filters** or **detectors**. The weights of the filters and the bias terms are free parameters learned during model training, in order to get the output of the model close to some ground truth values, obtained independently of the model (see Section A.9).

In the context of convolutional neural networks applied to genomic sequences, it is common to use **1-dimensional (1d) convolutions** over the input channels. Therefore, we will focus on this type of convolutional layer in this section. Let's consider a given input array $\boldsymbol{X}$ of size $l \times c$. We pass it through the first stage of a convolutional layer with *one* filter of size $r \times c$, where $r$ is the length of each filter along the spatial dimension. For example, in case the input of the convolutional layer is the one-hot encoded RNA sequence, $r$ corresponds to the length of subsequences composed of adjacent RNA bases. To characterize the convolutional layer, we define the filter array $\boldsymbol{W}$ of size $r \times c$ and a bias scalar $b$. Let $\boldsymbol{u}$ be the output of the first stage of the convolutional layer and $i \in \{1, \dots, l - r\}$ be a position along the spatial dimension of the output $\boldsymbol{u}$. Each output element can be written as:

$$u_i = \sum_{p=1}^{r} \sum_{v=1}^{c} W_{p,v} X_{p+i,v} + b \quad (4)$$

As a consequence, the 1d-convolution consists of scanning, by the detector, the input array along its spatial dimension, which results in an output array with as many elements as subarrays that have been scanned. The mechanism of a 1d convolutional layer with one filter is represented in

Figure 7.

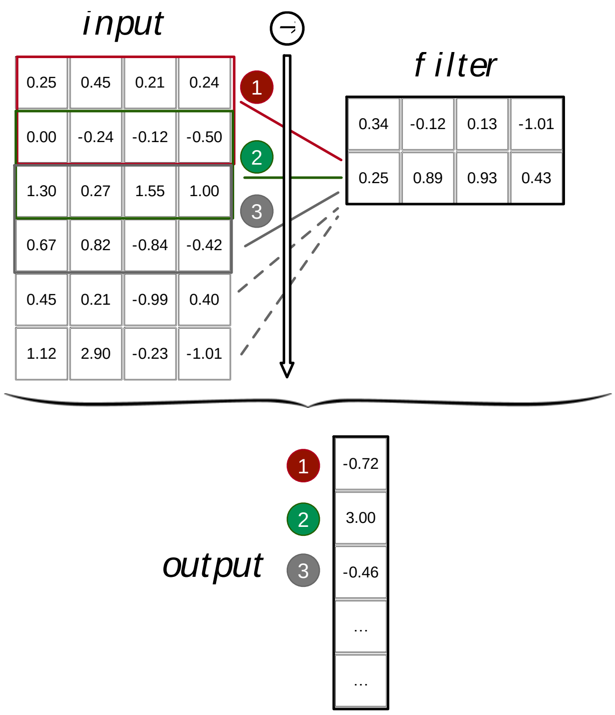

***Figure 7: 1-Dimensional Convolutional Layer****. The Frobenius inner product is calculated between the filter of size* $2 \times 4$ *(top right) and each subarray of size* $2 \times 4$ *of the input (top left). The input is scanned following the ordering illustrated by the numbers from 1 to 3. The resulting output is shown at the bottom. For simplicity, we account for a filter without bias weight. We can observe that the number of scalar elements of the output is smaller than the number of rows of the input. The input has six rows and the filter has two rows, therefore the input contains only five subarrays of the size of the filter.*

Let's assume that the convolutional layer has only one filter. Let's have $l$ the length along the spatial dimension of the (non-padded) input array, $c$ the number of channels of the input array and $r$ the length of the filter. The size of the filter is therefore $r \times c$ and the output of the convolutional stage will be of size $(l - r + 1) \times 1$, by sliding the detector from the beginning to the end of the input array along its spatial dimension. With this mechanism, after each convolutional layer, the spatial dimension of the layers' output is shorter by $r - 1$ elements than the one of its input, as we can see in

Figure 7. After a finite number of convolutional layers, the size of the output could be reduced to $1 \times 1$. In order to avoid this phenomenon and to gain flexibility in the number of layers and sizes of filters of the model, it is possible to use a **padding mechanism** called *"***same***"*. The latter consists of extending the input of each convolutional layer along the spatial dimension, with $r - 1$ channels of zeros, in a symmetric fashion. As a consequence, the modified input is of size $(l + r - 1) \times c$ and the convolutional layer's output is of size $l \times 1$, keeping the spatial dimension's length unchanged. The effect of padding the input of a 1d convolutional layer is illustrated in

Figure **8**.

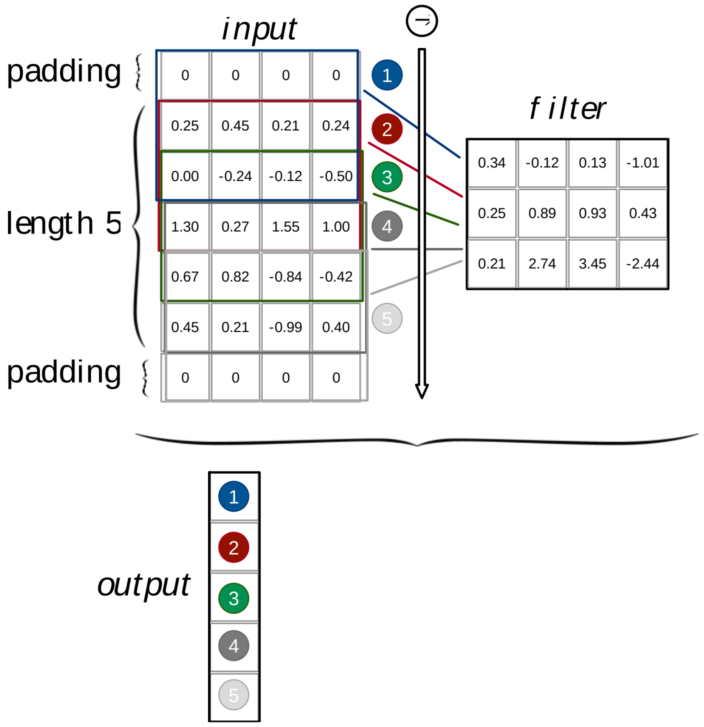

***Figure 8: Padding "same" of the Input of a 1d-Convolutional Layer.*** *The length of the spatial dimension is represented by the number of rows of each array. The input has originally 5 rows and the filter 3 rows. If the input were not padded, the output's spatial dimension would be of length 5 – 3 + 1 = 3. As we wish to keep the same number of positions along the spatial dimension in the input as in the output, we pad the input with two rows of zeros, one on both extremities of the spatial dimension. As a consequence, after the 1d convolution, the output has 5 rows, instead of 3. For simplicity, we account for a filter without bias weights.*

However, the number of filters can be arbitrarily large, as far as computational resources allow. Let $k$ be the number of filters for a given convolutional layer. The number of channels in the output array would then be $k$ and the layer output would be of size $l \times k$ . An illustration of a convolutional layer containing several filters is available in Figure 9.

Let's consider a given input array $\boldsymbol{X}$ of size $l \times c$. We pass it through a convolutional layer with $k$ filters of size $r \times c$, where $r$ is the length of the spatial dimension of each filter. To this end, we define $k$ filter arrays $\mathbf{W}_{:,j}$ of size $r \times c$ and bias scalars $b_j$, for $j \in \{1, \dots, k\}$. The input array $\boldsymbol{X}$ is padded with the "same" padding, therefore its size becomes $(l + r - 1) \times c$, we keep the notation $\boldsymbol{X}$ for simplification purposes. Let $\boldsymbol{U}$ be the output of the first stage of the convolutional layer, $i \in \{1, \dots, l\}$ the position along the spatial dimension and $j \in \{1, \dots, k\}$ the position along the second dimension. Each output scalar element can be written as:

$$U_{i,j} = \sum_{m=1}^{r}\sum_{v=1}^{c} X_{m+i,v} W_{m,v,j} + b_j \quad (5)$$

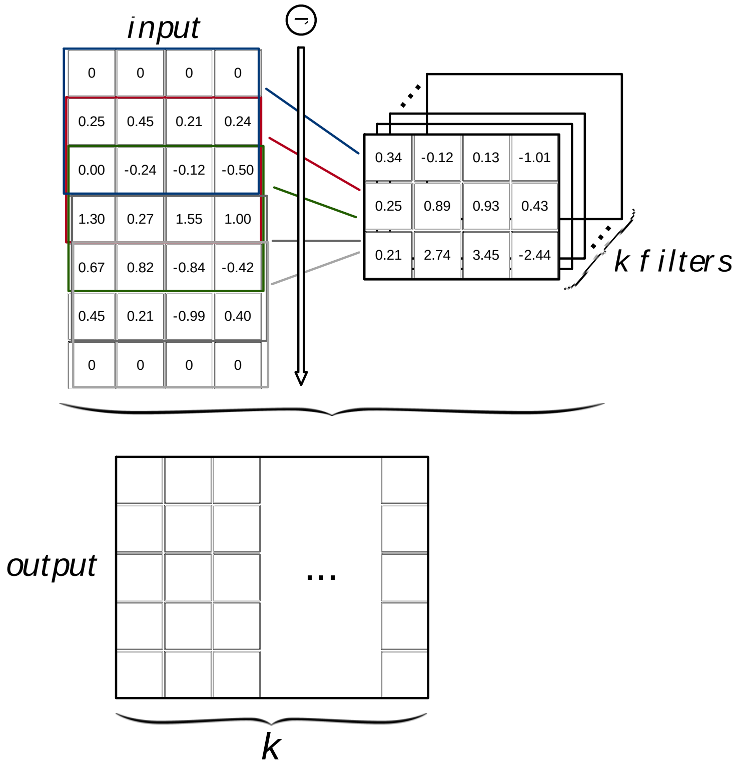

*Figure 9: **1d-Convolutional Layer with k Filters.** k filters scan the input from top to bottom along the spatial dimension. Each convolution between the input and a filter results in a column in the output array. Therefore, if there are k filters, the output has k columns. For simplicity, we account for filters without bias weights.*

**Implementation:** Our model contains several convolutional layers stacked one after the other, as schematically drawn in

Figure **10**. The number of layers was determined by evaluating the model's predictive performance on a held-out validation set, to select a model that generalizes outside of the training set (see Section A.7). All convolutional layers, including intermediate ones, are 1-dimensional. We used the padding mechanism "same" for the input of each convolutional layer.

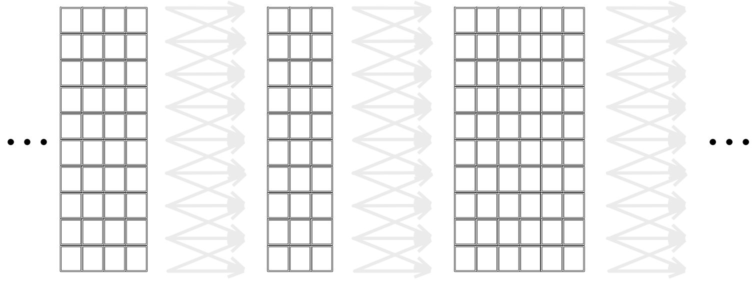

***Figure 10: Example of two Stacked Convolutional Layers.***

##### *Second stage of a convolutional layer*

The second stage of a convolutional layer is optional. It is a normalization operation that will be described in Section A.6.g. This second stage function does not change the size the convolution operations output.

##### *Third stage of a convolutional layer*

The third stage of a convolutional layer is a non-linear elementwise operation applied to the output of the convolution or normalization operations. This third stage function does not change the size of the array it is applied to. Most often, the same elementwise function is applied to each output scalar element, the function that is chosen is monotonously increasing and it is a non-linear function in order to introduce non-linearities in the model. The functions that are applied in the third stage of any layer are called **activation functions**. More details about activation functions will be given in Section A.6.f.

##### *Convolutional layer interpretation*

Informally, the role of a convolutional layer is to scan the subarrays of the previous layer output and detect contiguous groups of features, often referred to as **patterns**. The Frobenius inner product between the filter and a subarray that is being scanned is high if the subarray is similar to the filter and low if the subarray is very different from the filter, as it is explained in detail in the next paragraph and illustrated in Figure 11. Therefore, the action of each filter can be understood as looking for a specific pattern of interest, represented by the filter weights, along the spatial dimension of the input. Similarly, its output scalar elements can be seen as indicating the degree of agreement between the pattern represented by the filter and the subarray, as quantified by the Frobenius inner product. In summary, it is possible to view the convolution operation as a pattern scan. Since a convolutional layer typically contains multiple filters, several patterns can be searched in parallel such that each separate channel of the convolutional layer output corresponds to the result of scanning for each of the patterns represented by the layer filters. In summary, the channels of the output array indicate the similarity of the patterns represented by the filters at diverse positions of the input array. The output of the convolution operation is often subsequently normalized, e.g. by normalizing each channel separately to ensure all of them have comparable magnitude. Finally, an activation function, typically an elementwise nonlinear function, is applied to the (possibly normalized) output of the convolution. In particular, nonlinear activations allow stacked convolutional layers to behave nonlinearly, despite being built around an affine transformation, and thus allow deep convolutional networks to approximate complex nonlinear functions.

In case a convolutional layer is the first layer of the model, the subarrays that are scanned are those of the model input. In our model, the filters of the first convolutional layer scan subarrays of the one-hot encoded sequences. As these filters have four channels, one for each base, they can be interpreted as position weight matrices (PWMs). Several sequence motifs are searched in parallel and the agreement of each motif at different positions of the input sequence is represented by each channel of the convolutional layer output. While the same phenomenon is at work in the downstream convolutional layers, we can notice two major differences: a) the inputs do not have necessarily four channels, i.e. the number of channels will generally correspond to the number of filters in the previous layer and b) the channels do not represent the four bases, they represent agreement with motifs scanned by the previous layer. Thus, it is not possible anymore to interpret the filters of the convolutions as PWMs in such a straight-forward manner. Instead, the second convolutional layer can be understood roughly as looking for patterns of presence or absence of several RNA sequence motifs rather than patterns of single bases. With depth, these combinations become more complex, i.e. a larger number of motifs can be combined. Figure 11 illustrates how a stack of several convolutional layers can combine motifs together. In summary, it is possible to interpret a stack of convolutional layers as a way to create rules to combine sequence motifs such that the combinations would be predictive of the output target.

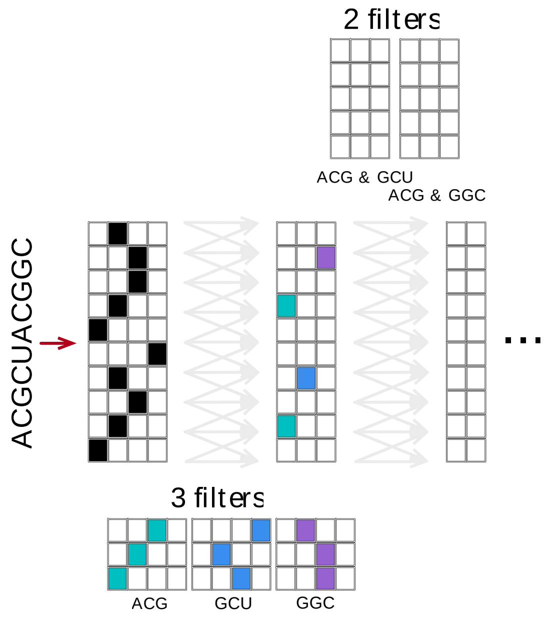

*Figure 11: **Example of a Series of two Convolutional Layers Applied to a One-Hot Encoded RNA Sequence.** The first layer has three filters, which respectively show a high inner product with the motifs ACG, GCU and GGC. The second layer looks for combinations of motifs that the first layer selected. The second layer has two filters, one that correlates with subarrays that contain the motifs ACG and GCU, and the second one that correlates with subarrays that contain the motifs ACG and GGC.*

**Implementation:** The first layers of our network are 1d convolutional layers. The total number of layers is a hyperparameter. Each convolutional layer has additionally two hyperparameters, the number of filters $k$ and the filter length $r$. The number of filters and the filter length can differ between convolutional layers. All the hyperparameters are optimized during a hyperparameter search phase, which is described in Section A.7. For each convolutional layer, the coefficients of the filters ($\mathbf{W}$ and $\boldsymbol{b}$) are optimized by gradient-based optimization during the training process (see Section A.8.c), to obtain output values that are close to some ground truth values, measured independently of the model.

##### c. *Fully-connected layer*

*First stage of a fully-connected layer*

The core operation of **fully-connected** layers (Goodfellow, 2016; Alipanahi, 2015) is a Frobenius inner product between a weight matrix and *all* input elements. Therefore, while each inner product only covers subarrays of the input in a typical convolutional layer, the "filter" of fully-connected layers is of the exact same size as the input: an inner product is computed between the whole input array $\boldsymbol{X}$, of size $l \times c$, and a weight matrix $\boldsymbol{W}$, of the same dimensionality. A bias scalar is then added to the inner product output. During the first stage of the fully-connected layer, the sequence of operation a) inner product and b) addition of a bias scalar is computed several times in parallel, for different weight matrices and different bias terms. The output of the first stage of the fully-connected layer is a vector, with as many elements as there are weight matrices.

Let $\boldsymbol{u}$ be the output of the first stage of the layer, $\boldsymbol{X}$ the input array of dimensions $l \times c$ and, $k$ the number of weight matrices $\mathbf{W}_{:,:,j}, j \in \{1, \dots, k\}$ and the number of bias vector elements $b_j$. $\boldsymbol{u}$ is therefore of size $k$ and is calculated as follows:

$$u_j = \sum_{m=1}^{l} \sum_{v=1}^{c} X_{m,v} \mathrm{W}_{m,v,j} + b_j \quad (6)$$

Figure **12** illustrates the mechanism of a fully-connected layer for an input $\boldsymbol{x}$, which we chose to be a vector to simplify the figure. The same inner product mechanism can be applied to input matrices or tensors: *all* input elements contribute to the output scalar elements.

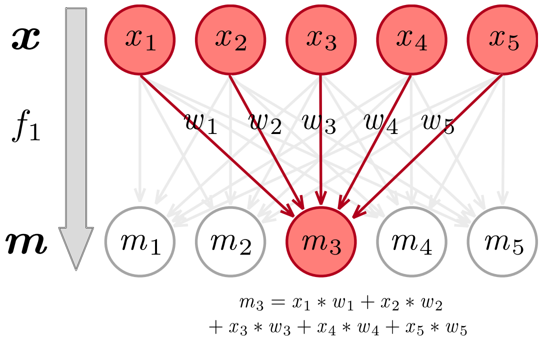

*Figure 12: Example of a Fully-Connected Layer. Each element of the output vector $\boldsymbol{m}$ is a function of the input scalars $x_i, i \in \{1, \dots, 5\}$. For example, the output scalar $m_3$ is a linear combination of the input scalars $x_i, i \in \{1, \dots, 5\}$, with the weights $\boldsymbol{w}$. Equivalently, $m_3$ is equal to the inner product between $\boldsymbol{x}$ and $\boldsymbol{w}$. For simplicity, we do not represent the bias weights.*

##### *Second and third stages of a fully-connected layer*

The second and third stages of a fully-connected layer are similar to the ones of the convolutional layer. An optional second stage consists of a normalization operation, described in Section A.6.g. The third stage implements a non-linear elementwise operation, called an activation function, as described in Section A.6.f.

**Implementation:** In our network, several fully-connected layers are used after the convolutional ones to obtain the final outputs of the network, as illustrated schematically in Figure 13. For each fully-connected layer, the number of output scalar elements $k$ is a hyperparameter and is learned during hyperparameter search (see Section A.7). The weights $\mathbf{W}$ and $\boldsymbol{b}$ are optimized during training as part of the general learning process, following a gradient-based optimization, as we aim to approximate the experimentally obtained ground truth values by the output of the model.

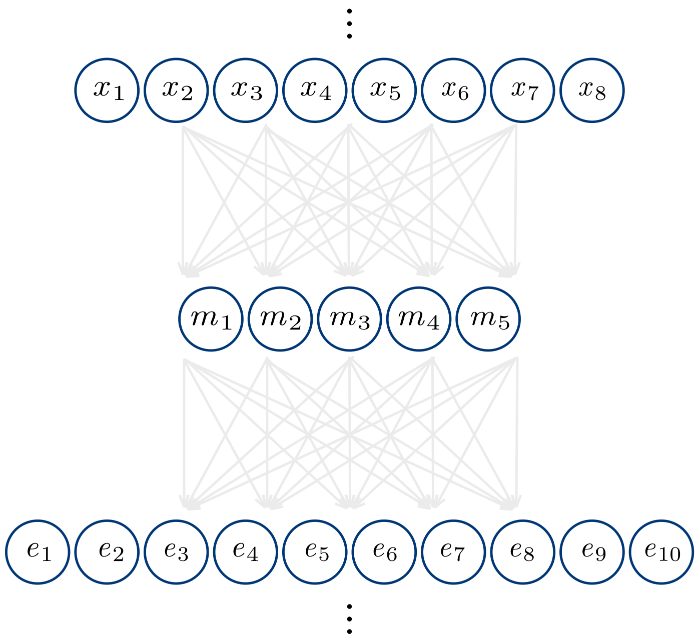

***Figure 13: Example of two Stacked Fully-Connected Layers.***

#### ***d. Skip connection***

**Skip connections** (He, 2016) have been introduced in order to avoid the problem of vanishing gradients for deep networks and to increase the speed of the training process. As its name suggests, a skip connection directly couples two layers' inputs that are at different depths in the network, skipping two or three intermediate layers, as illustrated in

Figure **14**. In the next paragraph, we will use the terms shallow and deep to differentiate between both layers' inputs.

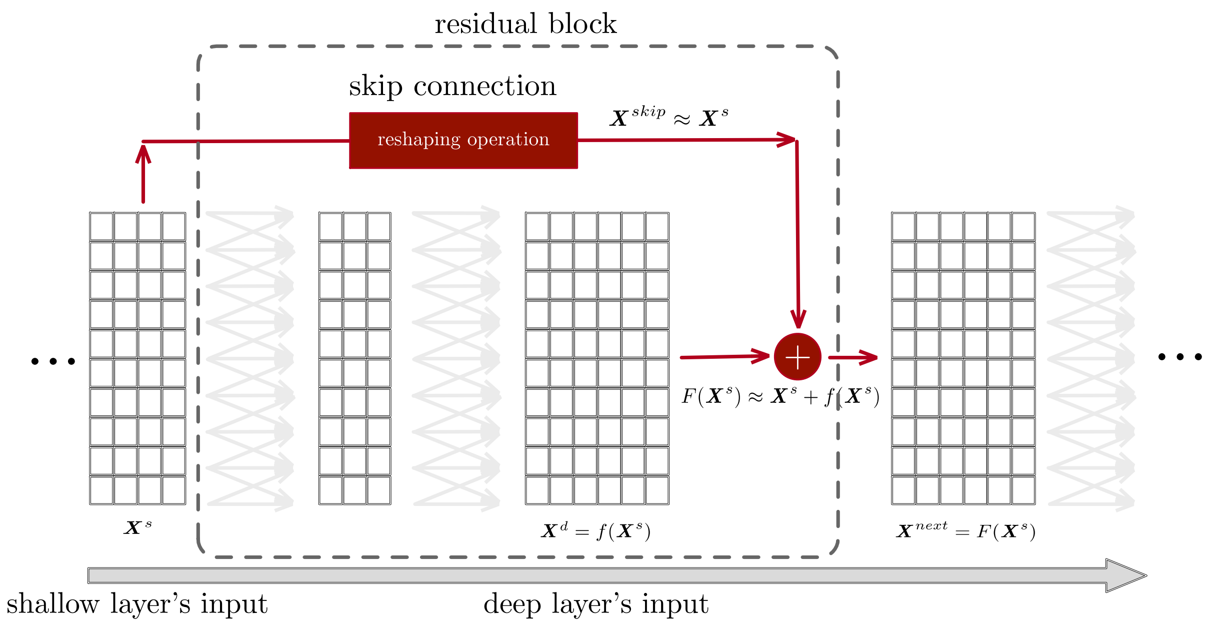

***Figure 14: Skip Connection.** The skip connection links together the shallow and deep layers' inputs, while both inputs are two convolutional layers apart in the main path of the neural network. In the skip connection, the shallow input array $X^s$ is reshaped to be of the same size as the deep input array $X^d$. The reshaping operation is needed in order to be able to add both arrays together (white plus shape). The result of the addition operation is a perturbation around the identity function $F(X^s) = X^{skip} + X^d \approx X^s + f(X^s)$. In some cases, learning $f(X^s)$ can be easier than learning $F(X^s)$ directly.* A residual block *describes all the operations from the shallow input $X^s$ to the results of the addition operation $F(X^s) = X^{next}$. Stacking several residual blocks sequentially allow the network to add together different levels of transformation of the input, from the identity mapping to a highly non-linear representation.*

In order to combine two distant input arrays, their shapes must be identical. Let us consider two input arrays, the shallowest of the two is $X^s$, of size $l \times c^s$, and the deepest one is $X^d$, of size $l \times c^d$. A 1d convolutional layer without activation function is used to **reshape** $X^s$ to obtain an array $X^{skip}$ of size $l \times c^d$. To this end, the 1d convolutional layer must contain as many filters as there are channels in the deep layer's input, i.e. $c^d$ filters in this example. In order to apply a reshaping transformation to the shallow input $X^s$ without modifying the input matrix (i.e. without padding), the $c^d$ filters are chosen to have a spatial dimension of length 1, their size is therefore $1 \times c^s$. As a consequence, the shape of the output of the skip connection $X^{skip}$ is $l \times c^d$, which is the same as the one of $X^d$. The two arrays are then added together element by element to form the input of the next layer $X^{next}$. The reshaping and addition operations are explained in detail in

Figure **15**. The block that is composed the intermediary convolutional layers, the skip-connection and the addition operation is called a **residual block**, as illustrated in

Figure **14**.

Let $i \in \{1, \dots, l\}$ be the position along the spatial dimension of the output and let $j \in \{1, \dots, c^d\}$ be the $j^{th}$ output channel number. Let $\mathbf{W}_{:,j}$ be the weight matrix and $b_j$ be the bias of the $j^{th}$ filter of the reshaping 1d convolutional layer. The output of the residual block can be expressed as a function of the deep and shallow input arrays:

$$X_{i,j}^{next} = X_{i,j}^{skip} + X_{i,j}^{d} = \sum_{v=1}^{c^s} X_{i,v}^{s} \mathrm{W}_{1,v,j} + b_j + X_{i,j}^{d} \quad (7)$$

**Implementation:** In our network, we use skip connections to link the inputs of two convolutional layers that are two layers apart. We chose to use those based on a preliminary hyperparameter search (see Section A.7). The number of residual blocks is fixed by hyperparameter search. The skip connections do not introduce hyperparameters as the number of filters and their sizes are fixed. However, the weights (**W** and ***b***) of the reshaping filters need to be learned during training.

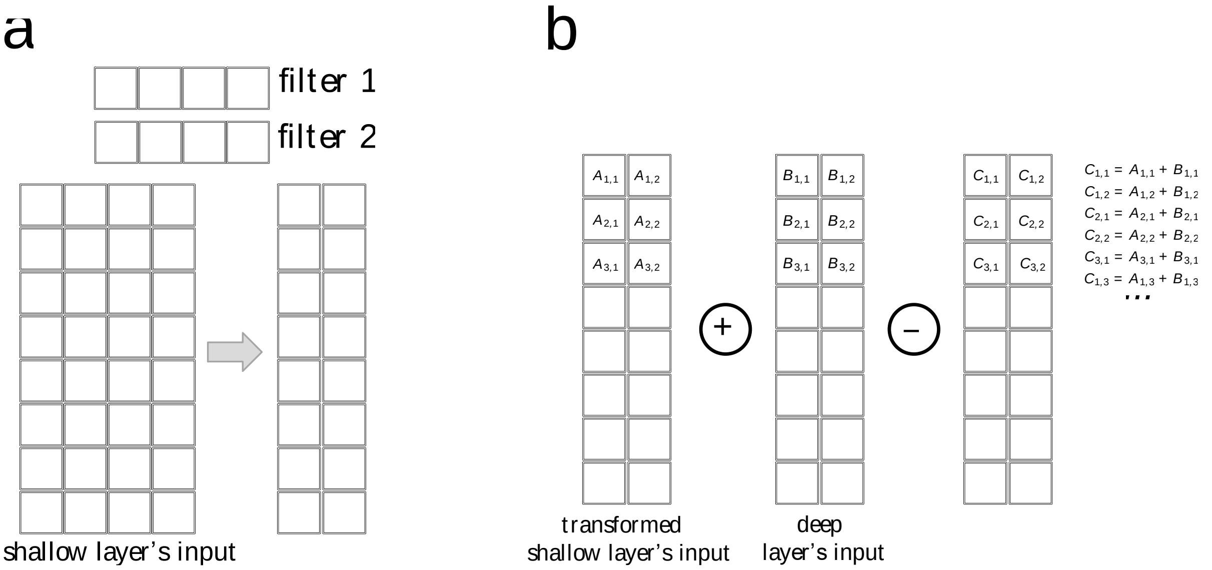

***Figure 15: Reshaping and Addition Operations of a Skip Connection. a**, Reshaping operation. Each filter has a spatial dimension of length 1 and as many channels as the shallow input, here 4. The output of the reshaping operation has as many rows as the input and as many columns as the number of filters. **b**, Addition operation. The transformed shallow input and the deep input are added together element by element.*

##### *e. Flattening step*

**Flattening** is another reshaping operation. For example, it transforms a 2D array, with $l$ rows and $c$ columns, into a 1D array, of $l \times c$ rows (and 1 column). They are useful to make the transition between a convolutional layer's output and a fully-connected layer's input, as the fully-connected layer is often implemented such that it only takes 1D inputs. Let $\boldsymbol{X}$ be the input array of the flattening step with $l$ rows and $c$ columns, $\boldsymbol{y}$ the output array, and $i \in \{1, \dots, l \times c\}$. A flattening step could be formulated as follows:

$$y_i = X_{\lfloor i/c \rfloor, i - \lfloor i/c \rfloor} \quad (8)$$

where $\lfloor i/c \rfloor$ means that we take the integer part of $i/c$. Another approach is to use a max (or average) **pooling** step, which would take the maximum (or average) over the spatial dimension for each channel. The computation of the max pooling of an array $\boldsymbol{X}$, of size $l \times c$, would result in $\boldsymbol{u}$, with $c$ rows, such that $u_j = max(X_{1,j}, X_{2,j}, \dots, X_{l,j})$ where $j \in \{1, \dots, c\}$. While pooling is commonly used in convolutional neural networks in protein-binding applications (Alipanahi, 2015), it presents a limitation in that the position information along the spatial dimension is lost. Therefore, pooling is less applicable to predicting RBS activity, for which the precise location of the Shine-Dalgarno and other motifs has crucial impact on RBS activity.

**Implementation:** When training the model on our application, we observed lower performances when using a pooling step instead of a flattened one between the last convolutional layer and the first fully-connected layer. As a consequence, we use a flattening step between the last convolutional layer and the first fully-connected layer. A flattening step has neither parameters nor hyperparameters.

##### *f. Activation function*

**Activation functions** (Goodfellow, 2016) are applied to every output scalar element of the second stage of a convolutional or a fully-connected layer in order to introduce non-linearities in the model. Typical activation functions are Tanh, Sigmoid, Softplus ($x \longrightarrow \log(1 + e^x)$) or Rectified Linear Unit (ReLU) functions (Nair, 2010; Sundararajan, 2017). The choice of such function can be done either by experience, by a manual exploration or during hyperparameter search. In this section, we focus on the ReLU function as it is often the default activation function and we use a variation of it in the model we develop. The **ReLU function** is defined as in Equation (9) and shown in Figure 16a:

$$f(x) = x^+ = max(x, 0) \quad (9)$$

$$i.e.\ f(x) = \begin{cases} 0 \ if \ x < 0 \\ x \ if \ x \geq 0 \end{cases}$$

The function output is the positive part of its argument. This function was introduced as its gradient is fast to compute and non-saturating. However, in some cases, the zero-slope part can also lead to a dying ReLU problem, as the gradient of the function is zero for negative input features. Therefore, the **Leaky ReLU activation function** was developed, which includes a small positive slope when the argument of the function is negative. The Leaky ReLU function, as illustrated in Figure 16b, is described by the following expression:

$$f(x) = max(x, 0.01 \times x) \quad (10)$$

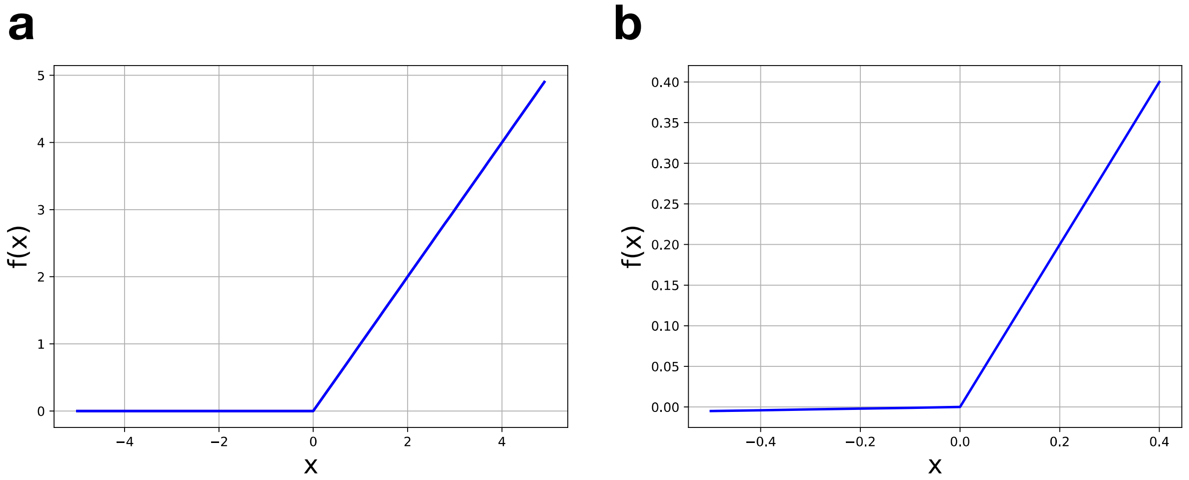

***Figure 16: Representations of Activation Functions. a**, ReLU function. **b**, Leaky ReLU function. The slope of the function for the negative part of its argument differs from the one of the ReLU function by being slightly positive and equal to* 0.01.

**Implementation:** The Leaky ReLU function is used as activation function for every layer of our model, except for the output layers (i.e. the two fully-connected layers that lead to the parameters $\alpha$ and $\beta$ in the rightmost part of the workflow Figure 4). It was chosen as it showed an improved performance on preliminary experiments. It is represented by light blue bars on the model on the right of the workflow Section A.4. For the output layers, we chose a Softplus activation function as the outputs need to be positive values (see Section A.9.a)

##### *g. Batch normalization*

**Batch normalization** (Ioffe, 2015) shifts and scales the output features of the first stage of a layer before they are used as input of the next stage. Scaling these intermediate output features allows to accelerate training and potentially get better results. As we will see in Section A.8.a, a model input (and therefore a layer input) is in fact the concatenation of several input samples, belonging to batch $\mathcal{B}$ of $n_{\mathcal{B}}$ samples $[\mathbf{X}_{:,i}]_{i=1}^{n_{\mathcal{B}}}$, where each $\mathbf{X}_{:,i}$ is of size $l \times c$. The batch normalization operation consists therefore of normalizing the elements of the output of the first stage of the layer across transformed samples of the batch $\mathcal{B}$. Batch normalization differs slightly between fully-connected layers and convolutional ones. In the case of a fully-connected layer, each element is normalized separately. In the case of a convolutional layer, the elements in the same channel are normalized the same way, but channels are normalized separately.

For a convolutional layer, the batch normalization pseudo-code can be described as follows:

Input: $\mathbf{X}$ a layer's output (3D tensor of size $l \times c \times n_{\mathcal{B}}$), $\varepsilon \ll 1$
Output: $\mathbf{Y}$ the batch normalization operation output (3D tensor of size $l \times c \times n_{\mathcal{B}}$)

1 Compute the mean of the batch along the sample and spatial dimensions: $\boldsymbol{m}_{\mathcal{B}} = \frac{1}{n_{\mathcal{B}} \times l} \sum_{i=1}^{n_{\mathcal{B}}} \sum_{j=1}^{l} \mathbf{X}_{j,:,i}$ *# $\boldsymbol{m}_{\mathcal{B}}$ is a vector of size $c$.*

2 Compute the variance of the batch: $\boldsymbol{\sigma}_{\mathcal{B}}^2 = \frac{1}{n_{\mathcal{B}} \times l} \sum_{i=1}^{n_{\mathcal{B}}} \sum_{j=1}^{l} (\mathbf{X}_{j,:,i} - \boldsymbol{m}_{\mathcal{B}})^2$ *# $\boldsymbol{\sigma}_{\mathcal{B}}$ is a vector of size $c$.*[2]

3 Normalize the output features: $\forall\, i \in \{1, \dots, n_{\mathcal{B}}\}\ \forall\, j \in \{1, \dots, l\}\quad \widehat{\mathbf{X}}_{j,:,i} = \frac{\mathbf{X}_{j,:,i} - \boldsymbol{m}_{\mathcal{B}}}{\sqrt{\boldsymbol{\sigma}_{\mathcal{B}}^2 + \varepsilon}}$ *# different scalar elements in a channel are normalized the same way.*

4 Scale and shift the normalized output: $\forall\, i \in \{1, \dots, n_{\mathcal{B}}\}\ \forall\, j \in \{1, \dots, l\}\ \mathbf{Y}_{j,:,i} = \boldsymbol{\gamma} \odot \widehat{\mathbf{X}}_{j,:,i} + \boldsymbol{\delta}$

***Algorithm 1: Batch Normalization Pseudo-Code for a Convolutional Layer.***

For a fully-connected layer, the batch-normalization pseudo-code can be written as follows:

Input: $\mathbf{X}$ a layer's output (3D tensor of size $l \times c \times n_{\mathcal{B}}$), $\varepsilon \ll 1$
Output: $\mathbf{Y}$ the batch normalization operation output (3D tensor of size $l \times c \times n_{\mathcal{B}}$)

1 Compute the mean of the batch along the sample and spatial dimensions: $\boldsymbol{M}_{\mathcal{B}} = \frac{1}{n_{\mathcal{B}}} \sum_{i=1}^{n_{\mathcal{B}}} \mathbf{X}_{:,:,i}$ *# $\boldsymbol{M}_{\mathcal{B}}$ is a 2D matrix of size $l \times c$.*

2 Compute the variance of the batch: $\boldsymbol{\Sigma}_{\mathcal{B}}^2 = \frac{1}{n_{\mathcal{B}}} \sum_{i=1}^{n_{\mathcal{B}}} (\mathbf{X}_{:,:,i} - \boldsymbol{M}_{\mathcal{B}})^2$ *# $\boldsymbol{\Sigma}_{\mathcal{B}}$ is a 2D matrix of size $l \times c$.*[3]

3 Normalize the output features: $\forall\, i \in \{1, \dots, n_{\mathcal{B}}\}\ \widehat{\mathbf{X}}_{:,:,i} = \frac{\mathbf{X}_{:,:,i} - \boldsymbol{M}_{\mathcal{B}}}{\sqrt{\boldsymbol{\Sigma}_{\mathcal{B}}^2 + \varepsilon}}$ *# all scalar elements are normalized differently.*

4 Scale and shift the normalized output: $\forall\, i \in \{1, \dots, n_{\mathcal{B}}\}\ \mathbf{Y}_{:,:,i} = \boldsymbol{\Gamma} \odot \widehat{\mathbf{X}}_{:,:,i} + \boldsymbol{\Delta}$

***Algorithm 2: Batch Normalization Pseudo-Code for a Fully-Connected Layer.***

In both algorithms (Algorithm 1 and Algorithm 2), $\mathbf{Y}$ is the normalized input of the third stage of the layer. In the last step of the batch normalization process, $\boldsymbol{\gamma}$ and $\boldsymbol{\delta}$ are vectors of parameters of size $c$ for convolutional layers and, $\boldsymbol{\Gamma}$ and $\boldsymbol{\Delta}$ are matrices of parameters of size $l \times c$ for fully-connected layers. These parameters are learned during training and render the network more flexible, without requiring to change all the weights of the next layer.

**Implementation:** Batch normalization is used at every layer of our model, between the linear operation (convolution or fully-connected operation) and the activation function (Leaky ReLU function). Preliminary experiments showed that it improved the performance of the model. It is represented by dark blue bars on the model on the rightmost part of the workflow Section A.4.

[2] $\boldsymbol{\sigma}_{\mathcal{B}}^2$ represents the elementwise square of $\boldsymbol{\sigma}_{\mathcal{B}}$.
[3] $\boldsymbol{\Sigma}_{\mathcal{B}}^2$ represents the elementwise square of $\boldsymbol{\Sigma}_{\mathcal{B}}$.

##### h. *Residual convolutional neural network*

***Implementation:*** *As represented in the rightmost box of*

**Figure 4**, the model we are using is composed of three residual blocks, each of them containing two convolutional layers. The output of the last residual block is then flattened and serves as input of two sets of two fully-connected layers, which yield two parameters $\alpha$ and $\beta$ (see Section A.9.b for the meaning of $\alpha$ and $\beta$). The depth of our model is therefore of eight layers, as there are eight layers between the input and each parameter. In addition, each layer of our model, except for the output layers (i.e. the ones that have as output $\alpha$ or $\beta$), is composed of three stages, including a batch normalization operation and a Leaky ReLU activation function. Models that present residual blocks and convolutional layers are often called **residual convolutional neural network** or ResNet. Section A.7 describes how the model architecture was chosen.

#### 7. Hyperparameter search

As explained in the glossary, **hyperparameter search** consists of a) training several models, distinct by their set of values of hyperparameters, on the training set, b) estimating a model selection criterion on the validation set for each model and c) selecting the best collection of hyperparameters. The latter, i.e. the best collection of hyperparameters, is determined based on the model selection criterion, which is often a function of the discrepancy between the model output and the ground truth target values of the samples that belong to the validation set. The lower (or the upper) this model selection criterion is, the better the collection of hyperparameters. In practice, the **automatic** hyperparameter search is often preceded in deep learning by a preliminary analysis to decide **manually** which hyperparameters will take part in the search, which intervals of values will be inspected (or which types of functions for hyperparameters that not scalars) and which hyperparameters will be fixed upstream. The preliminary analysis is also done on the validation set to avoid overfitting. Once the group of hyperparameters that are going to be analyzed automatically and their potential values are chosen, the space defined by the potential values taken by the hyperparameters can be explored by different means. A popular type of hyperparameter search is **grid search**, which probes *all* possible combinations of values of hyperparameters, which scales exponentially with the number of hyperparameters. A less exhaustive version is **random search**, which instead explores random combinations of values of hyperparameters. When the number of hyperparameters and potential values is large, grid search is too resource intensive while leading to a negligible gain in performance compared to random search. Both searches are in general done using parallel computing, where each model is trained in parallel on a different GPU (for deep learning).

**Implementation:** In order to choose the **hyperparameters**, we combined two approaches. First, we performed a preliminary analysis to fix some hyperparameters, such as the batch size and the activation function type. Second, we ran an extensive **random search**. It consisted of exploring 150 random combinations of hyperparameters. In our model, the hyperparameters can be classified in two categories, architecture parameters and optimization parameters. The hyperparameters listed below were the ones used to tune the ResNet model presented in the main manuscript.

##### *a. Architecture hyperparameters*

**Filter length of a convolutional layer**: The filter length is the length $r$ of the filter introduced in the Section A.6.b. The filter length defines the **receptive field**, which is the region of the input space that a particular convolutional layer output element is looking at, or a specific sequence motif length in the case of the input of the first convolutional layer (RNA sequences).

**Number of residual blocks**: The number of residual blocks is the number of pairs of convolutional layers, whose input and output are connected by a skip-connection.

**Number of filters of a convolutional layer**: The number of filters per convolutional layer corresponds to the variable $k$ defined in the Section A.6.c, which also matches the number of channels of the output of the layer.

**Number of output elements of a fully-connected layer**: The number of output elements of a fully-connected layer is equivalent to the number of matrix operations produced per fully-connected layer, i.e. $k$ as defined in the Section A.6.c.

**Activation function type**: The activation function type was chosen to be the Leaky ReLU function after a preliminary analysis, for all layers except for the last two fully-connected layers, as defined in Section A.6.f.

##### *b. Optimization hyperparameters*

**Learning rate**: The learning rate is a hyperparameter that determines how fast the weights of the model change as described in Section A.8.c. The incremental change of the weights at each iteration is proportional to the learning rate and to the gradient.

**Weight decay regularization constant**: The weight decay adds a squared norm regularization to the loss function. The objective of this penalty term is to bring the weights to small values. Its associated regularization constant $\lambda$ is a hyperparameter and settles the strength of the regularization term, which determines the pressure set on the weights to tend to zero.

**Batch size**: The batch size is the number of samples in each batch that are processed at the same time in the forward pass, as introduced in Section A.8.a.

#### 8. Fundamentals of the training process

This part of the appendix aims at describing briefly the training process of supervised deep learning models (Goodfellow, 2016). We will study how the data is passed as input to the model, how the information flows from the input to the output and back, and a standard optimization algorithm that can be used to minimize the cost function.

##### *a. Batches and epochs*

Deep learning models most often benefit from training on a small number of samples at a time instead of the full training set. The main reason is that the training dataset can be quite large ($> 10^6$ sample-target pairs) and that estimating the gradients when using a maximum of samples is more time consuming, while giving only marginally better mean estimates than training the model with a smaller sample size. Therefore, the training is done **batch** by **batch**, meaning that a predefined number of input samples (RNA sequences for example) are randomly picked and are given to the neural network at each iteration. The algorithm then optimizes a stochastic cost function, whose stochasticity comes from the one of the batches composition, and fixes temporarily the weights, before the next batch $\mathcal{B}$ is processed. Batches of samples can alternatively be called **minibatches** in the literature. In general, minibatch is used to design the pool of samples and batch is used when describing the size of the pool or the normalization operation. We will keep the term **batch** in the next paragraphs of the appendix.

Most often, batches contain non-overlapping samples and are processed one-by-one until the entire training set has been used. An **epoch** then coincides with the entire processed dataset. As deep learning is based on an iterative mechanism, it is often necessary to train on more than one epoch to obtain the best performance. The number of epochs is fixed by **early stopping**, which signifies that the training phase of the algorithm stops when the model selection criterion does not get better during a predefined number of consecutive epochs.

##### *b. Forward and backward passes*

In a nutshell, the weights of the model are initialized to small random values, following empirical rules (Glorot, 2010). During the **forward pass**, the information flows from the input **X**, through the functions defining the layers, to the model output $\boldsymbol{y}$. Then, the **cost function**, which roughly evaluates how far the estimated model output $\boldsymbol{y}$ is from its ground truth value, is calculated. In our case, the input **X** are one-hot encoded RNA sequences, sequenced with NGS, and the ground truth values are the normalized integrals of flipping profiles, experimentally measured. Next, the **back-propagation algorithm** allows the information to flow **backward**, from the output to the input, by computing the **gradient** (i.e. multi-dimensional derivative) of the cost function with respect to the weights of the model. Finally, the **optimization algorithm** (such as the stochastic gradient descent or the Adam optimization algorithm) exploits the estimation of the gradient to update the weights of the model such that the cost function decreases. As a consequence, the estimated output becomes closer to the ground truth values at each iteration. As seen above, the entire algorithm proceeds on batches. More detail about the training process can be found in (Goodfellow, 2016).

##### *c. Optimization algorithm*

The optimization algorithm is a key component of the training process and allows to update iteratively the weights of the model to decrease the cost function. Two common optimization algorithms are the **stochastic gradient descent** and the **adaptive moment estimation (Adam)** algorithms.

**Implementation:** We use the Adam optimization algorithm in our model as it worked best in some preliminary analyses. Therefore, we focus on this algorithm in this section.

Let $L(\boldsymbol{X}, y;\ \mathbf{W})$ be a **loss function**, which is a function of the difference between the model output and the ground truth value for a unique sample, with here, $\boldsymbol{X}$ an input sample, $y$ the ground truth target value and $\mathbf{W}$ the parameters (weights) of the model (Section A.9 gives a detailed description of the loss function of the proposed model). For simplicity, we do not include the biases $\boldsymbol{b}$ in the loss function but the same optimization operations are applied to $\mathbf{W}$ and to $\boldsymbol{b}$. Importantly, $L$ is differentiable with respect to the parameters $\mathbf{W}$, therefore a gradient can be estimated. We are interested in minimizing the **cost function** $F(\mathbf{X}, \boldsymbol{y};\ \mathbf{W})$ with respect to $\mathbf{W}$ in an iterative way, the cost function $F$ being the addition of a) the **expected loss**, i.e. the average $\mathbb{E}_{P_{data}(\boldsymbol{X},y)}[L(\boldsymbol{X}, y;\ \mathbf{W})]$ over the training sample-target pairs that follow $P_{data}(\boldsymbol{X}, y)$, and b) a regularization term $r(\mathbf{W})$ that will be defined in Section A.9 and only depends on the weights $\mathbf{W}$. The cost function $F$ is also called the training criterion. As we are passing the data to the model batch by batch, at each iteration number $t$ a different batch $\mathcal{B}_t = [\mathbf{X}_{t_{:,i}}]_{i=1}^{n_{\mathcal{B}_t}}$ is given to the model and used to calculate the cost function $F(\mathbf{X}_t, \boldsymbol{y}_t;\ \mathbf{W}_\mathrm{t}) = \mathbb{E}_{(\boldsymbol{X},y)\in(\mathbf{X}_t,\boldsymbol{y}_t)}[L(\boldsymbol{X}, y;\ \mathbf{W}_\mathrm{t})] + r(\mathbf{W}_\mathrm{t})$. Therefore, the quantity to minimize is stochastic, as it depends on random batches that are passed to the model at each iteration. In general, other elements can introduce stochasticity in the cost function, however this is not the case in the model developed in this manuscript.

The **adaptive moment estimation (Adam)** algorithm (Kingma, 2015) is a first-order gradient-based optimization algorithm of stochastic objective functions. It has been empirically shown that Adam is efficient at minimizing a stochastic cost function that can present local minima. For each iteration number $t$, the Adam optimization algorithm computes the **moving averages** of the first and second moments of the gradient of $F$, which are then used to update the weights with a first-order gradient. A moving average, as in Equations (11) and (12) below, averages the variable of interest over several contiguous iterations, weighting down older instances. Intuitively, the evolution of the weights is smoothened and the cost function converges better to a flat minimum. To summarize, the Adam algorithm is iteratively changing the weights in order to find a good attainable local minimum to the stochastic cost function.

Let $t$ be an iteration step, $t + 1$ the next iteration step and $\varepsilon$ a very small positive real value. At $t$, we assume we know $\mathbf{W}_t$ the parameters that are being optimized, $\mathbf{G}_{t+1} = \nabla_{\mathbf{W}}(F(\mathbf{X}_t, \boldsymbol{y}_t;\ \mathbf{W}_t))$ the gradient of the function $F$ with respect to the parameters $\mathbf{W}$, $\mathbf{M}_t$ the moving average of the first moment of the gradient and $\mathbf{V}_t$ the moving average of the second moment of the gradient.

The updated parameters $\mathbf{W}_{t+1}$ will therefore be written as in (13):

$$\mathbf{M}_{t+1} = \beta_1 \mathbf{M}_t + (1 - \beta_1)\mathbf{G}_{t+1} \quad (11)$$

$$\mathbf{V}_{t+1} = \beta_2 \mathbf{V}_t + (1 - \beta_2)\mathbf{G}_{t+1}^2 \quad (12)$$

$$\mathbf{W}_{t+1} = \mathbf{W}_t - \alpha \frac{\frac{\mathbf{M}_{t+1}}{1 - \beta_1^t}}{\sqrt{\frac{\mathbf{V}_{t+1}}{1 - \beta_2^t} + \varepsilon}} \quad (13)$$

The weights at time $t + 1$, $\mathbf{W}_{t+1}$, are estimated in (13) as a function of the weights at time $t$ and the moments of the gradients. The steps (11), (12) and (13) are repeated until a stopping criterion (see Section A.8.a for a brief description of early stopping). At the end of the optimization, the final weights $\mathbf{W}^*$ are saved for prediction on the hold-out test set. The denominators in (13), $1 - \beta_.^t$, correct for the biases of the estimates $\mathbf{M}_t$ and $\mathbf{V}_t$. The idea implemented in the Adam algorithm is that the weights are updated along the dimensions with largest gradients $\mathbf{M}_{t+1}$, which are penalized if the corresponding variance $\mathbf{V}_{t+1}$ is large, as the direction of the update becomes uncertain. The parameters $\beta_1$, $\beta_2$ and $\varepsilon$ are often kept fixed, respectively equal to 0.9, 0.999 and $10^{-8}$. The parameter $\alpha$ is called the **learning rate** and is determined during the hyperparameter search.

#### 9. Cost function of the residual convolutional neural network model

As explained in Section A.8.c, a **loss function** is a function that maps a datapoint $(\boldsymbol{X}, y)$ to a real-valued quantity that represents the "cost" associated to the prediction made for this datapoint. Most often, the better the prediction the lower the associated cost. The **cost function** measures an expected loss, averaged over the datapoints $(\mathbf{X}, \boldsymbol{y})$ that belong to the whole training set (in practice to a batch, see Section A.8.c), to which a regularization term is often added. The cost function, which is the training criterion, is minimized with respect to its parameters (Goodfellow, 2016). The cost function chosen in our model is detailed in this section.

##### *a. Modeling of the experimentally measured values*

In our study, the ground truth values to be predicted by the model are the experimentally measured normalized integrals of the flipping profiles ($IFP_{0\text{-}480min}$) that quantify RBS activity. As described in the main manuscript, the $IFP_{0\text{-}480min}$ values are continuous and defined on the interval $[0,1]$. Several approaches can be used to model such an output. In the context of neural networks, the most straight-forward way would be to use an appropriate activation function for the single output of the last fully-connected layer. Such activation function can be a **sigmoid function** ($f: \mathbb{R} \longrightarrow [0,1]; x \longmapsto \frac{1}{1+e^{-x}}$) or a **soft-clipping function** ($f: \mathbb{R} \times \mathbb{R} \longrightarrow [0,1]; (x, v) \longmapsto \frac{1}{v}\log(\frac{1+e^{vx}}{1+e^{v(x-1)}})$, with $v$ a hyperparameter) and the model output would then be defined on [0,1], as the ground truth value. However, as explained in Section A.11, we aim to estimate the uncertainty of the prediction together with the $IFP_{0\text{-}480min}$ values, which is generally not feasible with a single output value, as it is not sufficient in itself to estimate the uncertainty of each output value. Therefore, we chose to model each $IFP_{0\text{-}480min}$ value as a random variable whose distribution allows to get an estimate for the uncertainty of the prediction as well, as described Section A.11.

Towards this aim, we looked for a distribution that satisfies the following desiderata: i) be supported on the interval [0,1], ii) be differentiable with respect to its parameters, iii) be flexible enough to model the mean and the standard deviation independently, iv) be flexible enough to take a large number of shapes, such as asymmetric, non-monotonic or skewed distributions and v) be parsimonious, i.e. depend on a small number of parameters. Commonly-used distributions that match these requirements are the **beta distribution**, the **logit-normal distribution** and the **truncated normal distribution**. We discarded the logit-normal distribution as i) its mean and variance do not have a closed-form expression and have to be approximated differently, for example with a quasi Monte Carlo estimator and ii) preliminary experiments showed that the model had difficulties in parametrizing output distributions whose predictive mean ought to be close to the extremes of its support, i.e. near 0 or near 1. We also observed this difficulty in modelling very weak or very strong RBS activities when using the truncated normal distribution, albeit to a lesser extent. By contrast, the beta distribution did not suffer from any of those problems, and performed consistently well throughout the entire [0,1] range.

##### *b. Introduction to the beta distribution*

**Implementation:** As explained above, the **beta distribution** was the distribution of choice in our final residual convolutional neural network.

The beta probability density function (pdf) is defined as follows:

$$f(x; \alpha, \beta) = \frac{x^{\alpha-1}(1-x)^{\beta-1}}{B(\alpha, \beta)} \quad (14)$$

where $\alpha$ and $\beta$ are the two shape parameters of the beta pdf and $B(\alpha, \beta) = \int_0^1 u^{\alpha-1}(1-u)^{\beta-1}du$ is the beta function, which is a normalization constant that ensures that the total probability is 1. Roughly, the beta distribution can be interpreted as a generalization of the Bernoulli distribution and the shape parameters can be seen as expected numbers of draws of the two classes when they are larger than one. For example, in our context, the normalized integral can be seen as representing a proportion of flipped discriminator reads if we were looking at only one time point. If the total number of flipped discriminator reads would be $\alpha$ and the total number of non-flipped discriminator reads would be $\beta$, then we could estimate the proportion of flipped reads to be equal to $\frac{\alpha}{\alpha+\beta}$, which is exactly the mean of $x \mapsto f(x;\ \alpha,\beta)$ for $x \in [0,1]$, as $\int_0^1 \frac{u^{\alpha-1}(1-u)^{\beta-1}}{B(\alpha,\beta)} u du = \frac{\alpha}{\alpha+\beta}$.

##### c. *Negative log-likelihood as loss function*

**Implementation:** In this manuscript, the **loss function** of each residual convolutional neural network is the **negative log-likelihood** (see below) of the beta pdf. The associated **cost function** is the loss averaged over sample-target pairs in the training set, to which we added a weight decay $L_2$ regularization term.

Let $\mathcal{B}$ be a batch with $n_{\mathcal{B}}$ samples, let $i \in \{1, \ldots, n_{\mathcal{B}}\}$ be a sample number in the batch $\mathcal{B}$, $\mathbf{X}_{:,i}$ a one-hot encoded genetic sequence, $y_i$ its ground truth target value and $\mathbf{W}$ the weights of the model (we ignore the biases for simplicity of notation). The **likelihood** of having the ground truth target value $y_i$ given $\mathbf{X}_{:,i}$ as input of the model is $p_{\mathbf{W}}(y_i|\mathbf{X}_{:,i})$, which depend on the parameters $\mathbf{W}$. Likewise, the **log-likelihood** is $\log\left(p_{\mathbf{W}}(y_i|\mathbf{X}_{:,i})\right)$ and the **negative log-likelihood** is the opposite $-\log\left(p_{\mathbf{W}}(y_i|\mathbf{X}_{:,i})\right)$. We do not write the dependence on the hyperparameters $\phi$ as we can assume that they have been already selected on the validation set. The cost function can therefore be written as follows:

$$F(\mathbf{X},\boldsymbol{y};\mathbf{W}) = \underbrace{-\frac{1}{n_{\mathcal{B}}}\sum_{i=1}^{n_{\mathcal{B}}}\log\left(p_{\mathbf{W}}(y_i|\mathbf{X}_{:,i})\right)}_{\text{(a) Averaged negative log-likelihood over the samples in batch } \mathcal{B}} + \overbrace{\lambda}^{\text{(c) Weight decay hyperparameter}} \underbrace{\| \mathbf{W} \|_2^2}_{\text{(b) L2-norm regularization}} \quad (15)$$

(a) is equal to the averaged negative log-likelihood over the samples in batch $\mathcal{B}$. (b) is the regularization term, which is the sum of the square of each weight. It is multiplied by (c), the weight decay hyperparameter $\lambda$. Minimizing the cost function amounts to: i) minimizing (a) which corresponds to maximizing the averaged log-likelihood and to ii) minimizing (b) which shrinks the weights in order to prevent the model from overfitting to the training data and to increase the ability of the model to generalize on an unseen test data. The idea behind minimizing (a) over a batch $\mathcal{B}$ with respect to the parameters $\mathbf{W}$ is to find the optimal parameters that would ensure that for all elements in the batch, $y_i$ is very likely to be the output of the model when $\mathbf{X}_{:,i}$ is the input. As we chose to model the target variable with a beta probability density function, the likelihood can be written as $p_{\mathbf{W}}(y_i|\mathbf{X}_{:,i}) =$

$\frac{y_i^{\alpha_{\mathbf{W}}(\mathbf{X}_{:,i})-1}(1-y_i)^{\beta_{\mathbf{W}}(\mathbf{X}_{:,i})-1}}{B(\alpha_{\mathbf{W}}(\mathbf{X}_{:,i}),\beta_{\mathbf{W}}(\mathbf{X}_{:,i}))}$. The functions $\alpha_{\mathbf{W}}$ and $\beta_{\mathbf{W}}$ in the likelihood are both neural network functions, whose weights $\mathbf{W}$ are learned during training. Furthermore, $\boldsymbol{X} \longmapsto (\alpha_{\mathbf{W}}(\boldsymbol{X}), \beta_{\mathbf{W}}(\boldsymbol{X}))$ defines a space of beta pdfs and establishes a correspondence between an input sample $\mathbf{X}_{:,i}$ and a beta distribution. As the space of beta pdfs is constrained by the parameters $\mathbf{W}$, it is in principle not possible for the model to find the beta distributions that would maximize $p_{\mathbf{W}}(y_i|\mathbf{X}_{:,i})$ for each sample $\mathbf{X}_{:,i}$. However, we can hope that the model finds a space of beta distributions that allows to make the output $y_i$ very probable given $\mathbf{X}_{:,i}$ for all $i$.

Making the beta probability density function explicit, Equation (16) becomes:

$$
\begin{aligned}
F(\mathbf{X}, \boldsymbol{y}; \mathbf{W}) &= -\frac{1}{n_{\mathcal{B}}}\sum_{i=1}^{n_{\mathcal{B}}} \log\left(p_{\mathbf{W}}(y_i|\mathbf{X}_{:,i})\right) + \lambda \parallel \mathbf{W} \parallel_2^2 \quad (16) \\
&= -\frac{1}{n_{\mathcal{B}}}\sum_{i=1}^{n_{\mathcal{B}}} \log\left(y_i^{\alpha_{\mathbf{W}}(\mathbf{X}_{:,i})-1} \times (1-y_i)^{\beta_{\mathbf{W}}(\mathbf{X}_{:,i})-1}\right) + \lambda \parallel \mathbf{W} \parallel_2^2 \\
&+ \frac{1}{n_{\mathcal{B}}}\sum_{i=1}^{n_{\mathcal{B}}} \log\left(B\left(\alpha_{\mathbf{W}}(\mathbf{X}_{:,i}), \beta_{\mathbf{W}}(\mathbf{X}_{:,i})\right)\right) \\
&= -\frac{1}{n_{\mathcal{B}}}\sum_{i=1}^{n_{\mathcal{B}}} \left[\log\left(y_i^{\alpha_{\mathbf{W}}(\mathbf{X}_{:,i})-1}\right) + \log\left((1-y_i)^{\beta_{\mathbf{W}}(\mathbf{X}_{:,i})-1}\right)\right] + \lambda \parallel \mathbf{W} \parallel_2^2 \\
&+ \frac{1}{n_{\mathcal{B}}}\sum_{i=1}^{n_{\mathcal{B}}} \log\left(B\left(\alpha_{\mathbf{W}}(\mathbf{X}_{:,i}), \beta_{\mathbf{W}}(\mathbf{X}_{:,i})\right)\right) \\
&= -\frac{1}{n_{\mathcal{B}}}\sum_{i=1}^{n_{\mathcal{B}}} [(\alpha_{\mathbf{W}}(\mathbf{X}_{:,i})-1) \times \log(y_i) + (\beta_{\mathbf{W}}(\mathbf{X}_{:,i})-1) \times \log(1-y_i)] + \lambda \parallel \mathbf{W} \parallel_2^2 \\
&+ \frac{1}{n_{\mathcal{B}}}\sum_{i=1}^{n_{\mathcal{B}}} \log\left(B\left(\alpha_{\mathbf{W}}(\boldsymbol{X}_{:,i}), \beta_{\mathbf{W}}(\boldsymbol{X}_{:,i})\right)\right)
\end{aligned}
$$

##### *d. Mean and variance of the beta distribution*

The **mean** and the **variance** of the beta distribution are functions of the shape parameters. Let $i \in \{1, \ldots, n_{\mathcal{B}}\}$ be a sample number in a batch $\mathcal{B}$, $\mathbf{X}_{:,i}$ a one-hot encoded genetic sequence, $y_i$ its ground truth target and $\mathbf{W}^*$ the weights of the model estimated after minimization of the cost function.

After optimization, the mean of the beta pdf for sample $\mathbf{X}_{:,i}$ can be expressed as follows:

$$\mu_{\mathbf{W}^*}(\mathbf{X}_{:,i}) = \frac{\alpha_{\mathbf{W}^*}(\mathbf{X}_{:,i})}{\beta_{\mathbf{W}^*}(\mathbf{X}_{:,i}) + \alpha_{\mathbf{W}^*}(\mathbf{X}_{:,i})} \quad (17)$$

The corresponding variance can be written as:

$$\sigma^2_{\mathbf{W}^*}(\mathbf{X}_{:,i}) = \frac{\alpha_{\mathbf{W}^*}(\mathbf{X}_{:,i})\beta_{\mathbf{W}^*}(\mathbf{X}_{:,i})}{\left(\beta_{\mathbf{W}^*}(\mathbf{X}_{:,i}) + \alpha_{\mathbf{W}^*}(\mathbf{X}_{:,i})\right)^2 \left(\beta_{\mathbf{W}^*}(\mathbf{X}_{:,i}) + \alpha_{\mathbf{W}^*}(\mathbf{X}_{:,i}) + 1\right)} \quad (18)$$

For each sample, it is therefore possible to access the predicted mean and variance of its beta distribution once we learn the shape parameters. The predicted mean $\mu_{\mathbf{W}^*}(\mathbf{X}_{:,i})$ of a sample $i$ is the predicted target value and is an estimation for the ground truth target $y_i$. The predicted variance $\sigma^2_{\mathbf{W}^*}(\mathbf{X}_{:,i})$ contributes to the predictive uncertainty, as we will see Section A.11.

#### 10. Ensemble of residual convolutional neural network models

##### *a. Introduction to the notion of ensemble*

An **ensemble** is a set of predictive models whose individual outputs are combined, either in a weighted or unweighted fashion, in order to produce the estimated target of the input of interest (Friedman, 2001). Generally, ensembles are used to improve predictive performance. First, by averaging the output of the models in an ensemble, the algorithm reduces the risk to pick the wrong predictive model. Another advantage is that ensembles augment the space of functions that can be represented in comparison to a single model for finite training set sizes. Finally, in the context of neural networks, ensembles allow to run a search from different initial conditions and batch compositions.

##### *b. Mixture of beta distributions*

**Implementation:** In the manuscript we consider an ensemble containing $n_{\mathcal{M}} = 10$ residual networks, with different random initialization weights, random batch compositions (Lakshminarayanan, 2017) and hyperparameters. For every sample, each individual model predicts a beta pdf. The ensemble combines the individual predicted beta pdfs into a **uniformly-weighted mixture of beta probability density functions**. The predicted target of the sample of interest is given by the mean of the uniformly-weighted mixture of beta pdfs, as explained below.

Let $m \in \{1, \dots, n_{\mathcal{M}}\}$ be a model number, $i \in \{1, \dots, n_{\mathcal{B}}\}$ be a sample number in a batch $\mathcal{B}$, $\mathbf{X}_{:,i}$ a one-hot encoded genetic sequence, $y_i$ its ground truth target, $\mathbf{W}^*_m$ the optimized weights of model $m$, $\mu_{\mathbf{W}^*_m}(\mathbf{X}_{:,i})$ the mean of the beta pdf obtained with model $m$ for input $\mathbf{X}_{:,i}$ and $\sigma^2_{\mathbf{W}^*_m}(\mathbf{X}_{:,i})$ the variance of the beta pdf obtained with model $m$ for input $\mathbf{X}_{:,i}$. The output of the ensemble for each sample is a uniformly-weighted mixture of beta pdfs whose **mixture density** is as below:

$$p_{mixt}(y_i|\mathbf{X}_{:,i}) = \frac{1}{n_{\mathcal{M}}} \sum_{m=1}^{n_{\mathcal{M}}} p_{\mathbf{W}^*_m}(y_i|\mathbf{X}_{:,i}) = \frac{1}{n_{\mathcal{M}}} \sum_{m=1}^{n_{\mathcal{M}}} \frac{y_i^{\alpha_{\mathbf{W}^*_m}(\mathbf{X}_{:,i})-1} \times (1-y_i)^{\beta_{\mathbf{W}^*_m}(\mathbf{X}_{:,i})-1}}{B(\alpha_{\mathbf{W}^*_m}(\mathbf{X}_{:,i}), \beta_{\mathbf{W}^*_m}(\mathbf{X}_{:,i}))} \quad (19)$$

The predicted mean and variance of the mixture would then write as:

$$\mu_{mixt}(\mathbf{X}_{:,i}) = \frac{1}{n_{\mathcal{M}}} \sum_{m=1}^{n_{\mathcal{M}}} \mu_{\mathbf{W}_m^*}(\mathbf{X}_{:,i}) \quad (20)$$

$$\sigma_{mixt}^2(\mathbf{X}_{:,i}) = \frac{1}{n_{\mathcal{M}}} \sum_{m=1}^{n_{\mathcal{M}}} (\sigma_{\mathbf{W}_m^*}^2(\mathbf{X}_{:,i}) + \mu_{\mathbf{W}_m^*}^2(\mathbf{X}_{:,i})) - \mu_{mixt}^2(\mathbf{X}_{:,i}) \quad (21)$$

The mixture mean and variance are functions of the means and variances of each model in the ensemble, and by definition, of the respective shape parameters. When using an ensemble, we make the hypothesis that the normalized integral of the flipping profile ($IFP_{0\text{-}480min}$) is a random variable that follows a mixture of beta distributions, and not a beta distribution as it would be the case for a single model. The mixture mean corresponds to the predicted target and the mixture variance contributes to the predictive uncertainty, as we will see in Section A.11.

#### 11. Uncertainty estimate

As described in this section, obtaining high-quality **predictive uncertainty** estimates was achieved by a) using a **proper scoring rule** as training criterion (see below), b) using an ensemble of models and c) predicting the standard deviations of the target values, in addition to the target values themselves.

##### *a. Predictive uncertainty*

**Predictive uncertainty** can result from different sources: the **approximation uncertainty**, the **aleatoric uncertainty**, the **epistemic uncertainty** and the **distributional uncertainty** (Tagasovska, 2019; Malinin, 2018). First, the approximation uncertainty describes errors made by a model that would be too simple to fit complex data, for example if a linear regression is used to model a polynomial curve. Second, the aleatoric uncertainty accounts for the stochasticity of the data. It comes from measurement errors or a noisy observation process that would not be reduced by capturing more data in the same experimental conditions. Third, the epistemic uncertainty, also referred as model uncertainty, captures the inability of the model to properly optimize its weights. It can be reduced by adding more data. Finally, the distributional uncertainty comes from input space regions of low density (i.e. lack of training datapoints at some regions of the input space) and could also be reduced by adding datapoints at these low-density regions.

In our study, we are neglecting the approximation uncertainty as we are using a deep neural network, that is supposed to be a universal approximator (i.e. can fit in principle any function). We are also ignoring the distributional uncertainty, assuming that the sample density is sufficient at any region of the input space.

#### b. *Proper scoring rule*

The first step towards getting valid uncertainty estimates is to choose a **proper scoring rule** as training criterion (Lakshminarayanan, 2017). A **scoring rule** is a function that assigns a numerical score to a predictive distribution $p_{\mathbf{W}^*}(y|\boldsymbol{X})$ (here the weights $\mathbf{W}^*$ are the optimized ones) and rewards better calibrated predictive distributions, i.e. that are closer to the ground-truth conditional distribution of the targets given the samples $q(y|\boldsymbol{X})$. Let $S(p_{\mathbf{W}^*}, (\boldsymbol{X}, y))$ be a scoring rule that evaluates how well the predictive distribution matches the ground-truth conditional distribution $q(y|\boldsymbol{X})$ relative to the event $y|\boldsymbol{X}$. The expected scoring rule can be written $S(p_{\mathbf{W}^*}, q) = \int p_{\mathbf{W}^*}(y|\boldsymbol{X}) q(y, \boldsymbol{X}) dy d\boldsymbol{X}$. Most importantly, $S(p_{\mathbf{W}^*}, q)$ is a **proper scoring rule** if for all $p_{\mathbf{W}^*}$, $S(p_{\mathbf{W}^*}, q) \leq S(q, q)$, with equality if and only if $p_{\mathbf{W}^*}(y|\boldsymbol{X}) = q(y|\boldsymbol{X})$ for all sample-target pairs $(\boldsymbol{X}, y)$ in the dataset $D$.

**Implementation:** As described above in Section A.9.c, for individual models, we chose the log-likelihood as proper scoring rule, or equivalently the negative log-likelihood as loss function. As we saw in the previous section, the ground-truth conditional distribution of the targets given the samples was modeled as a beta probability density function.

#### c. *Aleatoric uncertainty*

The **aleatoric uncertainty** refers to the intrinsic uncertainty of the data, which would remain if we repeated the same experiment in the same conditions several times. It can for example be directly linked to a noisy observation process that cannot be reduced or captured with more datapoints under the same experimental conditions. In order for a single model to express a tailored aleatoric uncertainty, it is possible to model the target as a random variable and predict the variance $\sigma_\Theta^2$ of the distribution of the target for every sequence input (cf. Section A.9.d), in addition to the mean. In practice, the variance of the predictive distribution $p_{\mathbf{W}^*}(y|\boldsymbol{X})$ is a measure of the aleatoric uncertainty at each datapoint $\boldsymbol{X}$, assuming the weights $\mathbf{W}^*$ correspond to the true (unknown) weights.

#### d. *Epistemic uncertainty*

The **epistemic uncertainty** or model uncertainty refers to sources of uncertainty that would be reduced if additional information were given, for example a larger sample size. An example of such uncertainty is that the mathematical model could neglect certain measurable effects. In practice, this uncertainty can be accounted for by taking into consideration the uncertainty of the optimized parameters $\mathbf{W}^*$ of the single model $p_{\mathbf{W}^*}(y|\boldsymbol{X})$. As a matter of fact, the parameters $\mathbf{W}^*$ of the predictive distribution $p_{\mathbf{W}^*}(y|\boldsymbol{X})$ are estimated and do not necessarily correspond to the unknown ground truth parameters. In order to capture the uncertainty of these parameters, it is possible to use an ensemble of models, i.e. to average the predictive distributions over several models that are either initialized differently, use different random batches or have different hyperparameters. If we were to consider one model to predict the target, it would be equivalent to assuming that the parameters $\boldsymbol{W}^*$ are equal to the ground truth ones and therefore we would not account for the model uncertainty in the variance of the predictive distribution. By contrast, if we could learn all possible models, we could marginalize (i.e. integrate over) the parameters $\mathbf{W}^*$ to estimate the ground truth conditional distribution, $q(y|\boldsymbol{X})$, as the average over all the predictive distributions corresponding to the infinitely many optimized models: $q(y|\boldsymbol{X}) = \int p_{\mathbf{W}^*}(y|\boldsymbol{X}) r(\mathbf{W}^*|D) d\mathbf{W}^*$ where $D$ is the dataset of interest and $r(\mathbf{W}^*|D)$ is the posterior probability of the parameters given the dataset. However, in practice this is unfeasible, and the ensembles are a mean to capture some uncertainty, by estimating the posterior probability of the parameters $\boldsymbol{W}^*$, $r(\mathbf{W}^*|D)$, as a sum of Dirac delta functions centered on the optimized parameters of the $n_{\mathcal{M}}$ models of the ensembles, such that the predictive distribution can be estimated by $p_{mixt}(y|\boldsymbol{X}) = \frac{1}{n_{\mathcal{M}}} \sum_{m=1}^{n_{\mathcal{M}}} \int p_{\mathbf{W}^*}(y|\boldsymbol{X}) \delta(\mathbf{W}^* - \mathbf{W}_{\mathrm{m}}^*) d\mathbf{W}^*$. Therefore, calculating the variance $\sigma_{mixt}^2$ of the predictive distribution $p_{mixt}(y|\boldsymbol{X})$ allows to capture some epistemic uncertainty, together with the aleatoric uncertainty (cf. Section A.10.b).

##### *e. Validation of the predictive uncertainty*

For any prediction model, it is common to evaluate the accuracy of the predicted targets with metrics such as MAE or RMSE. In a similar way, it is also key to be able to evaluate predicted variances, which serve as proxy for predictive uncertainty. As we do not know the ground truth variances, it is not possible to directly compare the predicted values to the ground truth values with common metrics. To this end, different approaches have been developed, one of them is building a **reliability diagram** (Lakshminarayanan, 2017). The reliability diagram aims at establishing whether the predicted uncertainty is **well-calibrated**. It displays the percentage of ground truth values in the test set that fall into the $\tau\%$-confidence interval of their predicted beta probability density functions, for any $\tau \in [0, 100]$. This number is then compared to the theoretical percentage of ground truth values that should fall in the $\tau\%$-confidence interval, which is exactly $\tau\%$. If these two percentages agree for any $\tau\%$-confidence interval, i.e. if the identity mapping holds, we say that the model is well-calibrated and the predictive uncertainty is meaningful. If the percentage of ground truth values in a given $\tau\%$-confidence interval is smaller than $\tau\%$, it means that the model is over-confident and tends to be certain about weak predictions. In the opposite case, if the number of ground truth values in a given $\tau\%$-confidence interval is larger than $\tau\%$, it means that the model is underconfident and that the variances tend to be too large.

**Implementation:** In order to build the reliability diagram (Main Fig.4f), we used the estimated variances $\sigma_{mixt}^2$ of the mixture of the beta probability density functions in order to calculate the boundaries of the $\tau\%$-confidence interval for each sample and each $\tau$.

#### 12. Interpretability in deep learning

##### *a. Essentials on deep learning mechanisms*

One way to understand deep learning is to consider linear models and ways to overcome their limitations by introducing non-linearities (Goodfellow, 2016). Linear models, such as linear regression, have the main advantage that they are efficient and their weights are directly interpretable. However, they cannot capture interactions between features. A way around it that resembles the deep learning approach is to apply a linear model to a **kernel transformation** $\phi(\boldsymbol{x};\Theta)$ of the input $\boldsymbol{x}$ instead of to $\boldsymbol{x}$ directly. This is an example of a simple deep learning model with one intermediate layer $\phi$. While the user imposes the function $\phi$, the optimization algorithm of the deep learning model learns the weights $\Theta$ in order to find what corresponds to a good **representation** of the data. The linear model that is then applied on top of the new representation potentially corresponds to the output layer of the deep learning model $\boldsymbol{y} = \phi(\boldsymbol{x};\Theta)^T\boldsymbol{w}$. Therefore, at the core of deep learning, is the idea that we can improve models by learning new complex intermediary features.

Depth is another key aspect to the deep learning strategy and carries out two advantages (Goodfellow, 2016; Bengio, 2013; LeCun L. a., 2015). The first one is that they promote **reusing or combining features**. This can be intuitively understood as the number of paths from the input to the output grows exponentially with depth, and therefore the power of composition of input elements as well. As high-level features are obtained by composing lower ones, the deep learning model builds multiple levels of representation, or in other words, learns a **hierarchy** of features. In general, this enhances predictive performance as theoretical results have shown that deep representations can efficiently help to learn certain families of functions. The second advantage of deep models is that they learn different levels of feature abstraction, the deeper the more potentially abstract the feature representation. Importantly, learning more or less abstract features is key to solving the **specificity-invariance** trade-off, which requires an adapted balance between learning features that are sensitive to tiny input details and learning features that describe a general concept. Additionally, these abstract, non-linear feature representations potentially yield a better predictive power and are at the heart of entire machine learning research fields.

##### *b. Integrated gradients attribution method*

**Integrated gradients** are a mean to **attribute** the predictions of a deep network to its input features. For example, when the input is a genomic sequence whose features are bases, we aim to estimate the contribution of each base at each position to the output (Sundararajan, 2017). The intention is to be able to understand better the input-output behavior of the neural network. For example, this could help understanding which parts of the sequence and which specific bases lead to strong or weak RBSs. This knowledge could be further exploited in order to guess which subsequences would need to be modified when designing sequences with a specific behavior.

Formally, let $f: \mathbb{R}^l \longrightarrow [0,1]$ be a function that defines a deep learning model and $\boldsymbol{x} = (x_1, \dots, x_l)$ an input vector. Given a baseline input $\boldsymbol{x}' = (x'_1, \dots, x'_l)$, an **attribution** of the prediction at input $\boldsymbol{x}$ is a vector $\boldsymbol{a}(\boldsymbol{x}, \boldsymbol{x}') = (a_1, \dots, a_l)$, where $a_i$ is the contribution of input feature $x_i$ to the output $f(\boldsymbol{x})$ relative to $f(\boldsymbol{x}')$. The **integrated gradients** term stands for the calculations used to estimate the attribution vector $\boldsymbol{a}$. Gradients of the output $f(\boldsymbol{x})$ with respect to a feature $x_i$, $\frac{\partial f(x)}{\partial x_i}$, are estimated along the straightline path from $\boldsymbol{x}'$ to $\boldsymbol{x}$, i.e. for all $\boldsymbol{v} = \alpha\boldsymbol{x} + (1-\alpha)\boldsymbol{x}'$ for $\alpha \in [0,1]$ . These gradients, along the path, with respect to $x_i$, are then cumulated into an integral.

$$a_i(\boldsymbol{x}, \boldsymbol{x}') = (x_i - x'_i) \times \int_{\alpha=0}^{1} \frac{\partial f(\alpha\boldsymbol{x} + (1-\alpha)\boldsymbol{x}')}{\partial x_i} d\alpha \quad (22)$$

A direct consequence is that the sum of the integrated gradients over all features $i$ is equal to the output difference with the input $\boldsymbol{x}$ and the baseline $\boldsymbol{x}'$.

$$\sum_{i=1}^{l} a_i(\boldsymbol{x}, \boldsymbol{x}') = f(\boldsymbol{x}) - f(\boldsymbol{x}') \quad (23)$$

We usually refer to $\boldsymbol{a}$ as the integrated gradients or the **attribution profile**. The input is not necessarily a vector but can be a 2D or 3D arrays for example.

**Implementation:** In our model, the input is a 2D array and we are interested in the attributions of each element, i.e. of each base at each position. We chose the baseline to be the an $(l, 4)$-array of 0s. Therefore, each integrated gradient term $a_i(\boldsymbol{x}, \boldsymbol{x}')$ can be interpreted as the contribution of the element (specific base at specific position) to the predicted target to be above or below the output target of the all-zeros array, which is approximately 0.32.

### B. Data analysis: complementary section to the Methods

#### 1. Additional details: correlation of Bxb1-mediated recombination with cellular Bxb1-sfGFP levels

Before calculating summary statistics of the fluorescence profiles, we preprocessed the fluorescence profiles as follows. The preprocessing step consisted of imputing the fluorescence values a) for time points of profiles that are missing or b) at time point 0 for profiles that showed a fluorescence at 0 higher than the one at the following time point (50 minutes). To do so we fitted each of the biological replicate curves with a generalized logistic function ($x \mapsto \frac{K}{(1+Q\exp(-Bx))^{1/v}}$) using the values available and imputed the missing values or the inflated values (at 0) with the value of the fitted function at this time point.

To compute the summary statistics, the intervals of interest for the flipping profiles are [0, 360], [0, 480] and [0, 720] (minutes). Similarly, the fluorescence profiles intervals of interest are [0, 225], [0, 290], [0, 360] and [0, 480] (minutes). Integral-based summary statistics for the flipping profiles are estimated using the trapezoidal rule. Slope-based representations for the fluorescence profiles are calculated by fitting a linear regression to the datapoints within the interval of interest (boundaries included) of the three biological replicates. The slope of the fit serves as slope-based representation and the standard deviation of the slope is used to estimate the deviation around the estimated slope. Once the representations for both profiles are estimated, fits were evaluated from representatives of the flipping profiles to those of the fluorescence profiles. To do so linear ($x \longmapsto Ax + B$), log-linear ($x \longmapsto A\log(x) + B$) and general logistic fits ($x \longmapsto A + \frac{(K-A)}{1 + Q\exp(-Bx)}$) were used. The leave-one-out cross-validation consisted of learning the free parameters of the fitted functions on all but one internal-standard RBSs (here 30=31-1 datapoints) and predicting the output for the last datapoint that was not used for fitting. This step was performed 31 times (one time per inner-standard RBSs). The coefficient of determination was then calculated between the 31 inner-standard RBS representations of the fluorescence profiles and the 31 predictions.

#### 2. Logistic fit of the flipping profiles

A majority of flipping profiles present moderate irregularities, such as measurements at later time points might yield lower values than earlier ones. As the flipping profiles represent cumulative distributions of the proportion of flipped discriminator reads, this type of irregularities should ideally not be present. However, it is hardly possible to perfectly control all factors of the experimental protocol and this creates irreducible random noise. In order to smooth out the noise, it is possible to fit the profiles using a generalized logistic function. We used the following generalized logistic fit as it is a non-decreasing function and shows the same S-shape as the one observed on many of the profiles:

$$f(t;\ A, D, E, t_0, v) = \frac{A}{(E\ +\ e^{-D(t-t_0)})^{\frac{1}{v}}} \quad (24)$$

The fit summarizes the relationship between the time and the proportion of flipped discriminators per RBS.

As the profiles are numerous and diverse, $> 10^5$ observations, it is not possible to fit automatically all the profiles with the default parameters as it leads to several errors and unfitted profiles. To this end, we implemented a preprocessing algorithm to find automatically better initialization parameters, fit all the profiles in parallel and evaluate the normalized fitted flipping profile integrals. The algorithm can be described as follows:

Input: $\boldsymbol{Y}$ profiles array, $t_{start} = 0$ minutes beginning of integration interval, $t_{end} = 480$ minutes end of integration interval, $\boldsymbol{R}$ total reads array (same sample ordering as $\boldsymbol{Y}$), $mrt = 20$ minimal read threshold, $b_{\mathcal{C}} = 10$ number of bins, $n_{\mathcal{K}} = 50$ number of clusters
Output: $\boldsymbol{I}$ normalized integrals array

Crop the flipping profiles $\boldsymbol{Y}$ outside of the integration interval $[t_{start}, t_{end}]$.
Select all the flipping profiles that verify the minimal read threshold in the integration interval $mrt$. Let $\mathcal{S}_{mrt}$ be the set of profiles to be fitted.
Remove from $\mathcal{S}_{mrt}$ the flipping profiles that are constant and equal to 0 or to 1.
Evaluate the integral values of every profile in $\mathcal{S}_{mrt}$ according to the trapezoidal rule.
Rank the samples according to their integral values and cluster them by bins $\mathcal{B}_i$ of size $1/b_{\mathcal{C}}$, the bins' boundaries are $[1/b_{\mathcal{C}} \times (i-1), 1/b_{\mathcal{C}} \times i]$, for $i = \{1, \dots, b_{\mathcal{C}}\}$.
**for** $b$ **from** 1 **to** $b_{\mathcal{C}}$:
  Cluster the profiles in bin $\mathcal{B}_b$ with k-means, with $n_{\mathcal{K}}$ clusters.
  Compute the centroid profile $\mathbf{C}_{.,b,k}$ in for each cluster $\mathcal{C}_{b,k}$, for $k = \{1, \dots, n_{\mathcal{K}}\}$.
  **for** $k$ **from** 1 **to** $n_{\mathcal{K}}$:
    Fit the centroid profile $\mathbf{C}_{.,b,k}$. Save the parameters of the fit $\mathbf{P}^{init}_{.,b,k}$.
  **for** $k$ **from** 1 **to** $n_{\mathcal{K}}$: *# Step done in parallel on 32 CPU cores*
    Initialize the fitting functions for the cluster $\mathcal{C}_{b,k}$ with $\mathbf{P}^{init}_{.,b,k}$.
    Fit all profiles in $\mathcal{C}_{b,k}$ and save the new parameters $\mathbf{P}_{:,b,k}$.
Fit manually the profiles that are constant and equal to 0 or 1 (trivial fit).
Compute the normalized integrals of the fitted flipping profiles $\boldsymbol{I}$. *# Step done in parallel on 32 CPU cores*

***Algorithm 3: Preprocessing Pseudo-code to fit the Profiles and Compute the Normalized Integral of the Flipping Profiles IFP$_{0\text{-}480min}$.***

#### 3. Additional details: RBS library design

Degenerate RBSs are constrained to the IUPAC notation. As a reminder, the IUPAC notation is a standard DNA (or RNA) representation where each character encodes a set of nucleotides. The single nucleotides {A, C, G, T, U} are represented as such, {W, S, M, K, R, Y} represent pairs of nucleotides, {B, D, H, V} triplets of nucleotides and N any nucleotide. To conduct this RBS library design, we first calculated the normalized integrals of the flipping profiles using the trapezoidal rule between the time points 0 and 360 minutes (IFP$_{trz}$). The resulting normalized integral values were partitioned in ten bins of size 0.1 from 0 to 1. The position probability matrices of the two highest bins were computed. Then, the libraries High1 and High2 were designed by finding the IUPAC sequences that are closest in terms of minimal mean-squared error to the PPMs of each of the two highest bins, respectively. Finally, the creation of the library High3 involved the coupling of a prediction model and a genetic algorithm. To this end, we first investigated whether a deep learning-based model trained on the proof-of-principle dataset could achieve non-trivial predictive performance, despite the relatively small sample size, in order to meaningfully guide library design. For this purpose, a convolutional neural network (CNN) composed of one convolutional layer and two fully-connected layers, the last one's output being a scalar value, was used. The predictive performance of such a model was explored by means of 5-fold cross-validation on the proof-of-concept dataset, such that for each fold, 70% of the data is used for model fitting, 10% for model selection and the remainder 20% a held-out test set. The hyperparameters were selected with grid search on each validation set. As shown in Fig. S7, despite being outperformed by SAPIENs when trained on the final dataset, this smaller model achieves sufficient predictive power ($R^2$=0.644, MAE=0.055) to guide library design. Therefore, we selected a final model by an additional 5-fold cross-validation, in this case using 80% of the data in each fold for model fitting and the remainder 20% for model selection. The best set of hyperparameters found in this manner was composed of a filter size of 5 and a number of filters of 128 for the first convolutional layer, 16 output elements for the first fully-connected layer, a weight decay of 0.005, the learning rate equal to 0.0001 and the batch size set to 512. Once this final model is selected and fitted, the RBS sequences from the three highest bins were randomly mutated for 200 iterations, with 1 or 2 mutations, keeping only the mutations that led to an increase in $IFP_{trz}$ as measured by the predicted values of the CNN. The PPM of the pool of mutated sequences was calculated, a sub-sample of 20,000 sequences was randomly generated from this PPM and the predicted $IFP_{trz}$ was computed for each generated sequence of the sub-sample using the trained CNN. Starting with a random degenerate RBS sequence, we mutated it iteratively one position at a time (random ordering of the positions), and kept the IUPAC nucleotide at the corresponding position that led to the smallest Kolmogorov-Smirnov distance between the predicted $IFP_{trz}$ distribution of the sub-sample and the predicted $IFP_{trz}$ distribution of the 1000 sequences generated from the new degenerate RBS sequence. This iterative process was continued until the relative decrease in KS distance was less than $\epsilon = 10^{-3}$ for three consecutive iterations.

#### 4. Additional details: cross biological replicates analysis

We evaluated the ability of SAPIENs to generalize across biological replicates, that is, we would like to assess whether a model trained on measurements from one batch can accurately predict targets whose ground-truth values were measured on a different batch. To this end, we first collected data from three distinct biological replicates (batches) and normalized the targets of the two additional biological replicates to the reference replicate (see Methods). Next, each replicate dataset was randomly split into (stratified) training, validation and test subsets as previously done (see Methods). Finally, we removed any sequences from the test sets which did not pass the quality control criteria for all three replicates, allowing our results to be directly comparable across replicates.

After these preprocessing steps, we trained three instances of SAPIENs independently on the training subset of each replicate, using the corresponding validation subset to select any hyperparameters. In practice, we ran the models for 150 epochs, used an early-stopping criterion on the validation set and performed random search among 150 sets of hyperparameters. For each of the three models, we evaluate its predictive performance on: 1) test labels measured in the same batch, to assess the within-replicate performance; 2) test labels measured in the other two batches after normalization, to assess cross-replicate performance after proper normalization.
